# Community reorganization without collapse in a warming world: Habitat-contingent, trait-mediated biodiversity change within a conserved multi-scale structure

**DOI:** 10.64898/2026.05.21.726918

**Authors:** Masami Fujiwara, Fernando Martinez-Andrade, Joshua Lerner, Mark Fisher, Rusty A. Feagin

## Abstract

Biodiversity change under sustained environmental pressure is increasingly understood as a process of long-term reorganization across spatial scales rather than uniform loss or gain. Estuarine ecosystems, which experience strong environmental gradients and frequent short-term extreme disturbances, provide an ideal context for evaluating how local (α), spatial (β), and regional (γ) diversity reorganize across habitat domains and functional guilds through time. Here, we examined four decades of fish and invertebrate community data from a coastwide, ecological monitoring program, spanning multiple estuarine systems along the Texas coast. We asked three main questions: whether habitat context modifies the direction of trait-mediated biodiversity responses across the full α–β–γ structure; whether spatial homogenization is detectable through dominance-weighted β-diversity metrics and how they relate to richness-based metrics; and whether the pace of reorganization has intensified over time. Functional traits related to dispersal capacity and thermal affinity structured the magnitude of biodiversity change within habitat domains, but habitat context determined its direction, the same guilds exhibited opposite diversity trajectories across gear-defined habitat domains despite broadly similar functional composition, demonstrating that trait-based forecasting frameworks will mispredict reorganization direction when habitat context is ignored. Spatial homogenization proceeded undetected by richness-based β-diversity metrics but was revealed through dominance-weighted measures, indicating that dominance-weighted homogenization operates through two distinct habitat-dependent mechanisms: convergence on shared warm-adapted dominant taxa in shallow assemblages, and convergence toward impoverished assemblages as dominant taxa declined in deeper habitats, both processes invisible to richness-based metrics. Despite a measurable intensification in the pace of local and regional diversity change after approximately 2000, spatial turnover among bays remained stable and α–β–γ scaling relationships were preserved across all diversity orders, demonstrating that rapid reorganization can proceed without destabilizing fundamental metacommunity structure. Across all findings, biodiversity change was persistent, directional, and strongly habitat dependent, following a chronic rather than episodic trajectory consistent with sustained environmental processes as the primary driver.

## 1. Introduction

Global environmental change is actively reshaping patterns of biodiversity worldwide. However, biodiversity responses often differ between local communities and broader regional assemblages, as environmental shifts may disadvantage some species while facilitating others depending on ecological scale and historical context. Long-term observations demonstrate that local assemblages may gain species, experience shifts in dominance structure, or undergo substantial compositional turnover, even as regional diversity remains stable or changes more gradually (McGill et al. 2015; Blowes et al. 2019, 2024). For example, increases in local richness may coincide with declining spatial differentiation among communities, while temporal redistribution of dominant taxa can alter ecosystem structure and function without pronounced changes in overall species counts (Magurran et al. 2019). Such patterns underscore that biodiversity change is inherently multivariate and scale dependent, and that analyses confined to a single diversity metric risk overlooking broader ecological dynamics. Interpreting biodiversity change therefore requires explicit integration of local (α), spatial turnover (β), and regional (γ) components within a unified analytical framework (Whittaker 1960, 1972). Consideration of α–β–γ diversity, therefore, provides a mechanistic basis for understanding biodiversity change as coordinated reorganization across time and space, rather than as isolated gains or losses at any single level.

One principal mechanism by which environmental drivers reorganize biodiversity is through changes in spatial differentiation among communities, captured by spatial β-diversity. Spatial β-diversity quantifies reorganization across space, rather than changes in local species counts alone, and thus provides insight into how communities converge or diverge through time. Increasing attention has been directed toward biotic homogenization, defined as the tendency for assemblages to become more similar over time. This process can arise through changes in β-diversity and can influence both local α- and regional γ-diversity in different ways, depending on the underlying mechanism (Olden and Rooney 2006). Homogenization may result from species loss and nestedness (rare species disappear, leaving communities with increasingly similar assemblages), or alternatively from convergence in the identity and dominance of common taxa, even when local richness remains stable or increases (Dornelas et al. 2014; Magurran and Henderson 2010). Conventional richness-based measures of β-diversity are poorly suited to distinguish these pathways, as they are disproportionately influenced by rare species and may obscure changes in dominant species that strongly affect ecosystem processes. Partitioning approaches have clarified the relative roles of turnover (replacement of species) and nestedness (Baselga 2010), yet accumulating evidence indicates that long-term community dynamics and stability are often governed by changes in dominant species (McGill et al. 2007). Dominance-weighted measures of β-diversity therefore provide a critical, though underutilized, perspective for detecting spatial homogenization driven by convergence in species abundances rather than by taxonomic losses or gains. An underexplored implication is that studies relying solely on richness-based β-diversity metrics may systematically fail to detect homogenization that is already underway through convergence in dominant taxa, producing false assurance of spatial stability even as communities functionally converge.

Spatial reorganization and dominance shifts are ultimately based on individual species responses to environmental change, through their functional traits. Traits related to dispersal ability, thermal affinity, and life-history strategy influence how species redistribute across space and reorganize within communities in response to changing environmental conditions (Lavorel and Garnier 2002; McGill et al. 2006). Consequently, taxonomic responses may appear idiosyncratic, even when the underlying ecological processes are highly predictable. Trait-based approaches provide a means of identifying these latent regularities, revealing consistent pathways of community reassembly driven by shared ecological strategies rather than by species identity (Winemiller and Rose 1992). In marine systems, thermal traits in particular constrain the rate and direction of species redistribution under climate warming, producing coherent spatial responses across diverse taxa (Pinsky et al. 2013; Sunday et al. 2012). However, existing frameworks assume that trait identity predicts the direction of biodiversity response, that warm-affinity or high-dispersal species will consistently gain regardless of local setting. Whether habitat context modifies or overrides this prediction across the full α–β–γ diversity structure has rarely been tested in long-term empirical systems, leaving open a fundamental question about the transferability of trait-based forecasting frameworks across habitat domains.

The relative importance of persistent environmental change versus episodic disturbance events on biodiversity reorganization also remains unclear. Chronic environmental change is associated with continuous pressure on populations and communities, producing gradual but directional change through time (Parmesan and Yohe 2003; IPCC 2021). In contrast, acute disturbances, such as storms and freezing events, are discrete and highly visible and often presumed to dominate ecological through their immediate impacts (Turner 2010; Smith 2011). However, despite their short-lived but highly visible effects, the long-term importance of such events relative to persistent environmental forcing remains unresolved. Many empirical studies examine either chronic trends or individual disturbance events in isolation, rarely evaluating both within the same analytical framework or using identical response variables and spatial units (Jentsch et al. 2007). Resolving this uncertainty requires analytical approaches that directly compare chronic and acute drivers across long time horizons using consistent spatial units and response variables.

Estuaries provide a particularly informative context for examining multiscale biodiversity reorganization because they combine strong chronic environmental gradients in water temperature, salinity, and dissolved oxygen with recurrent episodic disturbances, including extreme weather and temperature events (Day et al. 2012). Furthermore, these systems exhibit pronounced spatial heterogeneity that is tightly coupled through tidal exchange, freshwater inflow, and organismal movement, resulting in local communities that are simultaneously shaped by environmental filtering and dispersal (Sheaves et al. 2015). Such coupling makes estuaries well suited for analyses that explicitly integrate α-diversity, β-diversity, and γ-diversity, as changes at one scale can propagate across others. However, estuarine biodiversity has often been assessed using short-term or spatially limited datasets, constraining inference about long-term reorganization and the relative importance of chronic versus acute drivers (Elliott and Whitfield 2011; Whitfield et al. 2012). Long-term, spatially replicated estuarine monitoring therefore offers a powerful opportunity to disentangle the processes governing biodiversity reorganization across scales.

The Texas estuarine monitoring program (Martinez-Andrade 2018) represents one of the longest and most spatially extensive datasets available for coastal fish and invertebrate communities. Spanning more than four decades and encompassing replicated bay systems along a continuous subtropical coastline, this program combines consistent sampling protocols across habitats with broad spatial coverage and concurrent environmental monitoring, providing an unusual opportunity to evaluate biodiversity reorganization across scales under sustained environmental forcing. However, prior analyses have largely focused on single diversity components, compositional ordinations, or restricted subsets of taxa, leaving unresolved how local, spatial, and regional diversity components change in concert, whether trait-based predictions of reorganization direction hold across gear-defined habitat domains, and whether ongoing homogenization is detectable only through dominance-weighted rather than richness-based β-diversity metrics. Integrating α-, β-, and γ-diversity within a unified framework therefore provides a critical next step in identifying general mechanisms of biodiversity reorganization along the Texas coast.

Our overall objective is to identify the mechanisms underlying long-term biodiversity reorganization across habitat domains in a dynamic coastal ecosystem. Specifically, we ask: (1) whether habitat context modifies or overrides trait-based predictions of reorganization direction across the full α–β–γ diversity structure; (2) whether spatial homogenization is detectable through dominance-weighted β-diversity metrics and how they relate to richness-based metrics; and (3) whether the pace of reorganization has changed. Addressing these questions advances understanding of how communities reorganize under sustained change, with implications for biodiversity monitoring, trait-based forecasting, and metacommunity theory.

## 2. Methods

All analyses were performed in R (version 4.5.0; R Core Team 2024), with all associated scripts publicly available on GitHub (https://doi.org/10.5281/zenodo.20329852).

### 2.1 Study area and environmental context

The Texas coast comprises a series of semi-enclosed estuarine systems extending approximately 600 km along the coastline (Fig. 1). These estuaries exhibit strong spatial heterogeneity in geomorphology, freshwater inflow, salinity, and temperature regimes (Longley 1994), generating pronounced environmental gradients that influence community structure and population dynamics. They differ markedly in hydrological connectivity, watershed size, residence time, and nutrient supply. Northern systems, such as Sabine Lake and Galveston Bay, receive substantial riverine discharge, whereas southern systems, including the Laguna Madre, are hypersaline lagoons with minimal freshwater inflow (Withers et al. 2017; Bugica et al. 2020). Consequently, estuaries along the Texas coast span broad gradients in salinity, turbidity, temperature variability, and benthic habitat composition, establishing powerful environmental filters that spatially structure fish and invertebrate assemblages by selectively favoring taxa whose physiological tolerances align with local environmental gradients.

**Figure 1.**
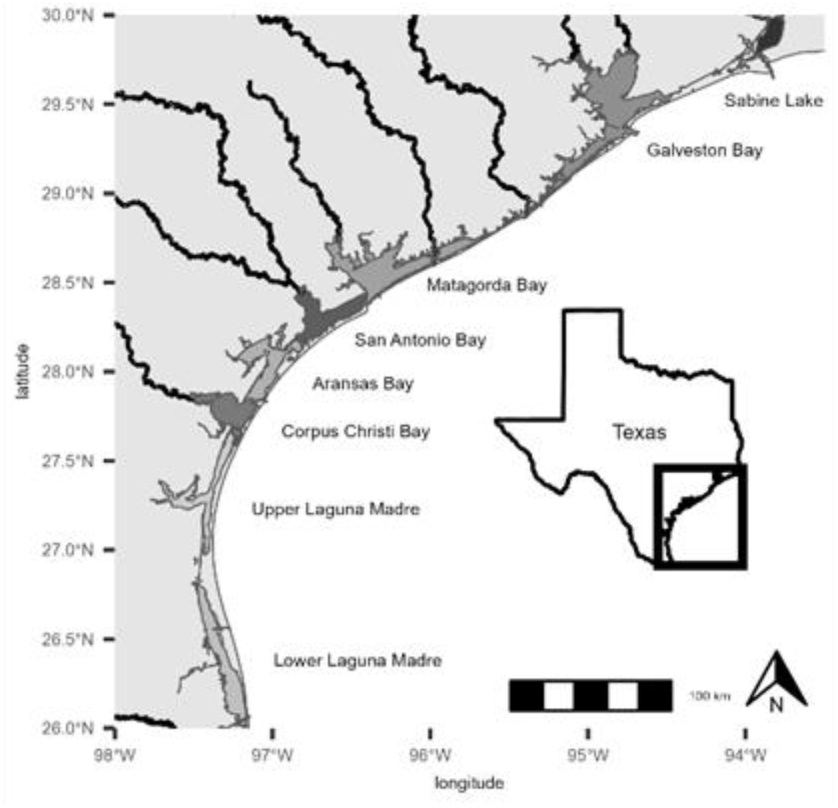
Study area and sampling design. Map of the major estuarine regions along the Texas coast, within which the Texas Parks and Wildlife Department sampling stations were located.

Environmental conditions along the Texas coast also vary strongly through time. Interannual temperature variability is influenced by large-scale climate modes such as the El Niño-Southern Oscillation, while episodic acute conditions, including winter freezes, summer heatwaves, and tropical cyclones, can produce rapid system-wide ecological responses (Patrick et al. 2020). These acute disturbances can alter salinity regimes (Du and Park 2019), increase turbidity (Poirrier et al. 2008), disrupt recruitment (Biggs et al. 2018), and cause short-term mortality across multiple taxa (Vincenzi et al. 2014), contributing to abrupt deviations from long-term community trajectories. In contrast, chronic environmental forcing operates gradually over longer time scales through sustained changes in background conditions.

Episodic disturbances are superimposed on a broader climatic trend of rising water temperatures, shifting thermal regimes, and intensifying precipitation extremes along the Texas coast (Tolan and Fisher 2009; Wang et al. 2023; Feagin et al. 2024). Long-term warming has been linked to poleward range expansions of warm-adapted species, contractions of temperate taxa, and evidence of progressive “tropicalization” in temperate and subtropical coastal ecosystems (Fujiwara et al. 2019; Vergés et al. 2019). Spanning this transition from temperate to subtropical conditions and encompassing estuaries with contrasting hydrology and habitat structure, the Texas coast serves as an ideal natural laboratory for examining long-term spatiotemporal variation in biodiversity and disentangling the relative influences of chronic environmental change and episodic disturbances on estuarine community dynamics.

### 2.2 Long-term monitoring program

Field sampling followed the procedures described in the *Coastal Fisheries Resource Monitoring Operations Manual* (Martinez-Andrade 2018), which specifies gear-specific deployment methods, fixed seasonal intervals, and standardized effort units by the Texas Parks and Wildlife Department (TPWD) Coastal Fisheries Division. The data were collected with three gears, Bag Seine (BS), Bay Trawl (BT), and Gill Net (GN). Each targeted a distinct component of the estuarine community and collectively they provided a comprehensive assessment of species occurring along the shoreline, in shallow subtidal habitats, and throughout the water column. The sampling followed the TPWD guidelines, which provided a consistent and standardized record of estuarine fish and invertebrate assemblages along the Texas coast. The monitoring program spanned the eight major estuarine systems: Sabine Lake (Sabine-Neches Estuary), Galveston Bay (Trinity-San Jacinto Estuary), Matagorda Bay (Colorado-Lavaca Estuary), San Antonio Bay (Guadalupe Estuary), Aransas Bay (Mission-Aransas Estuary), Corpus Christi Bay (Nueces Estuary), Upper Laguna Madre, and Lower Laguna Madre. These major bays/estuaries form the spatial framework for all subsequent analyses.

Bag seine sampling targeted shallow subtidal areas and was highly effective for nearshore resident and nursery species. Gillnet sampling targeted pelagic habitats and was highly effective for larger, mobile, and predatory species. Bay trawl sampling targeted benthic and near-bottom habitats and was highly effective for demersal fishes (e.g., flounder, croaker) and invertebrates living on or near the substrate (e.g., crabs, shrimp).

Bag seine and bay trawl surveys were each conducted approximately 20 times per month (January–December) within each major estuarine system, while gillnet surveys were conducted seasonally in spring and fall, with 45 samples collected per season in each major bay. The sampling intensity was largely consistent from 1986 through 2024, whereas sampling effort was much lower during the initial years of the monitoring program (1982–1985). During each survey, a measurement station was established to record environmental variables such as, water temperature, salinity, and dissolved oxygen were recorded using a YSI meter or equivalent instrument following standardized monitoring procedures.

The bag seines used in this study measured 18.3 m in length and 1.8 m in depth, with 19 mm mesh in the wings and 13 mm mesh in the bag. Deployment involved a fixed 15-m haul parallel to the shoreline, resulting in a consistent area sampled at each station. The bay trawl had a 5.7 m (18 ft 8 in) headrope and a 7.0 m (23 ft) footrope made of 38 mm stretched-mesh nylon. Sampling effort was standardized by towing the trawl for a fixed duration and distance at a constant vessel speed. The gillnets were 45.7 m (150 ft) passive entanglement nets composed of four stretched-mesh panels (76, 102, 127, and 152 mm). Sampling effort was standardized by a fixed soak-time. All captured individuals greater than 5 mm in length were identified to the lowest feasible taxonomic level and counted.

The selection of sampling locations followed a stratified random design. Each major estuarine system was subdivided into fixed 1-minute latitude x 1-minute longitude grids, a subset of which was randomly selected without replacement each month. Within each selected grid, one “gridlet,” defined as a twelve-by-twelve subdivision of the parent grid, was chosen at random. The gridlet was further partitioned into 50 shoreline segments of at least 15 m in length, and one segment was randomly chosen for sampling. This hierarchical spatial sampling scheme, combined with temporal dispersion throughout the month, ensured that sampling events were spatially and temporally independent, thereby minimizing pseudo-replication and preserving the statistical integrity of the long-term time series. Although the main part of the monitoring program began in 1982, the initial deployment year differed by gear and estuary. All available samples since the initial deployment were retained to maximize temporal coverage.

### 2.3 Biological data processing

To translate these standardized field observations into comparable community-level metrics, biological data were processed to ensure consistency across gear types, taxa, and years prior to analysis. When processing bag seine and bay trawl data, both fishes and invertebrates were included in the analyses because these gear types reliably sample a broad suite of estuarine taxa that collectively characterize dominant shoreline and shallow-subtidal assemblages. Gillnet analyses, however, were restricted to finfishes, as this passive sampling gear is designed to target mobile, midwater species and does not provide a consistent or interpretable measure of invertebrate abundance (Hubert et al. 2012).

Rare species can influence diversity metrics in several ways. In long-term monitoring programs, taxa detected only once or a few times may reflect genuinely uncommon estuarine species, transient marine stragglers, or even non-native ornamental fishes released from households. Such sporadic detections provide limited information about community structure, yet they can exert disproportionate effects on richness-based metrics as well as on abundance- or incidence-based dissimilarity measures (Gotelli & Colwell 2001). To reduce the influence of these incidental occurrences, species with fewer than five distinct sampling occasions over the full time series for a given gear were excluded prior to calculating diversity metrics. This procedure is consistent with previous analyses of the same data (Fujiwara et al. 2019, 2022), and reduces noise introduced by non-established or stochastic appearances.

### 2.4 Diversity metrics

Diversity was quantified at three hierarchical levels (Whittaker 1960) to characterize long-term variation in estuarine community structure: α-diversity (within-bay diversity), β-diversity (compositional differentiation among bays or between years), and γ-diversity (regional diversity). All diversity metrics were calculated separately for each sampling gear.

Seasons were defined to align with TPWD sampling protocols. Bag seine and bay trawl samples, which are collected year-round, were grouped into Winter (December–February), Spring (March–May), Summer (June–August), and Fall (September–November). Gill-net samples were assigned only to Spring (April–June) and Fall (September–December), corresponding to the standard deployment windows for this sampling gear (Martínez-Andrade 2018). Gill-net samples collected outside these periods were excluded from analysis.

#### α-diversity estimation

For each gear × bay × season × year stratum, species abundances from all stations were pooled to form a single, composite local assemblage. Because total catch and sampling completeness varied over time and among gears, diversity estimates were standardized using coverage-based rarefaction and extrapolation. For each stratum, the pooled assemblage was standardized to the number of individuals required to reach a target sample coverage (C* =0.98), ensuring that local (α) diversity comparisons were made at equivalent levels of community completeness rather than being confounded by differences in raw sample size (Chao & Jost 2012; Chao et al. 2014).

For pooled station assemblages within each stratum, sample coverage (Ĉ), a measure of the completeness of the resulting sample, was estimated using the Chao–Jost method, which relies on the number of species observed only once (singleton) or twice (doubleton) to standardize diversity estimates to equal coverage. If observed coverage (Ĉ_obs_) met or exceeded the target coverage (C* = 0.98), the assemblage was interpolated to the sample size attaining C*. If Ĉ_obs_ < C*, no extrapolation was performed and the observed sample size was retained. The target value C* = 0.98 provides a consistent level of completeness across assemblages while avoiding instability associated with extrapolation toward full coverage (Chao & Jost 2012).

To estimate local (α) diversity, the pooled assemblage for each stratum was subsampled to the target number of individuals (m*) using a multinomial draw. Hill numbers (q=0,1,2) were then calculated directly on this coverage-standardized composite sample. This ensured comparisons were made at equivalent levels of sampling completeness, representing the diversity of the local bay while limiting the influence of raw sample size differences. Coverage-based diversity estimation was implemented in R using iNEXT (Hsieh et al. 2016).

α-diversity was quantified using Hill numbers, or the “effective number of species”, facilitating standardized comparisons (Jost 2006). Richness (q = 0) reflects the total number of species present, Shannon diversity (q = 1) accounts for both richness and evenness, and dominance-weighted diversity (q = 2; inverse Simpson) emphasizes dominant species. Hill numbers provide a unified framework for comparing diversity across orders that differentially weight rare versus common species.

#### γ-diversity estimation

γ-diversity represents regional diversity, derived by pooling stations coastwide to quantify total species diversity across all bays for a given gear–season–year stratum. As with α-diversity, all γ-diversity estimates were derived from coverage-standardized abundances to ensure that temporal variation reflects biological change rather than differences in sampling effort or detection probability.

To estimate regional diversity, raw species abundances from all stations within a stratum were first pooled to form a single, coastwide regional assemblage. The target sample size for this regional pool was defined as the sum of the coverage-standardized sample sizes (m^∗^, attaining C^∗^=0.98) from the constituent local bays. The fully pooled regional assemblage was then subsampled to this total target size using a single multinomial draw.

γ-diversity was quantified using Hill numbers for q = 0, 1, and 2, and applied to this pooled, coverage-standardized composite sample. Because definitions and interpretations of Hill numbers are identical to those used for α-diversity, they are not repeated here. Diversity metrics were calculated mathematically using custom scripts in base R (R Core Team 2024).

#### β-diversity estimation

β-diversity quantifies compositional differentiation among bays (spatial β-diversity) and changes in community composition through time within bays (temporal β-diversity). All β-diversity metrics were computed under equal-coverage conditions to ensure that observed differences reflect biological variation rather than heterogeneity in sampling effort. Coverage-standardized abundance vectors were derived for each gear × bay × season × year combination and aggregated as required to construct spatial or temporal community matrices.

#### Multiplicative β-diversity based on Hill numbers

Community-level β-diversity was first quantified using multiplicative partitioning of Hill numbers:

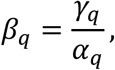

for q = 0, 1, and 2, corresponding to richness-based, Shannon-weighted, and dominance-weighted diversity, respectively. This formulation represents the effective number of distinct assemblages contributing to regional diversity and allows direct comparison of compositional differentiation across diversity orders. All α- and γ-diversity components used in this calculation were coverage-standardized, ensuring comparability across space, time, and sampling gear.

##### Incidence-based temporal and spatial β-diversity (Baselga partitioning)

Temporal β-diversity within bays was evaluated by comparing community composition between consecutive years for each gear × bay × season combination. For each year, coverage-standardized abundances were pooled within bays and converted to presence–absence data to construct a year × species incidence matrix stratified by gear and season. Only adjacent year pairs (Δyear = 1) were retained, capturing short-term compositional change while minimizing confounding from long-term directional trends.

Incidence-based β-diversity was quantified using Baselga’s (2010) partitioning of Sørensen dissimilarity, implemented with the *betapart* package (Baselga & Orme 2012). For any pair of communities, total Sørensen dissimilarity is defined as

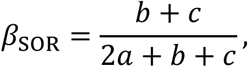

where *a* is the number of shared species and *b* and *c* are the numbers of species unique to each community. Total dissimilarity was partitioned into a turnover component,

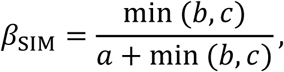

which captures species replacement independent of richness differences, and a nestedness-resultant component,

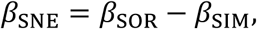

which reflects dissimilarity arising from systematic species loss or gain.

Spatial β-diversity among bays was quantified separately for each gear × season × year combination. Coverage-standardized abundance vectors were pooled within each bay, converted to presence–absence, and assembled into a bay × species incidence matrix. Pairwise β_SOR_, β_SIM_, and β_SNE_ values were computed among all bays within each year using the same definitions as for temporal β-diversity, and then averaged to summarize annual coastwide spatial differentiation.

### 2.5 Dispersal-related traits and guild classification

Species were classified into dispersal guilds to represent differences in typical movement scale and potential for spatial coupling among estuaries. Trait information for fishes was obtained from FishBase (Froese and Pauly 2019) and for invertebrates from SeaLifeBase (Palomares and Pauly 2025), accessed via the *rfishbase* (Boettiger et al. 2012) interface. Taxonomic names were standardized using the *worrms* package (Chamberlain and Vanhoorne 2024) for invertebrates and the “validate_names” function in the *rfishbase* package for fishes, incorporating synonym resolution and conservative fuzzy matching of specific epithets when necessary to maximize trait coverage.

For each species, a suite of traits expected to influence dispersal potential was compiled. For both fishes and invertebrates, these traits included habitat position (pelagic, benthopelagic, demersal), migratory category (oceanodromous, anadromous, catadromous, amphidromous), depth range, reproductive mode, fertilization type, batch spawning, and maximum body size.

Additional traits reflecting movement capacity were incorporated where available, including schooling behavior and adult mobility. For invertebrates, supplementary ecological and life-history traits from SeaLifeBase were used, including the presence of planktonic or pelagic larval stages, evidence of ontogenetic migration between estuarine and shelf or oceanic habitats, and genus-level indicators of high dispersal capacity (e.g., penaeid shrimps and portunid crabs).

Each trait was translated into a directional component score representing its expected contribution to realized dispersal potential, where larger values indicated greater movement capacity or broader spatial connectivity. For fishes, component scores were derived from habitat position (pelagic > benthopelagic/demersal), migratory category (e.g., oceanodromous/anadromous/catadromous > amphidromous > none/unknown), vertical habitat breadth (large depth range), reproductive strategy (non-guarding/broadcast tendencies), fertilization mode (external), batch spawning, and body size (larger maximum length). For invertebrates, the same core traits were supplemented with SeaLifeBase ecology flags that reflect mobility and exposure to advective transport (e.g., mobile/schooling behavior; pelagic or oceanic affinity; estuarine use coupled with offshore/pelagic affinity as an indicator of ontogenetic connectivity). Component scores were summed to obtain a raw dispersal score. Because trait availability varied among taxa and databases, the raw score was then normalized by the maximum attainable score given the subset of traits actually present for each species (i.e., the sum of the weights for available components), yielding a continuous trait-based dispersal index bounded between 0 and 1 and minimizing bias due to missing fields. Normalized scores were mapped to three preliminary dispersal categories using fixed thresholds: Low (<0.35), Intermediate (0.35–0.60), and High (≥0.60). When a normalized score could not be computed because insufficient trait information was available, clear evidence of pelagic connectivity, such as a pelagic habitat designation or pelagic/oceanic ecology flags, was used to assign a provisional High dispersal classification rather than leaving the species unclassified.

To promote biologically conservative and taxonomically coherent guilds, these preliminary trait-based assignments were reconciled with conservative taxonomic priors where higher-level taxonomy provides reliable expectations of dispersal capacity. Priors were specified at the Phylum, Class, or Family level for clades with well-established dispersal constraints or enhancements. For example, taxa with high potential for broad-scale connectivity (e.g., penaeid shrimps, portunid crabs, cephalopods, echinoderms, and pelagic schooling fish families such as Clupeidae, Scombridae, and Carangidae) were assigned High dispersal priors, whereas taxa dominated by sessile or weakly mobile life histories (e.g., sponges, bryozoans, oysters, serpulid polychaetes, and most bivalves) were assigned Low dispersal priors. Gastropods were assigned an Intermediate default prior unless trait evidence indicated otherwise. When both a trait-based category and a taxonomic prior were available, the trait-based category was used by default, but two explicit conflict-resolution rules were applied to avoid implausible assignments. First, taxa with a High dispersal prior were conservatively downgraded to Intermediate only when the normalized multi-trait score was strongly low (<0.35) and no pelagic connectivity signal was present, reflecting the expectation that a single weak trait or incomplete trait coverage should not force a strong reversal of a high-mobility prior. Second, taxa with a Low dispersal prior were upgraded to High only when the normalized multi-trait score was strongly high (≥0.60), ensuring that departures from low-mobility expectations required consistent evidence across multiple traits. Cases in which the trait-based and prior assignments agreed were flagged as “confirmed,” whereas disagreements were recorded as overridden to retain an audit trail of how reconciliation was performed.

A final migration-dominant override was then applied because explicit large-scale migratory behavior implies high realized dispersal regardless of other traits. Species categorized as oceanodromous, anadromous, or catadromous were assigned to the High dispersal guild even if the preliminary reconciled category was lower. Amphidromous species were assigned to at least the Intermediate guild, retaining a High classification when supported by the reconciled evidence. Finally, a small number of species with sparse, contradictory, or ambiguous trait information received expert-knowledge overrides, and these were explicitly marked as manual adjustments to ensure ecological plausibility and transparency.

### 2.6 Thermal-affinity guild classification

Thermal-affinity guilds were assigned to approximate species’ typical thermal niches and to evaluate patterns associated with long-term warming and the increasing representation of warm-adapted taxa. Guild assignments were derived primarily from occurrence-based latitude distributions compiled from online biodiversity databases, rather than relying solely on categorical climate-zone labels. For each species, georeferenced occurrence records (latitude/longitude) were retrieved from the OBIS (2025) using the REST-API (https://api.obis.org). When no usable OBIS coordinates were available, occurrences were obtained from the GBIF (2025) occurrence search API as a fallback. Coordinates were filtered to retain finite values within valid geographic bounds (latitude −90 to 90; longitude −180 to 180), and duplicate coordinate pairs were removed to reduce over-representation of repeatedly sampled locations.

Thermal affiliation was summarized using robust quantiles of the cleaned latitude distribution for each species. Specifically, the 10th percentile latitude (lat₁₀) and 90th percentile latitude (lat₉₀) were calculated, along with the observed minimum and maximum latitudes. A central latitude index was then computed as the midpoint of the quantile bounds,

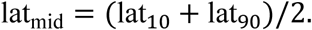

Using quantile-based bounds reduces sensitivity to rare vagrant records and emphasizes the core latitudinal distribution represented by the majority of occurrences.

Species were assigned to a categorical thermal-affinity group based on lat_mid_ using the following thresholds: Tropical (< 15°), Subtropical (15°- <30°), Warm temperate (30° - <45°), and Cold temperate (≥ 45°). Species lacking sufficient occurrence data from both OBIS and GBIF were retained with missing thermal-affinity assignments; downstream analyses used the saved lookup table rather than re-running live API calls.

### 2.7 Environmental variables

Environmental variation was characterized at multiple temporal scales to capture both short-lived events and longer-term shifts in estuarine conditions. To evaluate short-lived phenomena, we defined two distinct classes of short-term extremes: acute disturbances (system-wide named storms and freezes) and environmental spikes (localized 5th/95th percentile anomalies). We contrasted these with chronic multi-year anomalies and a long-term temporal trend.

Environmental observations were collected at the station level and aggregated to the gear × bay × season × year scale. For each combination, seasonal means were calculated for water temperature, salinity, and dissolved oxygen (DO), with station-level variability summarized using standard deviations. Hypoxic exposure within a season was quantified as the proportion of stations with DO < 3 mg L⁻¹.

#### Acute disturbances

Acute disturbances were defined as episodic events capable of generating large and immediate ecological impacts across the Texas coast. These included severe winter freeze events and major storms associated with hurricanes or heavy rainfall. Documented freeze events were identified from published syntheses of historical fish mortality events and recent peer-reviewed assessments of extreme cold impacts, whereas storm events were compiled from official post-storm Tropical Cyclone Reports produced by the NOAA National Hurricane Center.

All named hurricanes and tropical storms affecting the Texas coast during the study period were treated as coastwide events, reflecting the broad spatial scale of associated physical forcing (e.g., air temperature anomalies, wind-driven mixing, storm surge, and freshwater inflow). Each event was assigned to the season in which it occurred and summarized in Table 1.

**Table 1.** Acute disturbance events used to define episodic coastwide forcing along the Texas coast.

| Event name | Year | Month | Description / documented impacts | Primary reference |
| --- | --- | --- | --- | --- |
| **Freeze** |  |  |  |  |
| December 1983 coastwide freeze | 1983 | 12 | Severe coastwide hard freeze; widespread estuarine fish mortality (~14 million individuals) | McEachron et al. 1994 |
| February 1989 mid–lower coast freeze | 1989 | 2 | Severe freeze affecting East Matagorda Bay through Lower Laguna Madre; ~11 million fish killed | McEachron et al. 1994 |
| December 1989 coastwide freeze | 1989 | 12 | Coastwide freeze with extensive fish mortality (~6 million individuals) | McEachron et al. 1994 |
| February 2021 Winter Storm Uri | 2021 | 2 | Extreme cold event causing widespread estuarine fish mortality (~3.8–3.9 million individuals) | Texas Parks and Wildlife Department 2021 |
| **Storm** |  |  |  |  |
| Hurricane Allen | 1980 | 8 | Major hurricane affecting south Texas; wind, surge, and mixing | NOAA NHC |
| Hurricane Alicia | 1983 | 8 | Category 3 Galveston landfall; severe wind and surge | NOAA NHC |
| Hurricane Chantal | 1989 | 8 | Category 1 landfall near High Island; wind and rainfall | NOAA NHC |
| Hurricane Jerry | 1989 | 10 | Category 1 Galveston landfall; surge and wind | NOAA NHC |
| Hurricane Dean | 1995 | 8 | Tropical cyclone affecting south Texas coast | NOAA NHC |
| Hurricane Bret | 1999 | 8 | Category 4 landfall in south Texas; intense wind and surge | NOAA NHC |
| Tropical Storm Allison | 2001 | 6 | Extreme rainfall and flooding across upper Texas coast | NOAA NHC |
| Hurricane Rita | 2005 | 9 | Category 3 storm near TX–LA border; major surge | NOAA NHC |
| Hurricane Ike | 2008 | 9 | Category 2 Galveston landfall; extensive surge and mixing | NOAA NHC |
| Hurricane Harvey | 2017 | 8 | Category 4 landfall near Port Aransas; extreme rainfall and freshwater inflow | NOAA NHC |
| Tropical Storm Imelda | 2019 | 9 | Catastrophic rainfall across southeast Texas | NOAA NHC |
| Hurricane Hanna | 2020 | 7 | Category 1 south Texas landfall; surge and wind | NOAA NHC |
| Hurricane Beryl | 2024 | 7 | Category 1 storm affecting Texas coast; rainfall and surge | NOAA NHC |

For each gear × bay × season × year combination, binary indicators were created to denote exposure to freeze or major storm events. When an event indicator showed no variation within a given gear-specific dataset (e.g., all zeros), it was omitted from the corresponding model to avoid rank-deficient fits.

#### Chronic environmental anomalies

Chronic environmental conditions were estimated from in situ measurements of water temperature, salinity, and dissolved oxygen collected during routine TPWD sampling for all gears. To quantify gradual, persistent departures from baseline conditions, 5-year right-aligned rolling means were calculated separately for each gear × bay × season combination. Partial rolling windows were permitted at the beginning of each time series.

Each rolling mean was then expressed as an anomaly relative to the long-term climatological mean for the corresponding gear, bay, and season. This rolling-window approach captures sustained environmental change while reducing sensitivity to short-term variability. Sampling occasions lacking temperature, salinity, or DO measurements were excluded from rolling-mean calculations to avoid artificial smoothing or interpolation.

#### Within-season environmental spike metrics

To represent fine-scale exposure to short-term conditions within individual years, spike metrics were derived from station-level environmental observations. For temperature and salinity, long-term thresholds were defined as the 5th and 95th percentiles calculated separately for each gear × bay × season across the full time series. For dissolved oxygen, hypoxic exposure was defined using a threshold of 3 mg L⁻¹.

For each gear × bay × season × year combination, we calculated the proportion of sampling stations experiencing unusually low or high temperature, low or high salinity, or hypoxic conditions. These spike metrics quantify the spatial extent and intensity of exposures to short-term extremes that may not strongly influence multi-year averages but can have substantial ecological effects within a given year.

#### Long-term temporal trend

Long-term directional change was incorporated directly into all models through the inclusion of year (mean-centered across the monitoring period). This term captures sustained system-wide trends, such as gradual warming, long-term salinity shifts, or progressive changes in species pools, while allowing the effects of acute disturbances and short-term anomalies to be evaluated independently.

### 2.8 α- and γ-diversity trend models

Long-term temporal trends in diversity were evaluated using coverage-standardized α- and γ-diversity estimates expressed as Hill numbers (q = 0, 1, 2), which quantify richness (q₀), Shannon diversity (q₁), and dominance-weighted diversity (q₂) on a common effective-species scale. All analyses were conducted separately for each sampling gear (bag seine, bay trawl, and gillnet) to avoid conflating gear-specific sampling properties and sampling domains.

#### α-diversity: coastwide trends with bay-level replication

For each sampling gear and diversity order, α-diversity trends were analyzed using bay-level observations indexed by bay × season × year. Two complementary model classes were used, each addressing a distinct aspect of temporal change.

##### Linear mixed-effects models: overall directional trends

To quantify the overall direction and magnitude of long-term change, α-diversity trends were estimated using linear mixed-effects models (LMMs) with year and season as fixed effects and bay as a random intercept:

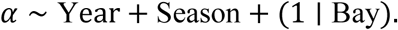

These models estimate a single coastwide temporal slope for each gear and diversity order while accounting for seasonal differences in mean α-diversity and repeated observations within bays. The year coefficient, representing the average annual rate of change, was multiplied by ten to report decadal trends. LMMs were fitted using maximum likelihood as implemented in the lme4 framework (Bates et al. 2015).

##### Generalized additive models: nonlinear temporal structure

To evaluate whether α-diversity trajectories exhibited nonlinear temporal structure, generalized additive models (GAMs) were fitted separately for each gear and diversity order:

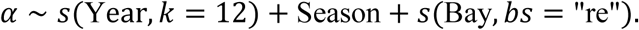

These models allow flexible, smooth temporal responses while retaining seasonal effects and accounting for bay-level heterogeneity through a penalized random-effect term. GAMs were fitted using restricted maximum likelihood in mgcv (Wood 2017). For visualization, marginal temporal trends common across bays were generated by excluding the bay random-effect term during prediction, allowing direct comparison of seasonal trajectories for each gear-type.

#### γ-diversity: coastwide trends by season

Regional γ-diversity was evaluated using coverage-standardized Hill numbers calculated at the coastwide scale. For each gear type, a single γ-diversity estimate was retained for each contribution of season and year. Because γ-diversity is not replicated among bays, no spatial random effects were included. As with α-diversity, two complementary model classes were used.

##### Linear models: overall directional trends

To quantify overall temporal trends in γ-diversity, linear models were fitted separately for each gear and diversity order. When data from two or more seasons were available, models included both year and season:

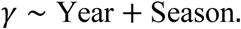

In these models, season was treated as a categorical factor affecting baseline γ-diversity, while a single temporal slope was estimated across seasons. This structure accommodates gear-specific seasonal sampling patterns (e.g., gillnet γ-diversity was estimated for spring and fall only) while focusing inference on the overall direction of long-term change.

##### Generalized additive models: nonlinear temporal structure

To assess potential nonlinear temporal patterns in γ-diversity, GAMs were fitted separately for each gear and diversity order. Models were specified as:

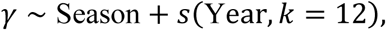

with the seasonal term omitted when not supported by the data. These models were used to visualize temporal trajectories and departures from linear trends, while retaining seasonal effects when they improved the model in a meaningful way.

To account for multiple comparisons across diversity orders and gear types, p-values associated with temporal slopes were adjusted using the false discovery rate (FDR) procedure (Benjamini and Hochberg 1995).

### 2.9 Guild-specific diversity trends

Guild-level analyses were conducted separately for dispersal groups (Low, Intermediate, High) and thermal-affinity groups (Tropical, Subtropical, Temperate) to assess whether long-term diversity trends differed systematically among functional categories. For each guild dimension, coverage-standardized Shannon α- and γ-diversity (q = 1) were analyzed to estimate distinct long-term slopes for every gear–guild combination. Models included fixed effects for year (continuous), gear, season, guild group, and all lower-order interactions, including the year × gear × guild interaction. Because local (α) diversity is replicated among bays, it was analyzed using linear mixed-effects models (LMMs) with bay (as delineated by TPWD) included as a random intercept to account for persistent spatial differences:

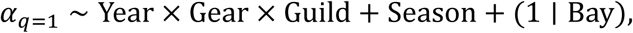

In contrast, because regional (γ) diversity is pooled coastwide, it was analyzed using standard linear models without spatial random effects:

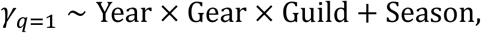

Consistent with recommended practice for mixed-effects models, inference focused on estimated temporal slopes and their associated uncertainty for each gear–guild combination, as these quantities provide more interpretable and robust measures of long-term change than model-level p-values (Bolker et al. 2009). To account for multiple comparisons across the extracted gear–guild trajectories, p-values associated with temporal slopes were adjusted using the false discovery rate (FDR) procedure.

### 2.10 β-Diversity trends and Baselga decomposition

Long-term trends in spatial and temporal β-diversity were evaluated using two complementary metric families: (i) Hill-number–based multiplicative β-diversity (q = 0, 1, 2) and (ii) incidence-based Baselga dissimilarity and its turnover and nestedness components. Definitions and calculation procedures for β-diversity metrics are provided in Section 2.4; this section describes statistical modeling and inference. Linear models were fitted using ordinary least-squares regression (the “lm” function), and linear mixed-effects models were fitted using the lme4 package, with inference provided by the lmerTest package (Bates et al. 2015; Kuznetsova et al. 2017).

#### Spatial β-diversity trends from multiplicative partitioning

For each diversity order (q = 0, 1, 2), coastwide temporal trends in spatial differentiation among bays were evaluated using linear models:

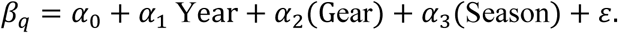

Here, *β*_*q*_ is multiplicative spatial β-diversity for a given bay × gear × season × year observation; Year is treated as a continuous predictor; Gear and Season are categorical fixed effects. The coefficient *⍺*_1_ estimates the long-term linear trend (per year), and decadal rates were obtained as 10*⍺*_1_. Uncertainty in *⍺*_1_ was summarized using nonparametric bootstrap resampling of rows (500 resamples) to obtain a bootstrap standard error and 95% percentile confidence interval.

#### Guild-specific spatial β-diversity (q = 2)

To evaluate whether dominance-weighted spatial differentiation varied among functional groups, spatial β-diversity at q = 2 (*β*_*q*=2_) was analyzed separately for dispersal and thermal-affinity guild systems using linear mixed-effects models:

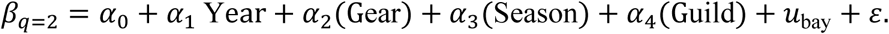

Here, guild denotes the guild category within a guild system (e.g., Low/Intermediate/High dispersal or Tropical/Subtropical/Temperate thermal affinity), and *u*_bay_ is a random intercept for bay to account for among-bay differences.

#### Temporal β-diversity from Baselga dissimilarity (Δyear = 1; non-guild)

Temporal β-diversity within bays was quantified using Baselga’s total Sørensen dissimilarity between consecutive years (Δyear = 1). Interannual dissimilarity values were analyzed using mixed-effects models:

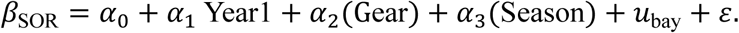

Here, *β*_SOR_ is temporal dissimilarity for a bay × gear × season between Year1 and Year2 **=** Year1 + 1, and *u*_bay_ is a bay random intercept. The coefficient *⍺*_1_ estimates long-term change in year-to-year compositional turnover.

#### Spatial Baselga components: total, turnover, and nestedness (non-guild)

To identify mechanisms underlying changes in spatial differentiation, multi-site incidence-based dissimilarity among bays was decomposed into total dissimilarity, turnover, and nestedness. Each component was modeled using:

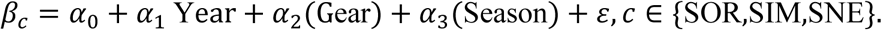

Here, *β*_*c*_ is the annual multi-site component for a gear × season × year observation (total SOR, turnover SIM, nestedness SNE). Consistent with the approach used for multiplicative spatial β-diversity, uncertainty in *⍺*_1_ was summarized using 500-resample nonparametric bootstrap standard errors and 95% percentile confidence intervals.

#### Guild-specific Baselga spatial and temporal analyses

Baselga metrics were further analyzed within dispersal and thermal guild systems to assess whether mechanisms differed among functional groups.

Guild-specific spatial Baselga components were modeled with linear models including guild as a categorical predictor:

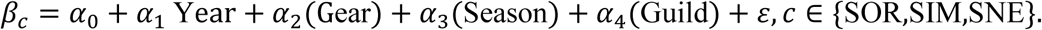

Bootstrap resampling (500 resamples) was used to summarize uncertainty in *⍺*_1_ for these guild-specific spatial component models.

Guild-specific temporal β_SOR_ between consecutive years (Δyear = 1) was modeled with a logit transform to accommodate bounded dissimilarity values:

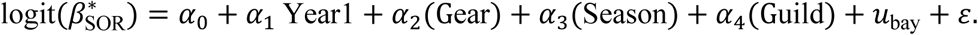

Here, 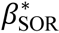 is *β*_SOR_ clipped to [*∈*, 1 − *∈*](with *∈* = 10^−6^) prior to transformation, and *u*_bay_ is a bay random intercept.

#### Guild-level slope comparison analyses

To test whether rates of change differed among guild categories, interaction models were fitted with Year **×** Guild (or Year1 **×** Guild) terms for: (i) dominance-weighted spatial β-diversity (*β*_*q*=2_) within dispersal and thermal affinity systems, (ii) guild-specific spatial Baselga components, and (iii) guild-specific temporal *β*_SOR_(logit scale). Guild-specific slopes were estimated using marginal trend estimation implemented with the “emtrends” function in the emmeans package (Lenth 2024), and pairwise differences in slopes among guild categories were evaluated with post hoc contrasts. Multiple comparisons were adjusted using Holm’s method. Parallel slope-difference tests for *β*_*q*=2_ were conducted to see if guild-specific rate of homogenization differed among the sampling gear types.

#### Model diagnostics

Model adequacy was evaluated by refitting each β-diversity model using the same specification and analysis dataset, with residuals versus fitted values and quantile–quantile plots used to assess residual behavior, and bay-level random-intercept distributions examined for mixed-effects models.

### 2.11 GAM-based derivative diagnostics for changes in diversity trajectories

Generalized additive models (GAMs) were used to characterize non-linear temporal patterns in diversity time series and to identify years in which the direction or curvature of long-term trends changed. Analyses were conducted separately for each gear × season combination using a population-smooth formulation that retained bay-level replication and explicitly accounted for persistent spatial differences among bays via a random-effect smooth.

#### Response metrics and data resolution

Three diversity metrics were analyzed: α-diversity as coverage-standardized Shannon diversity (Hill number *q* = 1), β-diversity as multiplicative β-diversity (*q* = 1), and γ-diversity as coverage-standardized richness (*q* = 0). The response data were maintained at the bay × gear × season × year resolution, such that multiple bay-level observations informed each annual population-level trend. Year was treated as an integer-valued continuous predictor.

#### GAM specification

For each gear × season time series, GAMs of the form

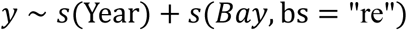

were fit using the mgcv package with restricted maximum likelihood (REML). The smooth term *s*(Year)represents the population-level temporal trajectory shared across bays, while *s*(Bay, bs = “re”) accounts for bay-specific differences in mean diversity. The basis dimension for the temporal smooth was selected adaptively as a function of the number of sampled years in each series, with safeguards to prevent overfitting in shorter time series. Series with insufficient temporal replication were excluded from derivative diagnostics.

#### Error distributions and response constraints

α-diversity was modeled using a Gaussian error distribution with an identity link. β- and γ-diversity were modeled using Gamma error distributions with log links. To satisfy the strict positivity requirement of the Gamma family, response values were lower-bounded prior to model fitting using a small constant. Effective degrees of freedom (edf) and p-values for the temporal smooth were extracted from model summaries as measures of statistical support for non-linear temporal structure.

#### Derivative estimation and change-point diagnostics

Temporal derivatives of the fitted population smooths were computed using the gratia package on a dense grid spanning the observed year range (Simpson 2023). Two complementary derivative-based diagnostics were derived.

First, changes in trend direction were identified using the first derivative of the smooth. Derivative sign was classified conservatively: the derivative was considered positive only when its confidence interval lay entirely above zero, negative only when entirely below zero, and otherwise indeterminate. Short runs of indeterminate sign were bridged by propagating the nearest non-zero sign, and candidate sign-change years were defined as years where the resulting non-zero derivative sign differed from the preceding interval.

Second, periods of strongest curvature were identified using the second derivative of the smooth. Peaks in the absolute value of the second derivative were interpreted as acceleration or deceleration extrema, subject to minimum temporal separation and significance constraints to avoid selecting spurious or closely spaced peaks.

#### Multiple-testing control and diagnostics

Within each diversity metric (α, β, γ), p-values associated with the temporal smooth were adjusted for multiple testing using the Benjamini–Hochberg false discovery rate procedure. Model adequacy was evaluated using quantile–quantile plots of deviance residuals and additional diagnostic checks to confirm appropriate smoothness, residual behavior, and convergence.

### 2.12 Chronic versus acute environmental drivers of biodiversity

To evaluate how persistent environmental change and short-lived extreme events influence estuarine biodiversity, coverage-standardized Shannon diversity (Hill number q = 1) was related to (i) chronic multi-year environmental anomalies and (ii) acute event exposure and within-season spike metrics derived from temperature, salinity, and dissolved oxygen (DO), as described in Section 2.7. Analyses were conducted separately for each sampling gear (BS, BT, GN) using bay-level observations indexed by bay × season × year stratum, consistent with the α–β–γ analytical framework.

#### Biodiversity responses and data integration

For each gear × bay × season × year stratum, coverage-standardized Shannon diversity was assembled for local diversity *A*_*g*,*b*,*s*,*t*_(α-diversity), multiplicative spatial turnover *B*_*g*,*b*,*s*,*t*_(β-diversity), and regional diversity *G*_*g*,*b*,*s*,*t*_(γ-diversity). Because *B*_*g*,*b*,*s*,*t*_ > 0 by definition, β-diversity was analyzed on the log scale:

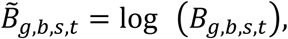

where *B̃*_*g*,*b*,*s*,*t*_ denotes log-transformed multiplicative β-diversity. Calendar year was centered prior to analysis.

##### Stage 1: selection of acute-event encoding and spike inclusion

For each sampling gear and response variable (*A*, *B̃*, *G*), candidate models incorporating alternative representations of acute disturbance events, with and without spike metrics, were compared using AIC. Two fixed-effect specifications were evaluated for each acute-event encoding:

**Base model (chronic + acute):**

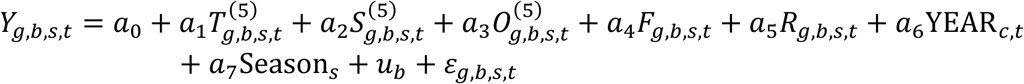

**Spike-augmented model (chronic + acute + spike):**

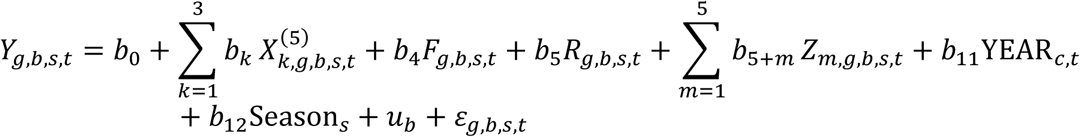

where:

###### Response variables

- *Y*_*g*,*b*,*s*,*t*_ denotes one of:

- *A*_*g*,*b*,*s*,*t*_ : local (α) diversity
- *B̃*_*g*,*b*,*s*,*t*_ : spatial β-diversity
- *G*_*g*,*b*,*s*,*t*_ : regional (γ) diversity

###### Chronic environmental anomalies

- *T*^(5)^ : 5-year rolling temperature anomaly
- *S*^(5)^ : 5-year rolling salinity anomaly
- *O*^(5)^ : 5-year rolling dissolved oxygen anomaly

###### Acute disturbance indicators

- *F* : freeze exposure (binary, encoding-specific)
- *R* : storm exposure (binary, encoding-specific)

###### Within-season spike metrics

- *Z*_*m*_ represents within-season spikes, including:

- hypoxia extent (DO < 3 mg L⁻¹)
- low- and high-temperature spikes
- low- and high-salinity spikes

###### Temporal structure

- YEAR_*c*_ : centered calendar year
- Season : categorical seasonal effect

###### Random effects and errors

- 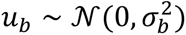 : bay-level random intercept (α and β models only)
- *ɛ* : residual error

Models were fit separately by sampling gear. For α-diversity and spatial β-diversity, bay was included as a random intercept and models were fit using maximum likelihood (REML = FALSE). γ-diversity models were fit without a bay random effect because γ-diversity is defined at the regional scale. For each response × gear combination, the best-supported acute-event encoding and spike inclusion was selected based on minimum AIC.

##### Stage 2: driver-class importance within the selected structure

Using the best-supported Stage 1 structure for each response × gear combination, State 2 evaluated relative support for major driver classes by fitting nested reduced models. Starting from the full model, reduced variants removed: (i) centered year, (ii) chronic anomalies, (iii) acute disturbance indicators (freeze and storm jointly), (iv) spike metrics (when included), and (v) centered year and chronic anomalies jointly. Inference focused on ΔAIC relative to the full model within each response × gear stratum.

#### Model diagnostics

A standardized diagnostic workflow was applied to a representative subset of fitted models (one per gear × response), using the spike-augmented specification with a 12-month duration encoding as the reference model where applicable. Diagnostics included residuals-versus-fitted plots and normal quantile–quantile plots. For mixed-effects models, singular fits were flagged and marginal and conditional *R*^2^ values were computed when available. The rolling calculations were conducted using the zoo package (Zeileis & Grothendieck 2005), and mixed-effects modeling was supported by lmerTest (Kuznetsova et al. 2017).

### 2.13 Use of Generative AI

ChatGPT (OpenAI) was used as a language model to assist with editing and refining manuscript text and to help identify and correct coding errors in R during data analysis. The authors take full responsibility for all analytical decisions, results, and interpretations.

## 3. Results

Across the full monitoring record, the three types of sampling gear captured distinct community assemblages with clear differences in both taxonomic composition and functional structure (Table 2). In total, 266 unique taxa were recorded in bag seine samples, 306 unique taxa were recorded in bay trawl samples, and 116 unique taxa were recorded in gillnet samples. Also, distinct functional assemblages were associated with each sampling gear type. Low-dispersal taxa comprised the largest guild in both bag seine and bay trawl assemblages, while gillnet samples included a more balanced representation of low-, intermediate-, and high-dispersal taxa. Thermal affiliations spanned tropical through cold-temperate groups in bag seine and bay trawl assemblages, though subtropical taxa dominated in both. Gillnet assemblages similarly included tropical, subtropical, and warm-temperate taxa but did not contain any cold-temperate species.

**Table 2.**
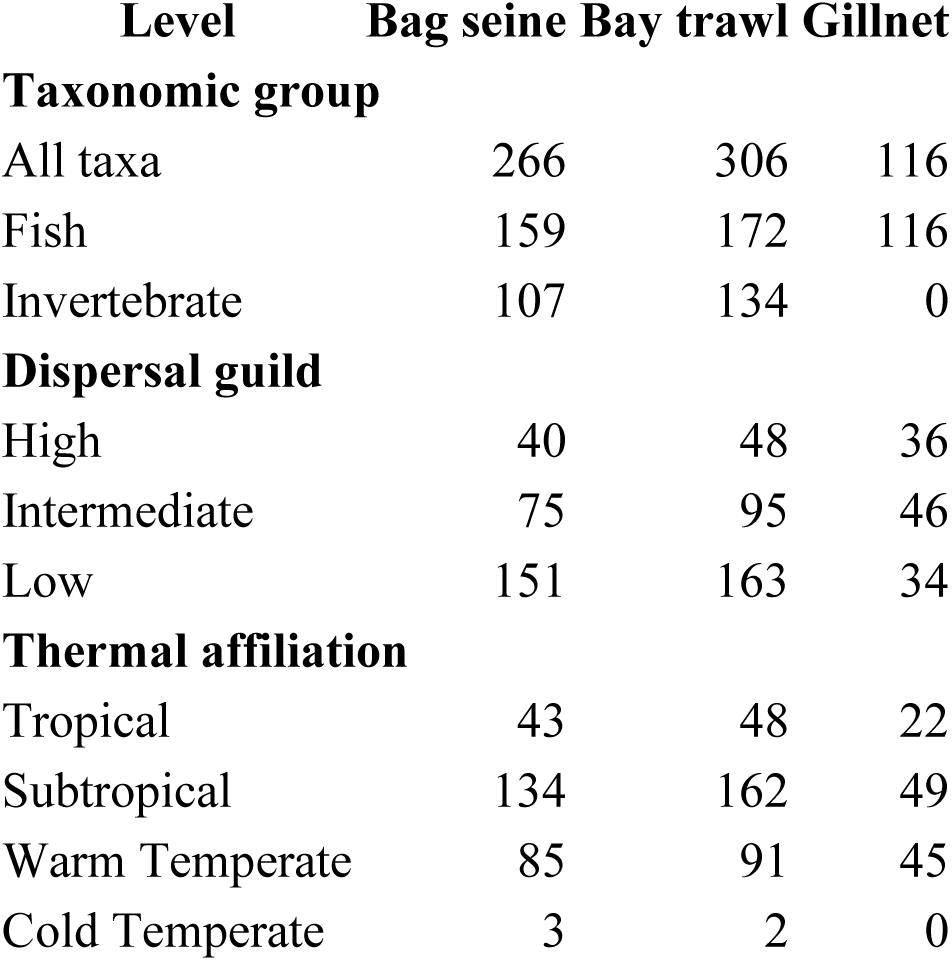
Taxonomic and functional composition of assemblages by sampling gear. Numbers indicate the total count of unique taxa contributing to each taxonomic group, dispersal guild, or thermal affiliation within bag seine, bay trawl, and gillnet assemblages across the full monitoring record. These summaries describe the composition of guild inputs used in subsequent analyses and do not represent temporal trends or response direction.

| **Level** | **Bag seine** | **Bay trawl** | **Gillnet** |
| --- | --- | --- | --- |
| **Taxonomic group** |  |  |  |
| All taxa | 266 | 306 | 116 |
| Fish | 159 | 172 | 116 |
| Invertebrate | 107 | 134 | 0 |
| **Dispersal guild** |  |  |  |
| High | 40 | 48 | 36 |
| Intermediate | 75 | 95 | 46 |
| Low | 151 | 163 | 34 |
| **Thermal affiliation** |  |  |  |
| Tropical | 43 | 48 | 22 |
| Subtropical | 134 | 162 | 49 |
| Warm Temperate | 85 | 91 | 45 |
| Cold Temperate | 3 | 2 | 0 |

### 3.1 Long-Term Trends in Local (α) and Regional (γ) Diversity

α- and γ-diversity exhibited strong, gear-specific long-term directional trends (Table S1). Linear mixed-effects models revealed consistent increases in α-diversity for bag seine (BS) samples across all diversity orders, with significant positive slopes for richness (q = 0), Shannon diversity (q = 1), and dominance-weighted diversity (q = 2), corresponding to decadal increases of 3.16 ± 0.12, 1.43 ± 0.07, and 0.94 ± 0.06 effective species, respectively (all FDR-corrected p ≪ 0.001). In contrast, bay trawl (BT) α-diversity showed weak but directionally consistent declines, with significant negative trends for richness (decadal slope = −0.79 ± 0.14) and Shannon diversity (decadal slope = −0.26 ± 0.10) and no detectable trend in dominance-weighted diversity after FDR correction. Gillnet (GN) α-diversity exhibited intermediate but consistently positive trends across all diversity orders, with decadal increases of 0.83 ± 0.09, 0.43 ± 0.06, and 0.34 ± 0.05 effective species for richness, Shannon, and dominance-weighted diversity, respectively. These patterns indicate that long-term changes in local diversity differed systematically by sampling gear, reflecting habitat domains.

Generalized additive models (GAMs) provided complementary visualization of temporal structure in α-diversity across seasons and diversity orders (Fig. 2; Figs. S1, S2). Across Hill diversity orders, α-diversity exhibited broadly consistent temporal patterns (q = 0–2). Bag seine samples showed steady increases across all seasons, with more pronounced gains emerging after approximately 2010, whereas bay trawl samples exhibited relatively stable or weakly declining trajectories across seasons. Gillnet α-diversity increased gradually in both spring and fall beginning in the late 1990’s, with moderate interannual variability. Together, these results indicate that richness, Shannon diversity, and dominance-weighted diversity convey concordant signals of long-term change in local diversity across sampling gears.

**Figure 2.**
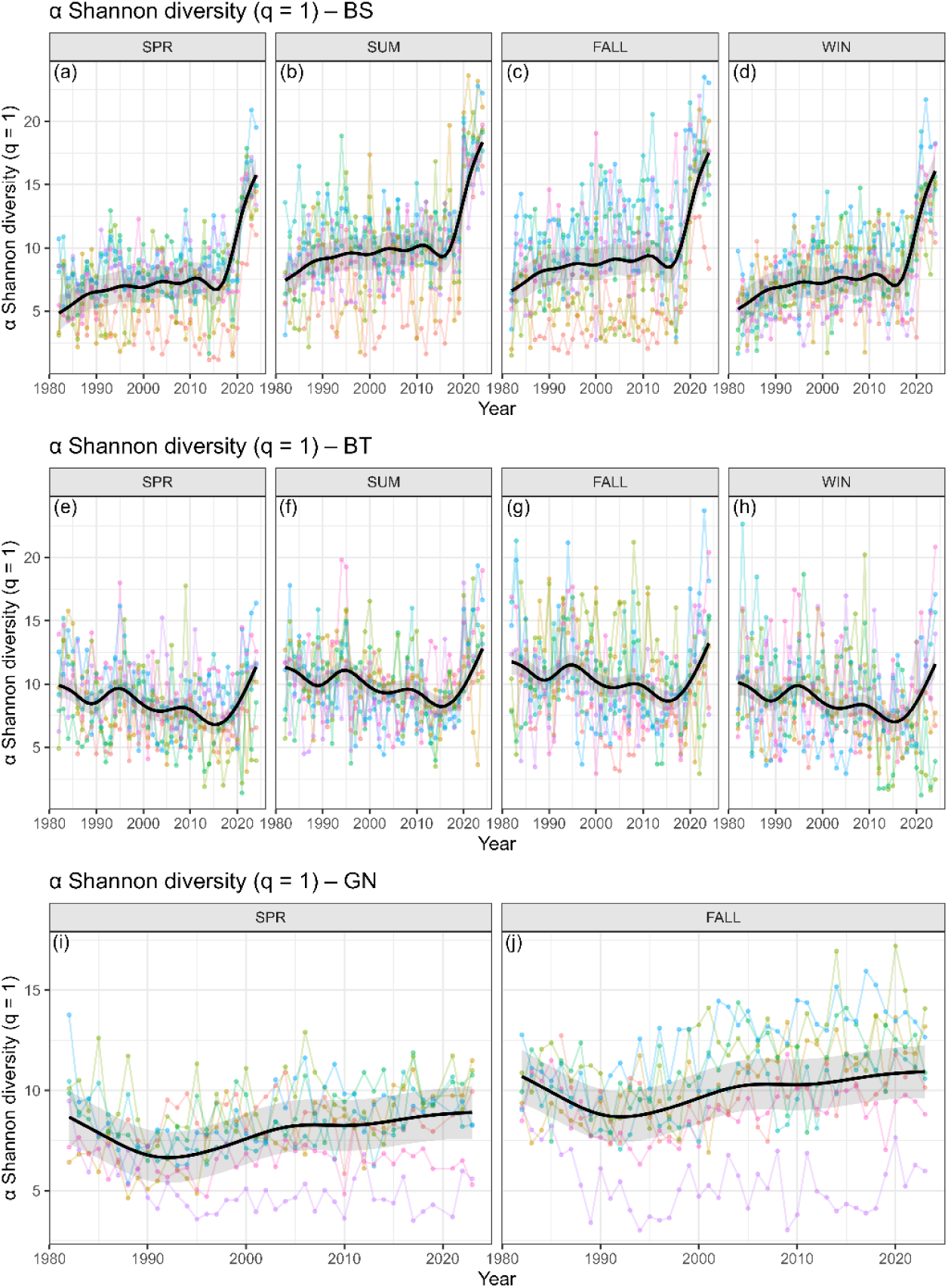
Long-term trends in local (α) diversity (Shannon q = 1) across gears and seasons, 1982–2024. Points show annual, coverage-standardized estimates of Shannon α-diversity (Hill number q = 1) for individual bays, and solid black lines show fitted generalized additive model (GAM) population smooths with shaded ribbons indicating 95% confidence intervals. Panels correspond to gear–season combinations: bag seine (BS; panels a–d: spring, summer, fall, winter), bay trawl (BT; panels e–h: spring, summer, fall, winter), and gillnet (GN; panels i–j: spring, fall). Colors distinguish individual bays. Local diversity increased steadily across all seasons for bag seine and gillnet assemblages, reflecting sustained gains in local richness and evenness, whereas bay trawl α-diversity exhibited weaker or negative temporal trends across seasons, indicating that long-term changes in local community structure differ systematically among habitat domains.

Patterns at the regional scale closely paralleled those observed locally but were larger in magnitude (Table S1, Fig. 3). γ-Shannon diversity (q = 1) increased strongly for bag seine samples across all seasons (decadal slope = 1.92 ± 0.23), declined consistently for bay trawl samples (decadal slope = −1.16 ± 0.21), and increased modestly for gillnet samples (decadal slope = 0.55 ± 0.09). GAM smooths indicated gradual temporal change, with pronounced post-2000 increases in bag seine γ-diversity, whereas bay trawl γ-diversity showed long-term stagnation or decline across all seasons (Fig. 3). γ-richness (q = 0) and γ-dominance-weighted diversity (q = 2) showed similar gear-specific patterns over time (Figs. S3, S4), with strong and sustained increases for bag seine samples (decadal slopes: q = 0, 7.28 ± 0.49; q = 2, 1.24 ± 0.17), declining or weakly negative trends for bay trawl samples (decadal slopes: q = 0, −1.16 ± 0.54; q = 2, −0.67 ± 0.15), and intermediate increases for gillnet samples (decadal slopes: q = 0, 1.71 ± 0.30; q = 2, 0.39 ± 0.08).

**Figure 3.**
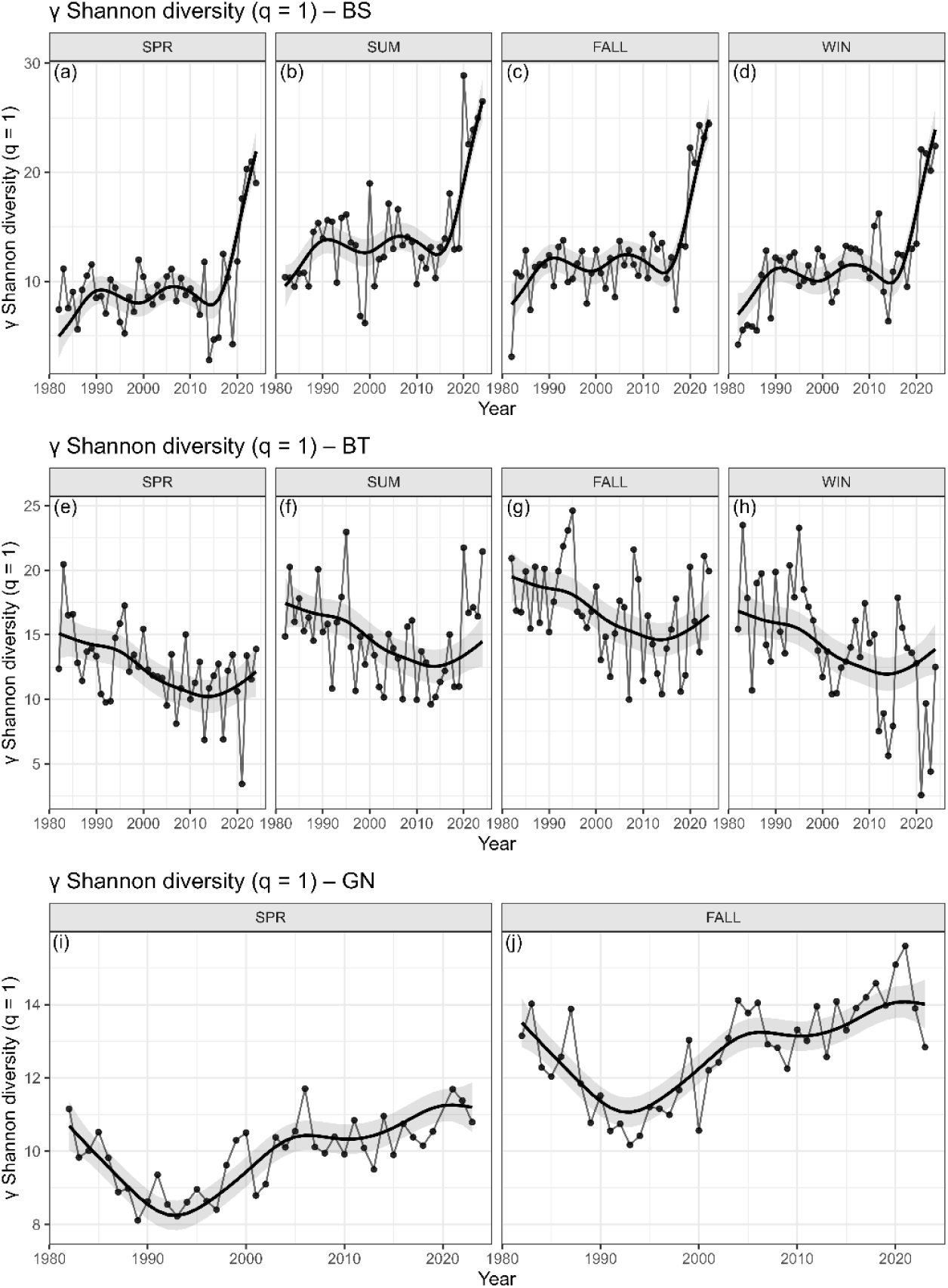
Long-term trends in regional (γ) diversity (Shannon q = 1) across gears and seasons, 1982–2024. Points show annual, coverage-standardized estimates of Shannon γ-diversity (Hill number q = 1) pooled coastwide, and solid lines show fitted GAM population smooths with shaded ribbons indicating 95% confidence intervals. Panels correspond to gear–season combinations: bag seine (BS; panels a–d: spring, summer, fall, winter), bay trawl (BT; panels e–h: spring, summer, fall, winter), and gillnet (GN; panels i–j: spring, fall). Regional diversity increased strongly and consistently across seasons for bag seine assemblages and increased modestly for gillnet assemblages, indicating sustained expansion of the coastwide species pool in shallow habitats. In contrast, bay trawl γ-diversity declined consistently across seasons, paralleling the negative α-diversity trends observed for this gear and indicating long-term restructuring of deeper-water assemblages rather than a sampling artifact.

α- and γ-diversity trends varied by gear type, yet were directionally consistent across all diversity orders within each gear. Because each sampling gear targets a distinct habitat domain, α- and γ-diversity were evaluated within gear-specific regional pools. Within these habitat domains, repeated increases or declines at local scales occurred alongside corresponding changes in regional diversity over time, as reflected by the concordance between linear trend estimates and GAM-based temporal smooths.

### 3.2 Guild-Level Contributions to Increasing Diversity

#### Dispersal guilds

Coverage-standardized Shannon diversity (q = 1) exhibited strong, gear-dependent differences in long-term trends among dispersal guilds at both local (α) and coastwide (γ) scales (Fig. S5; Fig. 4; Table S2). Across all guilds, temporal slopes were consistently positive for bag seine (BS), negative or weak for bay trawl (BT), and generally weak for gillnet (GN), but the magnitude of change varied systematically among dispersal groups. The strongest and most consistent gains occurred for intermediate- and low-dispersal taxa in BS assemblages, which showed the largest positive slopes at both α- and γ-diversity scales, while high-dispersal taxa exhibited comparatively muted responses across all gears, suggesting greater temporal stability or reduced sensitivity to the broader processes underlying the observed community shifts, potentially due to their higher mobility. In BT assemblages, declines were concentrated in intermediate- and low-dispersal guilds at both scales, whereas high-dispersal taxa showed no detectable trend. Complete gear–guild slope estimates and associated uncertainty are provided in Table S2.

**Figure 4.**
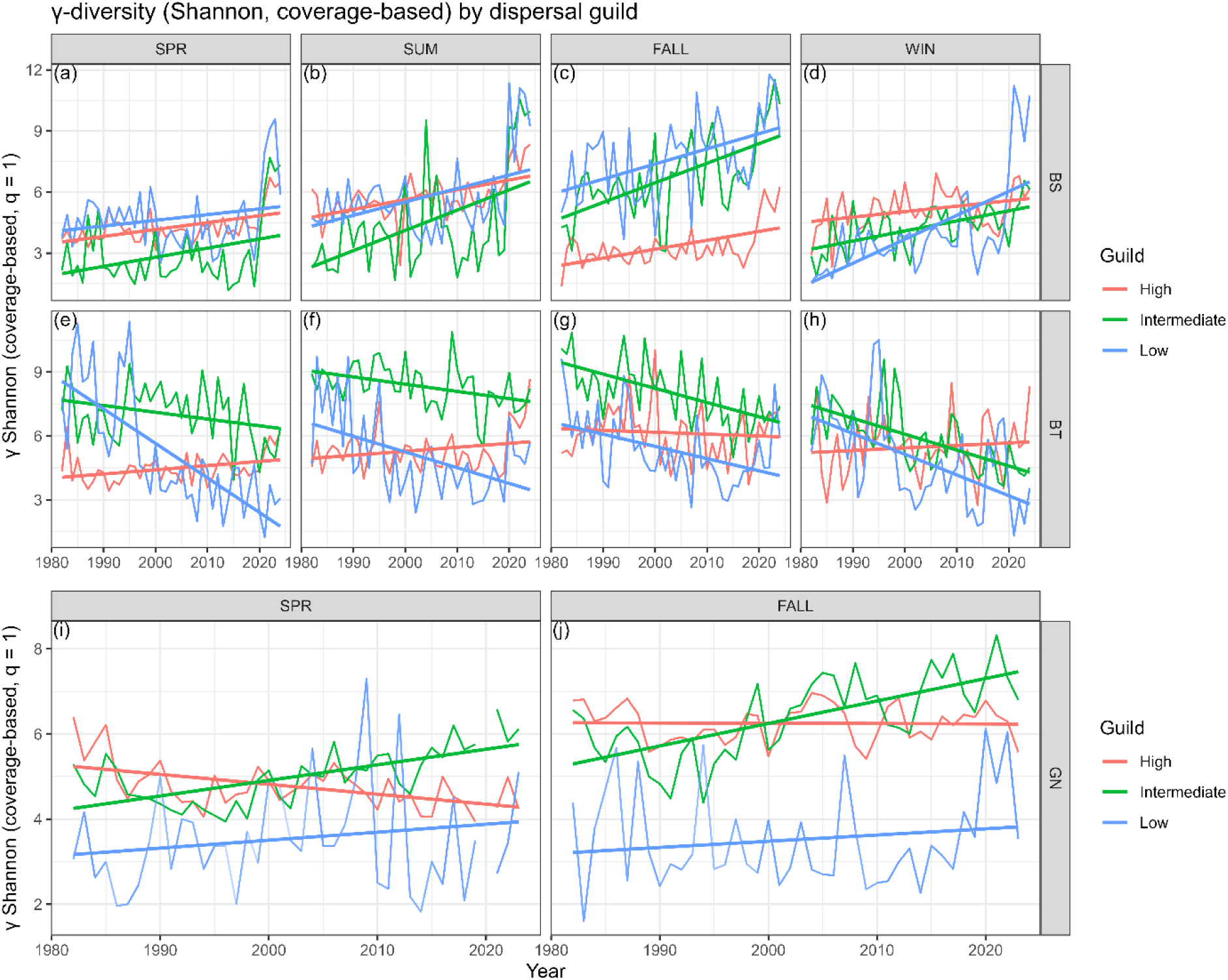
Long-term trends in coverage-standardized Shannon γ-diversity (q = 1) for dispersal guilds across gears and seasons, 1982–2024. Thin lines show annual γ-diversity trajectories for each dispersal guild (High, Intermediate, Low), and thick lines show linear trends estimated from mixed-effects models. Panels correspond to gear–season combinations: bag seine (BS; panels a–d: winter, spring, summer, fall), bay trawl (BT; panels e–h: winter, spring, summer, fall), and gillnet (GN; panels i–j: spring, fall). Colors distinguish dispersal guilds: red = High, green = Intermediate, blue = Low.

Patterns at the γ-diversity scale largely mirrored those observed for α-diversity but were amplified in magnitude (Fig. 4; Table S2). The most pronounced contrast was between low-dispersal taxa in BS assemblages, which showed strong γ-diversity increases (slope = 0.0711 yr⁻¹), and low-dispersal taxa in BT assemblages, which showed the steepest declines of any guild–gear combination (slope = −0.0987 yr⁻¹). Together, these results indicate that long-term community reorganization was strongly associated with dispersal capacity, with intermediate-and low-dispersal taxa driving the strongest habitat-specific changes while high-dispersal taxa remained comparatively stable across habitats.

#### Thermal-affinity guilds

Coverage-standardized Shannon diversity (q = 1) exhibited pronounced, gear-dependent differences among thermal-affinity guilds, with subtropical taxa showing the strongest and most consistent increases over time, particularly in shallow, nearshore habitats (Fig. S6; Fig. 5; Table S3). In BS assemblages, subtropical taxa drove the largest α- and γ-diversity gains across all seasons, while temperate taxa showed moderate increases in BS but sharp declines in BT, the strongest negative signal among all thermal guilds. Tropical taxa showed comparatively modest and variable responses across gears and diversity scales. Complete gear–guild slope estimates and associated uncertainty are provided in Table S3.

**Figure 5.**
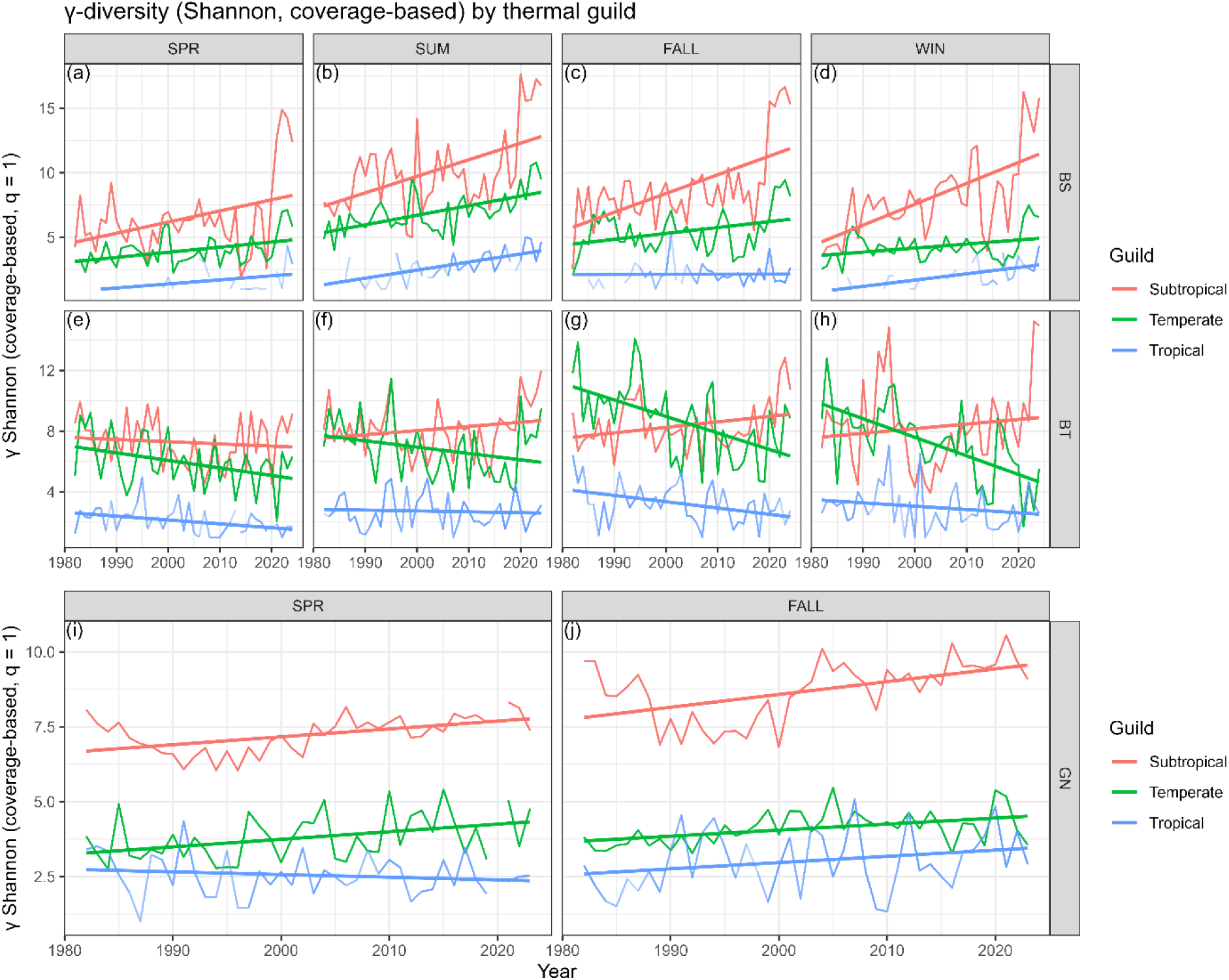
Long-term trends in coverage-standardized Shannon γ-diversity (q = 1) for thermal-affinity guilds across gears and seasons, 1982–2024. Thin lines show annual γ-diversity trajectories for each thermal guild, and thick lines show linear trends estimated from mixed-effects models. Panels correspond to gear–season combinations: bag seine (BS; panels a–d: winter, spring, summer, fall), bay trawl (BT; panels e–h: winter, spring, summer, fall), and gillnet (GN; panels i–j: spring, fall). Colors distinguish thermal guilds: red = Subtropical, green = Temperate, blue = Tropical.

The most interpretively important contrast at the γ-diversity scale was between subtropical taxa in BS assemblages, which showed the largest positive slope of any thermal guild–gear combination (slope = 0.1304 yr⁻¹), and temperate taxa in BT assemblages, which showed the steepest decline (slope = −0.0814 yr⁻¹). Together, these results indicate that long-term increases in estuarine diversity are driven disproportionately by subtropical taxa in shallow habitats, consistent with progressive warm-affinity community reorganization, while temperate taxa experienced pronounced losses in deeper, trawl-sampled assemblages. Tropical taxa showed comparatively modest responses overall, suggesting either slower rates of expansion or greater sensitivity to habitat-specific constraints.

### 3.3 Spatial and temporal β-diversity across bays, gears, and functional guilds

#### Non-guild spatial and temporal β-diversity

Spatial and temporal β-diversity exhibited contrasting long-term patterns across the Texas coast. Multiplicative spatial β-diversity declined significantly through time when weighted toward common and dominant taxa (Shannon q = 1: slope = −0.0026 ± 0.0010 SE, p = 0.010; Dominance q = 2: slope = −0.0025 ± 0.0011 SE, p = 0.019; Fig. 6; Table S4), whereas richness-weighted spatial β-diversity showed no significant long-term change (q = 0; slope = 0.0024 ± 0.0012 SE, p = 0.052). Incidence-based Baselga metrics indicated modest increases in coastwide total spatial dissimilarity (βSOR: slope = 0.0003 ± 0.0001 SE, p = 0.025) and turnover (βSIM: slope = 0.0005 ± 0.0002 SE, p = 0.005) among bays, with no directional change in nestedness (βSNE: slope = −0.0001 ± 0.0001 SE, p = 0.330; Fig. 7; Table S4). Temporal β-diversity (Δyear = 1) showed no detectable long-term trend despite substantial interannual variability (slope = 0.0001 ± 0.0001 SE, p = 0.375; Fig. 8; Table S4). The apparently contradictory signals, incidence-based βSOR and βSIM increasing modestly while dominance-weighted βq₁ and βq₂ declined, are mechanistically coherent rather than contradictory. Incidence-based metrics are sensitive to rare and newly arriving species; if range-expanding subtropical taxa are appearing in some bays before others, incidence-based turnover increases because different bays temporarily host different newcomers. Dominance-weighted metrics, by contrast, track the abundant taxa that structure communities; as the same warm-adapted dominant species gain across all bays simultaneously, dominance-weighted β declines. Together these patterns indicate that long-term coastwide reorganization involves simultaneous divergence in the rare-species layer and convergence in the dominant-species layer, a signature consistent with ongoing tropicalization combined with coastwide homogenization in community dominance structure.

**Figure 6.**
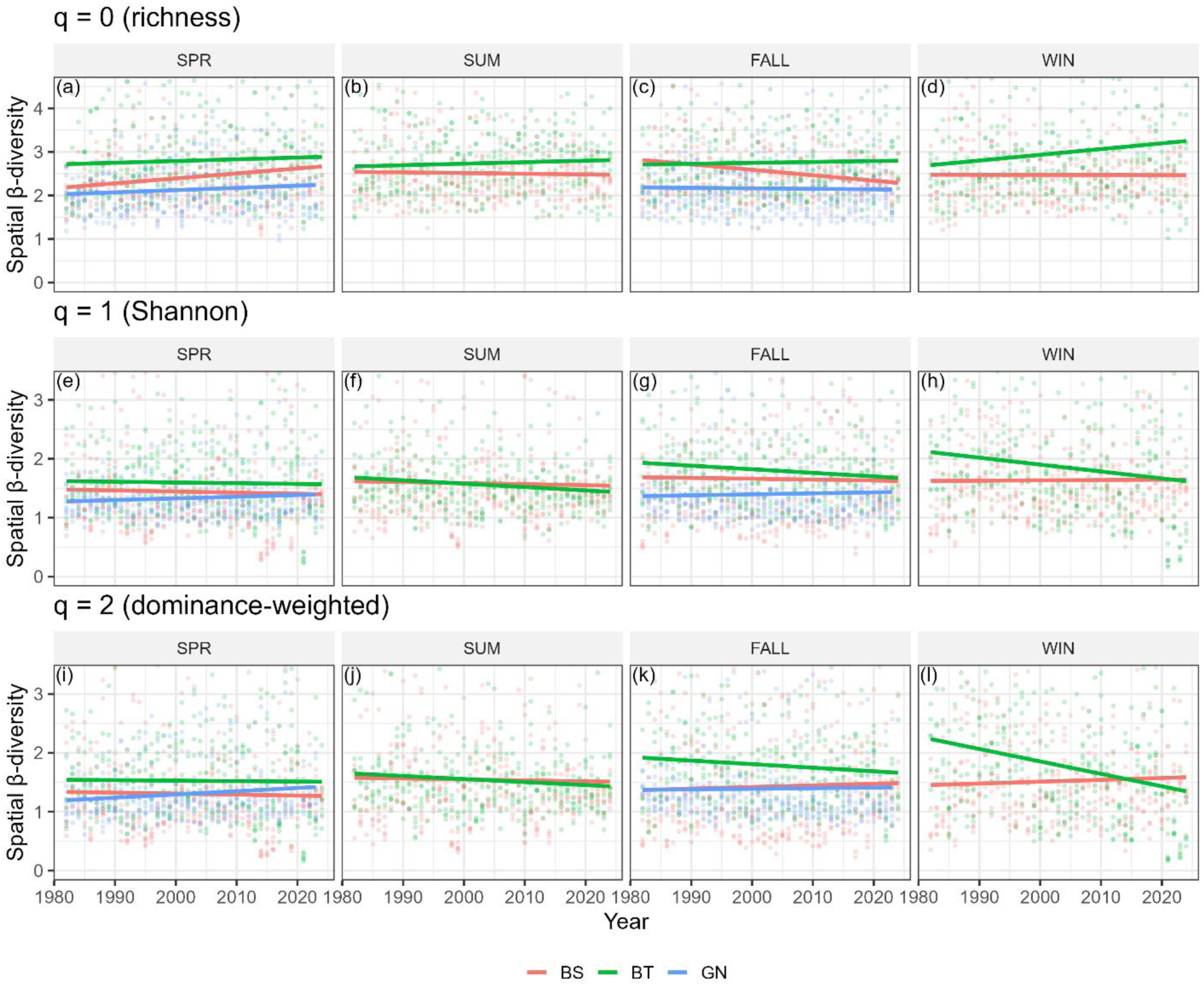
Long-term trends in spatial β-diversity among bays estimated using multiplicative partitioning of Hill numbers, 1982–2024. Points show annual estimates for each gear type, and solid lines show linear trends fitted separately for each gear. Panels are arranged with rows corresponding to diversity order, richness-weighted (q = 0; panels a–d), Shannon-weighted (q = 1; panels e–h), and dominance-weighted (q = 2; panels i–l), and columns corresponding to season (spring, summer, fall, winter). Colors distinguish sampling gear: red = bag seine (BS), blue = bay trawl (BT), green = gillnet (GN). Shannon-weighted (q = 1) and dominance-weighted (q = 2) spatial β-diversity declined significantly through time across most gears and seasons, indicating gradual spatial homogenization driven by convergence in dominant community structure rather than species loss. Richness-weighted spatial β-diversity (q = 0) showed no significant directional change, reflecting stability in the rare-species layer despite ongoing convergence in dominant taxa, a pattern consistent with simultaneous range expansion of newcomers and coastwide consolidation of warm-adapted dominants. Y-axis limits are truncated to central quantiles to improve visualization of long-term trends.

**Figure 7.**
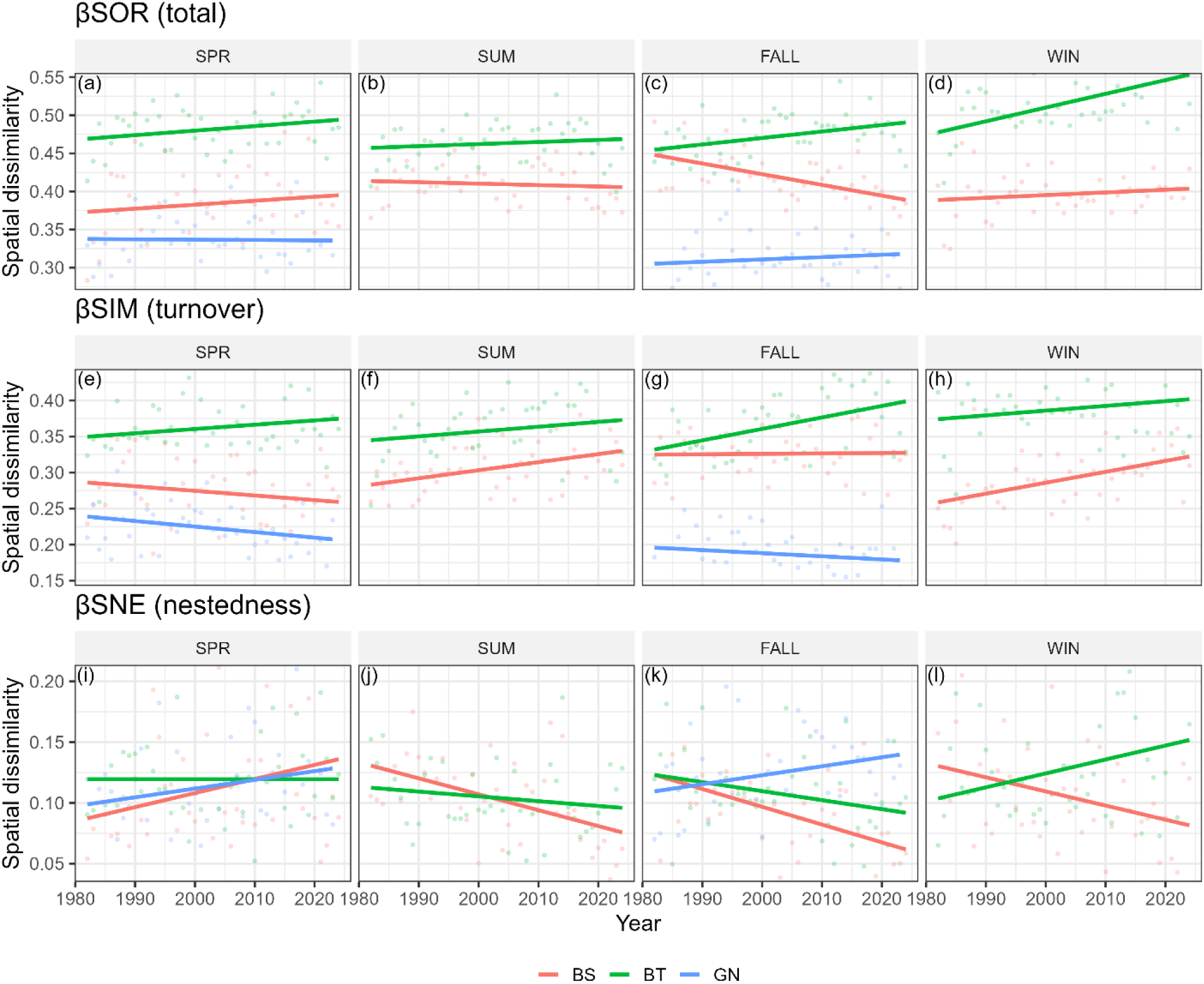
Decomposition of spatial β-diversity among bays into total dissimilarity (βSOR), turnover (βSIM), and nestedness (βSNE) using Baselga’s multi-site framework, 1982–2024. Points show annual mean values calculated across bays for each gear–season combination, and solid lines show linear trends fitted separately for each gear. Panels are arranged with rows corresponding to β-diversity component — total dissimilarity (βSOR; panels a–d), turnover (βSIM; panels e–h), and nestedness (βSNE; panels i–l) — and columns corresponding to season (spring, summer, fall, winter). Colors distinguish sampling gear: red = bag seine (BS), blue = bay trawl (BT), green = gillnet (GN). Total spatial dissimilarity and turnover increased modestly through time, while nestedness showed no directional change, indicating that spatial reorganization among bays proceeded primarily through species replacement rather than nested loss or gain. Y-axis limits are truncated to central quantiles to emphasize long-term trends.

**Figure 8.**
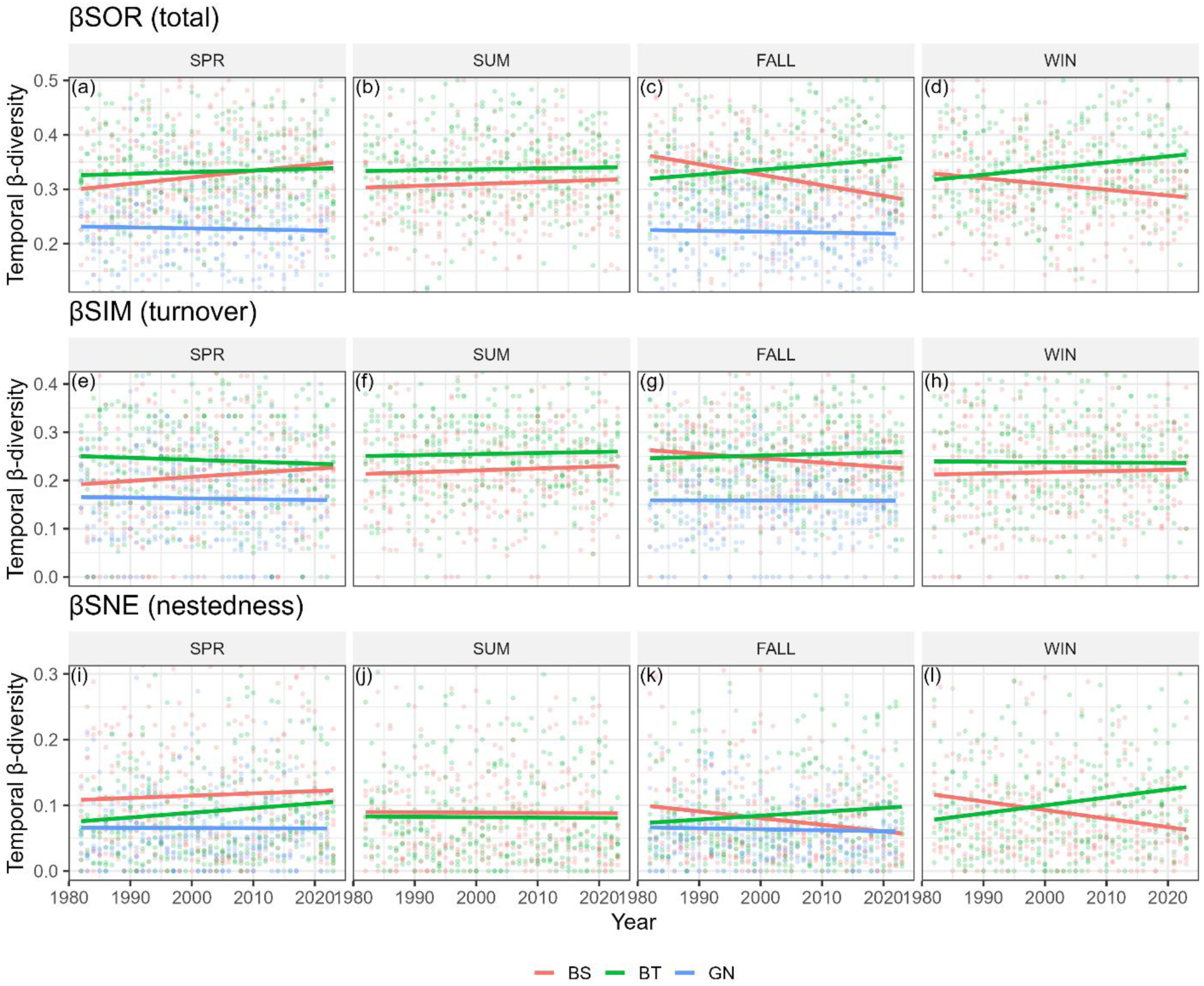
Temporal β-diversity between consecutive years (Δyear = 1) quantified using Baselga’s Sørensen dissimilarity (βSOR) and its turnover (βSIM) and nestedness (βSNE) components, 1982–2024. Points show annual dissimilarity values indexed by the starting year of each consecutive pair, summarized across bays, and solid lines show linear trends fitted separately for each gear. Panels are arranged with rows corresponding to β-diversity component — total dissimilarity (βSOR; panels a–d), turnover (βSIM; panels e–h), and nestedness (βSNE; panels i–l) — and columns corresponding to season (spring, summer, fall, winter). Colors distinguish sampling gear: red = bag seine (BS), blue = bay trawl (BT), green = gillnet (GN). Temporal β-diversity showed no consistent directional trend across gears or seasons, indicating that year-to-year compositional turnover remained stable over the monitoring period despite substantial long-term changes in α- and γ-diversity. Y-axis limits are truncated to central quantiles to improve visualization of long-term patterns.

#### Guild-structured spatial and temporal β-diversity

Guild-structured analyses revealed that dominance-weighted spatial homogenization was strongest for low-dispersal taxa and tropical taxa, which showed the steepest declines in spatial βq₂ across gears (Tables S5–S7; Figs. S7–S13). In contrast, high-dispersal taxa and subtropical taxa showed weak or no directional change in spatial β-diversity, indicating that homogenization was concentrated in taxa with limited movement capacity and those most sensitive to thermal change. These guild-specific patterns reinforce the interpretation that dominance-weighted homogenization reflects convergence in thermally filtered, habitat-resident taxa rather than a general coastwide process affecting all functional groups equally. Full guild-structured β-diversity results, including Baselga component decompositions, gear-specific patterns, and temporal β-diversity by guild, are provided in Supporting Information (Tables S5–S7; Figs. S7–S13).

### 3.4 Derivative-based detection of changes in diversity trends

Generalized additive models revealed pronounced non-linear temporal structure in α- and γ-diversity across gears and seasons, whereas β-diversity exhibited comparatively weak temporal variation. To clarify the nature of these dynamics, derivative-based diagnostics were used to distinguish between changes in **trend direction** (first-derivative sign changes) and changes in the **rate of change** (second-derivative behavior). Based on the joint behavior of first and second derivatives, three recurring cases were identified:

i. **accelerating increases**, in which diversity was increasing and the rate of increase intensified
ii. **accelerating declines**, in which diversity was decreasing and the rate of decline intensified
iii. **decelerating change**, in which diversity continued to increase or decrease but the rate of change weakened without reversing direction.

These cases provide a consistent framework for interpreting non-linear diversity trajectories across gears and diversity components.

Consistent with the GAM-based patterns described in Section 3.1, derivative analyses indicate that post-1990 increases in bag seine α- and γ-diversity primarily reflect periods of accelerating increases rather than constant linear growth, whereas bay trawl assemblages are characterized by accelerating declines or decelerating change around flat or declining trajectories without sustained positive shifts.

#### 3.4.1 Changes in trend direction (first-derivative diagnostics)

Confidence-supported changes in trend direction, defined here as statistically supported sign changes in the first derivative of the fitted smooths, where confidence intervals exclude zero, were common for α- and γ-diversity but were not detected for β-diversity (Table S8). These sign changes indicate reversals in long-term trajectories and delimit intervals within which acceleration cases can be interpreted.

**α-diversity** exhibited multiple confidence-supported sign changes across gears and seasons (Fig. 9). In bag seine samples, sign changes were detected only in the fall during the 2010s, marking transitions between periods of accelerating increases and subsequent decelerating change. Spring, summer, and winter series exhibited strong curvature but no confidence-supported reversals, indicating sustained increases punctuated by changes in rate rather than direction. Bay trawl α-diversity showed sign changes in all seasons, exhibiting one or more directional reversals between the mid-1990s and late 2010s (e.g., a mid-1990s reversal in fall, and late-2010s reversals in spring, fall, and winter). Gillnet α-diversity exhibited a single confidence-supported sign change in both spring and fall, occurring earlier than in other gears, primarily in the mid-1990s.

**Figure 9.**
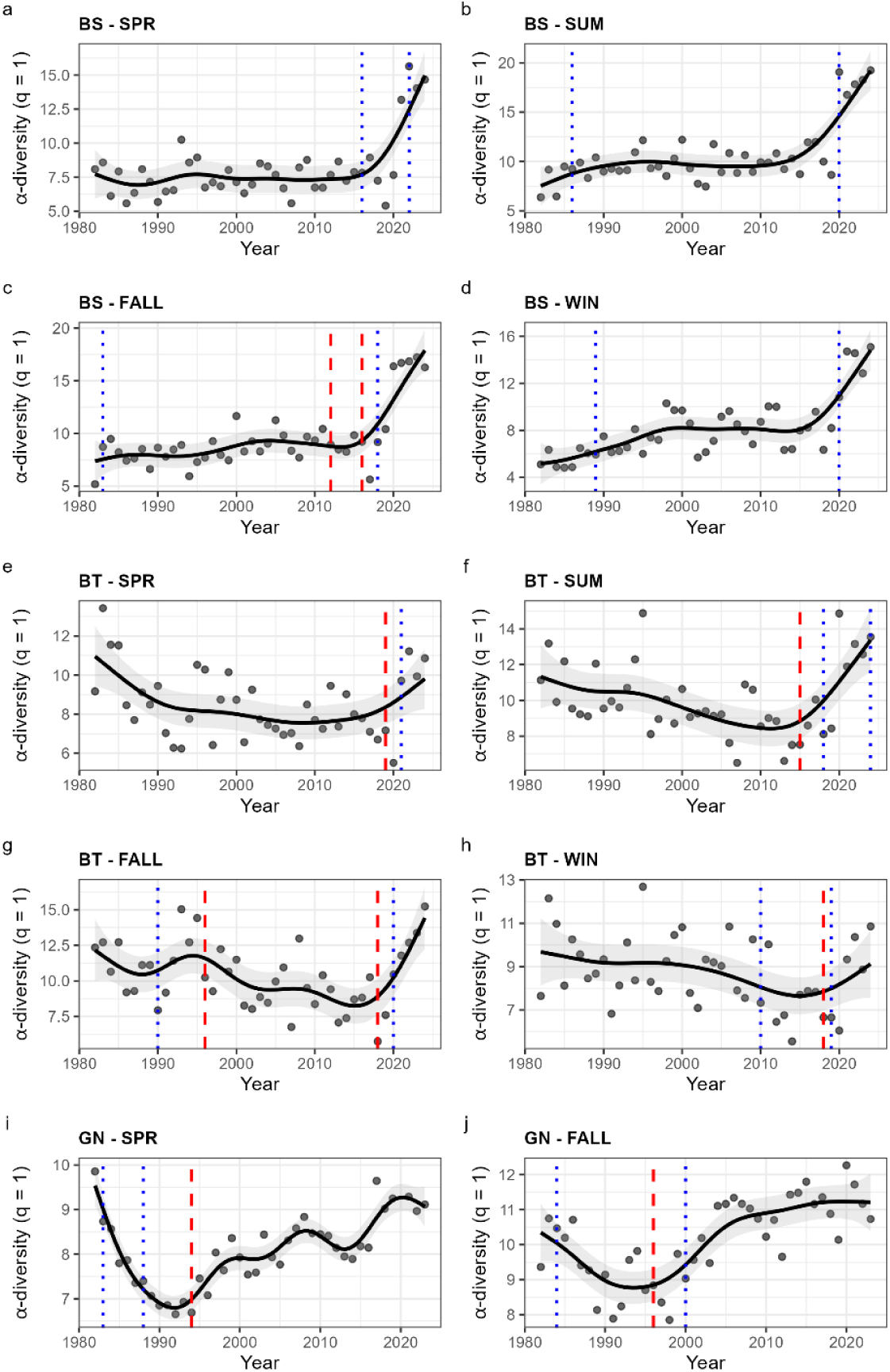
Long-term trajectories of local (α) diversity across gears and seasons, 1982–2024. Points show annual, coverage-standardized estimates of Shannon α-diversity (Hill number q = 1) averaged across bays, and solid lines show fitted generalized additive model (GAM) population smooths with shaded ribbons indicating 95% confidence intervals. Panels correspond to gear–season combinations: bag seine (BS; panels a–d: spring, summer, fall, winter), bay trawl (BT; panels e–h: spring, summer, fall, winter), and gillnet (GN; panels i–j: spring, fall). Vertical red dashed lines mark years in which the first derivative of the fitted smooth exhibited a confidence-supported sign change, indicating a statistically supported reversal in long-term trend direction. Vertical blue dotted lines mark peaks in the absolute value of the second derivative, indicating years of maximum acceleration or deceleration in the rate of diversity change. GAMs were fitted

**γ-diversity** showed even more complex directional dynamics (Fig. S15). Bag seine γ-diversity exhibited multiple sign changes across all seasons, indicating alternation between accelerating increases and decelerating change during long-term expansion of the regional species pool. Bay trawl γ-diversity displayed frequent sign changes in every season, reflecting repeated transitions among accelerating declines, decelerating change, and brief recoveries. Gillnet γ-diversity also exhibited confidence-supported sign changes in both spring and fall despite more limited seasonal coverage.

In contrast, **β-diversity** showed no confidence-supported sign changes in the first derivative for any gear or season (Fig. 10; Table S8), indicating an absence of directional reversals and limiting interpretation primarily to variation in the rate of change.

**Figure 10.**
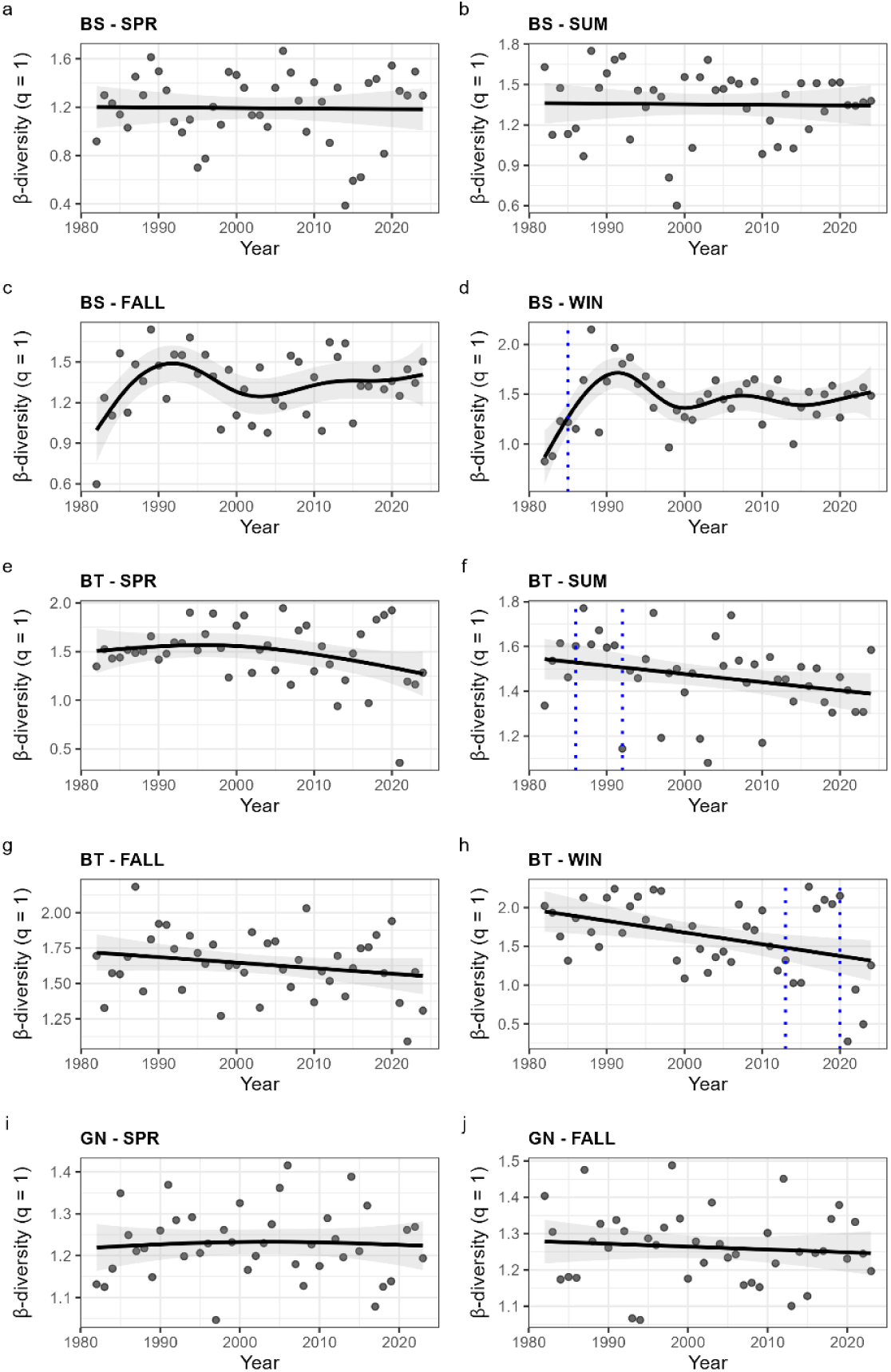
Long-term trajectories of spatial β-diversity (multiplicative β-diversity, q = 1) across gears and seasons, 1982–2024. Points show annual estimates averaged across bays, and solid lines show fitted generalized additive model (GAM) population smooths with shaded ribbons indicating 95% confidence intervals. Panels correspond to gear–season combinations: bag seine (BS; panels a–d: spring, summer, fall, winter), bay trawl (BT; panels e–h: spring, summer, fall, winter), and gillnet (GN; panels i–j: spring, fall). No confidence-supported reversals in trend direction were detected for β-diversity (i.e., no years in which the first-derivative confidence interval excluded zero with a sign change), and therefore no red dashed lines are shown. Vertical blue dotted lines mark peaks in the absolute value of the second derivative, indicating years of maximum acceleration or deceleration in the rate of spatial β-diversity change. GAMs were fitted

#### 3.4.2 Changes in the rate of change (second-derivative diagnostics)

Second-derivative diagnostics revealed widespread variation in the rate of change for α- and γ-diversity, including clear instances of accelerating increases, accelerating declines, and decelerating change (Table S8).

For **α-diversity**, bag seine samples exhibited pronounced acceleration peaks in the early portion of the time series (1980s) and again in the late 2010s to early 2020s, corresponding to periods of accelerating increases. Following these peaks, trajectories often transitioned into decelerating change without reversing direction. Bay trawl α-diversity showed repeated acceleration peaks across all seasons, frequently coinciding with accelerating declines during periods of long-term decrease or stagnation. Gillnet α-diversity exhibited multiple acceleration peaks despite fewer seasonal strata, indicating episodic intensification and subsequent slowing of change.

**γ-diversity** also exhibited strong second-derivative structure across gears and seasons. Acceleration peaks were common near the beginning and end of the time series and during intervals associated with first-derivative sign changes, indicating alternating phases of accelerating increases, accelerating declines, and decelerating change as regional diversity reorganized through time.

In contrast, **β-diversity** exhibited relatively few acceleration peaks, most notably in bay trawl summer and winter. These peaks did not coincide with directional reversals, indicating decelerating change or transient modulation of weak trends rather than sustained accelerating increases or declines.

#### 3.4.3 Synthesis

Together, first- and second-derivative diagnostics indicate that long-term biodiversity change along the Texas coast proceeds through episodic acceleration and deceleration rather than monotonic trends or sustained system-wide acceleration. α- and γ-diversity exhibit repeated transitions among accelerating increases, accelerating declines, and decelerating change, whereas β-diversity remains comparatively stable with limited temporal structure. This comparative stability in spatial turnover, alongside the tightly coupled trajectories of local and regional diversity, reflects a conserved multiscale structure where regional diversity scales strongly and positively with local diversity across all diversity orders (Fig. 11). These results confirm and refine the long-term patterns described in Section 3.1 by revealing how changes in diversity unfold through time rather than altering conclusions about net trend direction.

**Figure 11.**
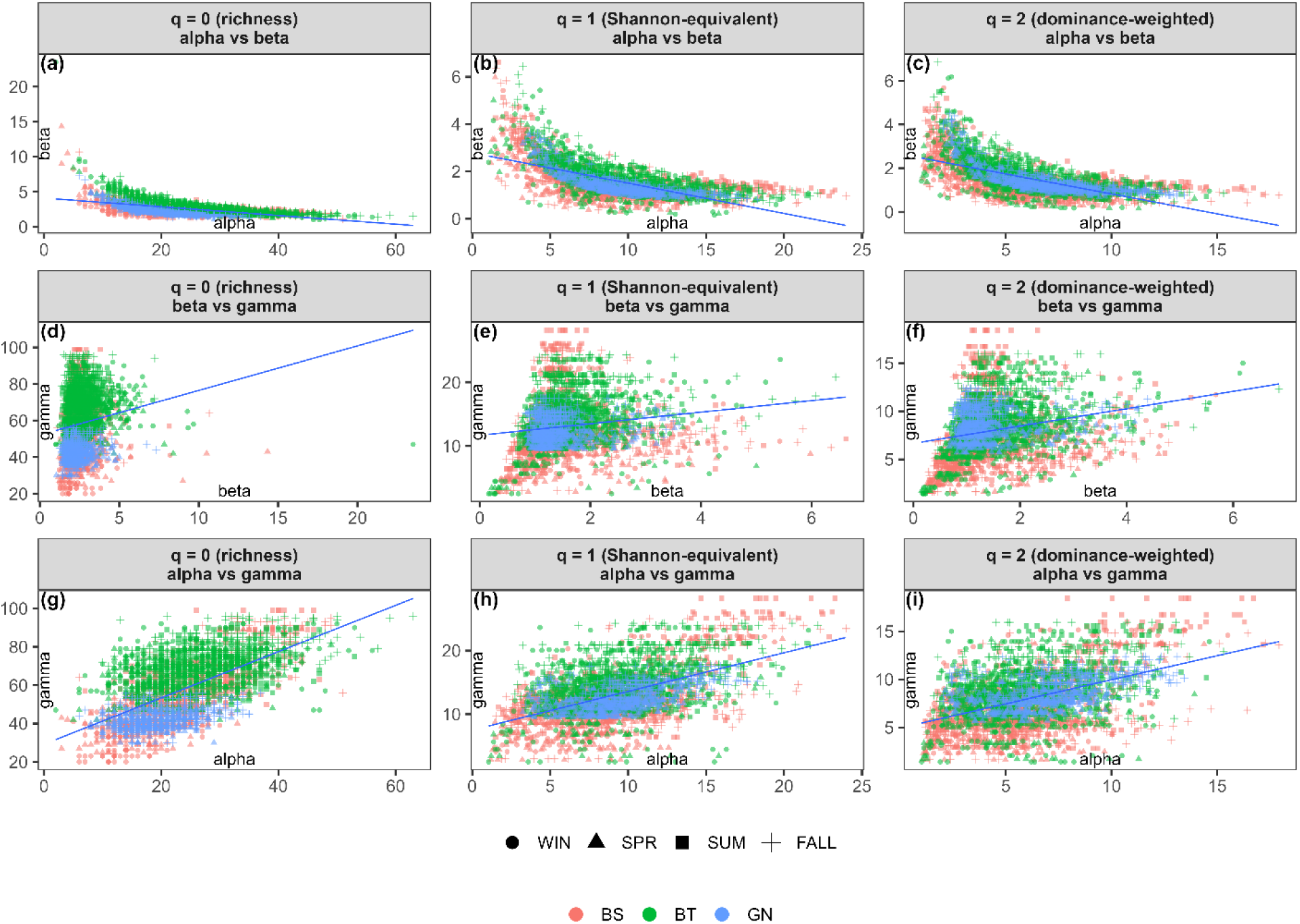
α–β, β–γ, and α–γ relationships across diversity orders. Scatterplots show pairwise relationships among local diversity (α), spatial turnover (β), and regional diversity (γ) for Hill numbers q = 0 (richness), q = 1 (Shannon-equivalent diversity), and q = 2 (dominance-weighted diversity). Rows correspond to diversity order (q), and columns show α vs β, β vs γ, and α vs γ relationships. Points represent bay-level observations for individual year × gear × season combinations, colored by sampling gear and distinguished by season using point symbols. Solid lines indicate ordinary least squares (OLS) smoothers fitted for visualization only. Axes are scaled independently across panels to highlight the shape and strength of relationships.

### 3.5 Chronic versus acute drivers of metacommunity diversity

To evaluate the relative importance of long-term environmental change, acute disturbance events, and gradual temporal trends, nested models were compared using ΔAIC within the best-supported acute-event structure for each response × gear combination identified in Stage 1 (see Section 2.13). Comparisons focused on the consequences of removing centered year, chronic environmental anomalies, acute event indicators, and spike metrics from the full model (Fig. 12; Table 4), with full Stage-2 model comparisons provided in Archive S1.

**Figure 12.**
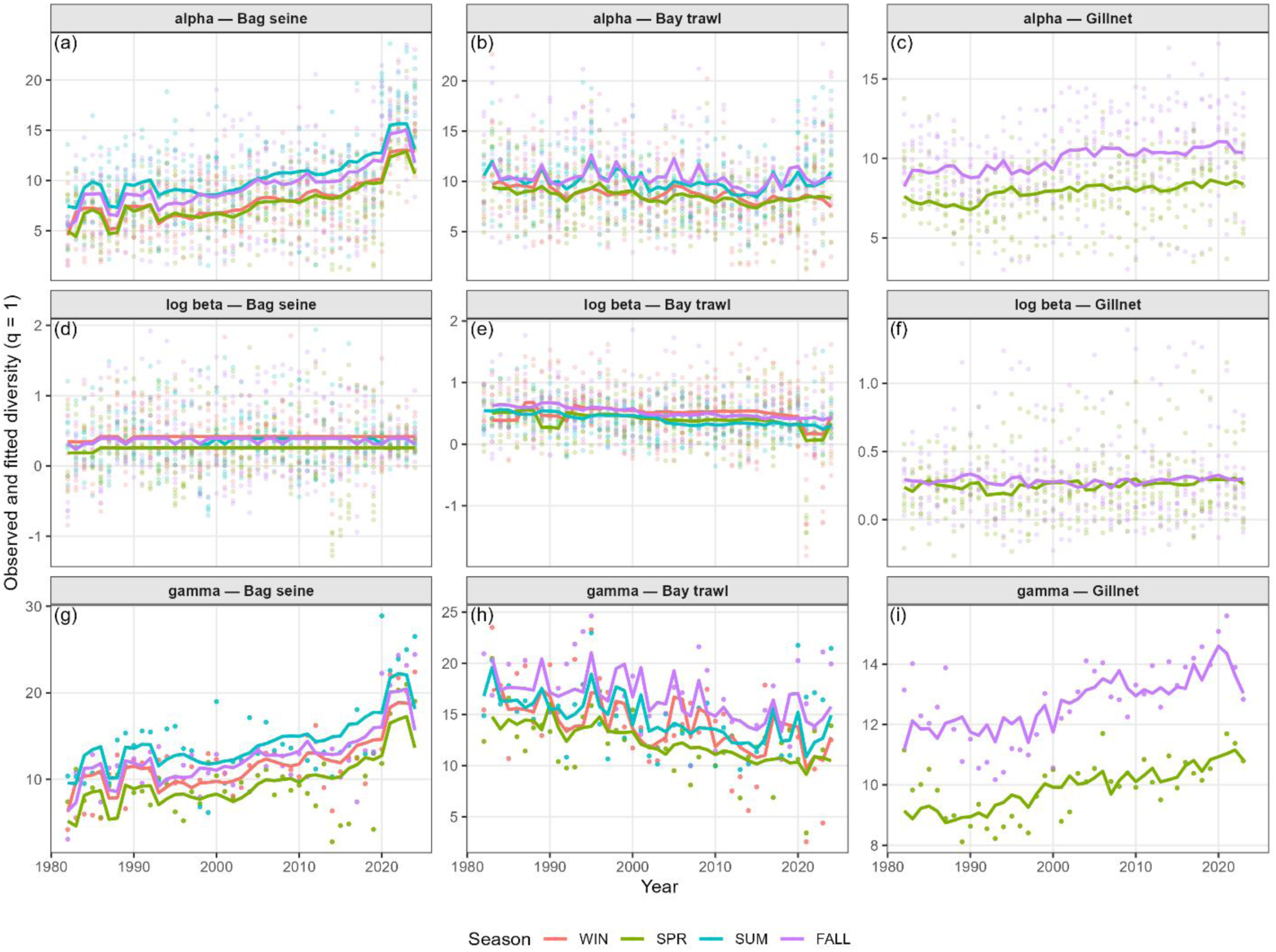
Observed and fitted temporal trajectories of metacommunity diversity. Observed annual diversity estimates (points) and fitted values (lines) from the best-supported chronic–acute models for α-, β-, and γ-diversity (q = 1), shown separately for bag seine, bay trawl, and gillnet data. For β-diversity, models were fitted and plotted on the log scale. Fitted lines represent predictions from the best model for each response × gear combination, selected by AIC in Stage 1, and incorporate chronic environmental anomalies, acute event terms, and a long-term year effect as supported. Colors indicate season. Panels (a–i) correspond to diversity component and gear combinations as labeled.

#### Relative importance of chronic, acute, and temporal effects

Across diversity components and sampling gears, models retaining both chronic environmental anomalies and the centered year term were consistently best supported (Fig. 12; Table 4). Removing the year effect resulted in large increases in AIC for nearly all α- and γ-diversity models, indicating strong support for gradual, long-term temporal change independent of measured environmental anomalies. Excluding chronic anomalies also reduced model support in most cases, though the associated ΔAIC values were generally smaller than those observed for removal of the year term.

In contrast, removing acute event indicators (freeze and storm exposure) produced substantially smaller changes in AIC and only rarely approached the magnitude of ΔAIC associated with chronic or temporal terms, regardless of how long acute events were encoded for in the models. Acute effects, therefore, contributed comparatively little to overall model support once long-term trends and chronic anomalies were accounted for.

These patterns were broadly consistent across gears and diversity components, although the magnitude of ΔAIC varied. Support for both year and chronic anomalies was strongest for α-and γ-diversity, whereas β-diversity exhibited weaker sensitivity to all driver classes, with smaller ΔAIC values across model reductions (Table 4).

#### Best-supported representations of acute events

The optimal representation of acute disturbances differed among responses and gears (Table 4; Archive S1). For α- and γ-diversity, models incorporating multi-month duration windows for freeze and storm exposure were most frequently supported, with longer windows (often 24–36 months) favored in several cases. In contrast, best-supported encodings for β-diversity varied more strongly by gear and included both short duration windows and seasonally repeated (0–2 yr) representations.

Despite this heterogeneity in acute-event encoding, the relative dominance of chronic anomalies and the year effect remained unchanged across alternative representations of acute exposure. Full Stage-1 and Stage-2 AIC comparisons demonstrate that conclusions regarding the primacy of long-term drivers are robust to how acute disturbances are parameterized (Archive S1).

#### Variance explained

Best-supported models explained a moderate fraction of variance in α- and γ-diversity, with marginal *R*^2^ values indicating meaningful contributions from fixed effects and conditional *R*^2^ values (where applicable) reflecting additional spatial structure captured by bay-level random intercepts (Table 4). In contrast, β-diversity models consistently exhibited low marginal *R*^2^ across gears, even after log transformation and in best-supported models, indicating that spatial turnover was only weakly associated with the chronic and acute environmental predictors considered here.

#### Direction and form of chronic and acute effects on α-diversity

Observed and fitted trajectories of α-diversity illustrate the contrasting influences of chronic forcing and acute disturbances (Fig. 12). Chronic temperature and salinity anomalies were associated with smooth, directional changes in α-diversity over time, broadly consistent across seasons within a given gear. In contrast, effects of acute freeze and storm exposure were generally weaker and more variable, with limited and gear-specific departures from long-term trends. These visual patterns reinforce the model comparison results by highlighting the persistent and predictable influence of chronic environmental change and gradual temporal trends, relative to the transient effects of discrete disturbance events.

## 4. Discussion

Understanding how biodiversity reorganizes under sustained environmental change requires a framework that links processes operating across spatial scales, functional traits, and temporal persistence (Levin 1992; Loreau et al. 2001; McGill et al. 2006). Biodiversity change is rarely expressed as uniform gain or loss; instead, it often emerges as habitat-specific restructuring shaped by the interplay of chronic environmental forcing, episodic disturbance, and trait-mediated responses that govern persistence, dispersal, and dominance (Turner 2010; Magurran et al. 2010; Blowes et al. 2019). A multiscale perspective that integrates α-, β-, and γ-diversity is therefore essential for distinguishing changes driven by local community dynamics from those arising through spatial turnover or regional species-pool restructuring (Whittaker 1960, 1972; Leibold et al. 2004). Central to this perspective is Levin’s (1992) insight that ecological patterns exhibit coherence only within characteristic spatial domains, a concept with direct implications for identifying the scales at which biodiversity reorganization is predictable. Viewed through this framework, ecological change can proceed as accelerating reorganization within a conserved scaling structure, revealing how communities adapt and redistribute without necessarily destabilizing broader spatial patterns (Dornelas et al. 2014; McCann 2000; Thibaut and Connolly 2013). Here, we report three findings that extend these frameworks in ways not anticipated by existing theory: that habitat context overrides trait-based predictions of reorganization direction; that spatial homogenization proceeds undetected by richness-based β-diversity metrics but is revealed through dominance-weighted measures, operating through distinct gain-driven and loss-driven mechanisms across habitat domains; and that reorganization has intensified in pace while fundamental α–β–γ scaling relationships have been preserved.

### 4.1. Diversity change is persistent, directional, and habitat dependent

Long-term changes in diversity along the Texas coast were persistent and directional, but differed systematically among estuarine habitats, as reflected by gear-specific trajectories of α- and γ-diversity (Figs. 2–3; Table 3). In shallow habitats sampled by bag seines (BS) and gillnets (GN), α-diversity increased steadily over four decades across seasons, rising from approximately 6–8 effective species in the early 1980s to more than 12–15 by the late 2010s (Fig. 2; Figs. S1–S2). Corresponding increases in γ-diversity, including a near doubling of regional diversity in BS assemblages (Fig. 3; Figs. S3–S4), reflect a broadening contribution of species to the assemblage, through reduced dominance and/or more consistent occurrence, rather than increased differentiation among bays. Together, these parallel trends indicate sustained accumulation and redistribution of diversity within shallow estuarine habitats rather than transient responses to short-term environmental variability or disturbance. These patterns are consistent with prior evidence from the same monitoring program that life-history traits structure which species increase or decrease under sustained environmental change, with species common during summer months, those better adapted to warmer conditions, disproportionately driving observed increases in incidence over time (Fujiwara et al. 2022).

**Table 3.** Synthesis of diversity change and spatial homogenization across functional guilds and habitats.

| Functional dimension | Gear | $\alpha$ trend | $\gamma$ trend | Spatial $\beta$ (q = 2) | Mechanism | Outcome |
| --- | --- | --- | --- | --- | --- | --- |
| **Dispersal** | BS | ↑ (all) | ↑ (all) | ↓ (low) | Dominance convergence | Winners |
|  | GN | ↑ (int.) | ↑ (int. > high) | ↓ (low) | Selective convergence | Winners |
|  | BT | ↓ (low, int.) | ↓ (low) | ↓↓ | Loss of distinctive taxa | Losers |
| **Thermal** | BS | ↑ (subtrop.) | ↑ (subtrop.) | ~ | Limited homogenization | Winners |
|  | GN | ↑ (subtrop.) | ↑ (subtrop.) | ↓ (temp.) | Dominance convergence | Winners |
|  | BT | ↓ (temp.) | ↓ (temp.) | ↓↓ | Loss-driven homogenization | Losers |

In contrast, bay trawl (BT) assemblages exhibited flat or declining trends in both α- and γ-diversity (Figs. 2–3; Figs. S1–S4), indicating that diversity gains in shallow habitats were not shared across the estuarine depth gradient. This is consistent with the recent finding of Williford and Anderson (2025). This habitat-dependent divergence supports the view that biodiversity change proceeds through spatially structured reorganization rather than uniform gain or loss of taxa (McGill et al. 2015; Blowes et al. 2019, 2024). By documenting sustained increases in shallow habitats alongside stagnation or decline in deeper systems, these results extend prior Texas coast studies (Fujiwara et al. 2019; Pawluk et al. 2021) and demonstrate that depth-linked habitat context constrains how local and regional diversity respond to shared environmental forcing, consistent with scale-dependent ecological processes (Levin 1992). Deeper soft-bottom habitats sampled by bay trawls may be particularly vulnerable to cumulative anthropogenic pressures, including bottom-contact fisheries, habitat alteration, and water-quality degradation, which could limit long-term diversity accumulation.

Several non-exclusive mechanisms may underlie this habitat-specific vulnerability. Long-term increases in the frequency and duration of hypoxic conditions in deeper bay waters, linked to warming and stratification, could disproportionately constrain demersal assemblages with limited vertical refuge (Bugica et al. 2020). Variability in freshwater inflow associated with intensifying precipitation extremes may additionally alter sediment composition and benthic habitat quality in ways that differentially affect soft-bottom communities, particularly taxa with low dispersal capacity and limited ability to track shifting habitat conditions (Bugica et al. 2020). Resolving the relative contributions of these mechanisms would require targeted analyses linking long-term environmental trajectories in deeper bay waters to the specific taxa responsible for declining BT diversity, a direction that merits explicit attention in future work.

These habitat-specific α- and γ-diversity trajectories cannot be attributed to differences in functional composition, as dispersal and thermal guilds were broadly similar across BS, GN, and BT assemblages (Table 2). Instead, divergence reflects habitat-dependent responses of comparable functional guilds, with several dispersal and thermal groups exhibiting negative diversity trends in BT habitats despite similar guild availability (Figs. 4–5; Tables S2–S3). Thus, functional traits modulate the magnitude of diversity change within a habitat domain, but habitat context determines its direction; the same guild can increase in shallow assemblages while declining in deeper ones. This result challenges purely trait-based forecasting frameworks, which implicitly assume that trait identity predicts reorganization direction independently of local habitat setting.

Finally, the strong parallelism between α- and γ-diversity trends within each gear type indicates that regional biodiversity change emerged primarily from parallel local responses to shared environmental conditions. Consistent with this interpretation, empirical scaling relationships demonstrate a strong, consistent positive association between local diversity (α) and regional diversity (γ) across all diversity orders, indicating that regional gains are tightly coupled to local changes (Fig. 12). Increases in γ, therefore, largely tracked local gains in α rather than increased spatial turnover among bays. Together with weak and nuanced temporal responses of β-diversity (Figs. 6–8), these patterns indicate that long-term regional biodiversity change reflects coordinated within-habitat responses to shared environmental conditions across bays rather than divergent regional trajectories.

### 4.2. Long-term processes dominate over acute disturbances

Having established that biodiversity reorganization is habitat-contingent and detectable through dominance-weighted metrics, we next ask whether long-term temporal structure or episodic disturbance better explains the observed patterns. Although acute events can generate abrupt demographic responses, their effects are often transient unless they permanently alter environmental baselines or push systems across critical thresholds (Scheffer et al. 2001). Uncertainty remains over whether long-term biodiversity patterns primarily reflect cumulative chronic forcing or episodic disturbance (Turner 2010; Smith 2011; Jentsch et al. 2007). Our results directly address this uncertainty by showing that gradual temporal change and chronic environmental anomalies were far more strongly associated with multiscale diversity patterns along the Texas coast than acute disturbances, whereas acute disturbances played a comparatively minor role (Table 4; Archive S1). For both α- and γ-diversity, removal of the centered year term resulted in substantially larger increases in AIC than exclusion of acute disturbance terms (Table 4), indicating that long-term temporal structure explained far more variation in diversity than discrete freeze or storm events, even in these disturbance-prone estuarine systems.

**Table 4.** Relative importance of driver classes based on ΔAIC. ΔAIC values represent the increase in AIC when a given driver class (YEAR, chronic anomalies, acute events, or spikes) is removed from the best-supported model for each response × gear combination after Stage 1 model selection. Larger ΔAIC values indicate stronger support for that driver class. ΔAIC values for spike terms are reported only for models in which spike metrics were retained during Stage 1 model selection; cells are blank where spikes were not included in the best-supported model.

| Response<br>(q = 1) | Gear | Event<br>encoding | Window | Spikes | n | AIC<br>(full) | $\Delta AIC$:<br>Year | $\Delta AIC$:<br>Chronic | $\Delta AIC$:<br>Acute | $\Delta AIC$:<br>Spikes |
| --- | --- | --- | --- | --- | --- | --- | --- | --- | --- | --- |
| log $\beta$ | BS | Duration | 3 m | No | 1360 | 1318.1 | -2.0 | -4.0 | 1.7 | — |
| log $\beta$ | BT | Repeat3mo | 0–2 yr | No | 1360 | 1291.9 | 18.7 | 14.5 | 46.3 | — |
| log $\beta$ | GN | Duration | 24 m | Yes | 656 | -462.4 | 2.2 | 9.7 | 5.1 | 0.3 |
| $\alpha$ | BS | Duration | 36 m | Yes | 1360 | 6951.3 | 341.2 | 26.6 | 116.7 | 2.8 |
| $\alpha$ | BT | Duration | 3 m | Yes | 1360 | 6959.5 | 0.9 | 26.7 | 26.7 | 14.1 |
| $\alpha$ | GN | Repeat3mo | 0–2 yr | Yes | 656 | 2413.7 | 27.5 | 21.4 | 2.9 | 4.1 |
| $\gamma$ | BS | Duration | 36 m | Yes | 1360 | 7050.7 | 569.9 | 69.3 | 240.4 | 18.2 |
| $\gamma$ | BT | Duration | 6 m | Yes | 1360 | 6811.7 | 123.5 | 37.9 | 183.4 | 39.7 |
| $\gamma$ | GN | Duration | 9 m | Yes | 656 | 1686.0 | 162.7 | 41.5 | 4.1 | 76.9 |

This pattern aligns with broader empirical evidence demonstrating that biodiversity responses to climate change are frequently governed by cumulative environmental exposure rather than by the immediate effects of short-term extreme events (Parmesan and Yohe 2003; IPCC 2021). While acute disturbances may leave strong short-term signatures, their influence on decadal diversity trajectories and evolutionary timescales is often attenuated by recovery, dispersal, and compensatory dynamics (Jentsch et al. 2007; Smith 2011).

This conclusion, however, should be interpreted as applying to coastwide, community-level diversity rather than to fine-scale patch dynamics. Storm impacts, particularly wind-driven surge, localized hypoxia, and freshwater pulse effects, are spatially heterogeneous at scales finer than the monitoring grid, which is structured around stratified random sampling within fixed 1-minute latitude × longitude cells. Standardized fixed-effort sampling of this kind is well suited to detecting coastwide trends but may average over localized post-disturbance signals that are spatially restricted and short-lived. As a result, AIC-based comparisons of chronic and acute driver classes may systematically underestimate the importance of acute disturbances at fine spatial scales, particularly in the immediate aftermath of major storm events. The relative dominance of chronic forcing demonstrated here therefore reflects the scale at which this monitoring program operates, and future work combining fine-scale post-disturbance surveys with long-term monitoring data would help clarify the degree to which episodic events shape biodiversity dynamics at patch scales.

### 4.3. Change is intensifying through time, but without destabilization

One of the unexpected results of this study is that biodiversity change has intensified in recent decades, expressed as episodic acceleration in α- and γ-diversity rather than abrupt shifts in trend direction. Periods of accelerated diversity change became most frequent after approximately 2000 (Fig. 9; Fig. S14; Table S8). Importantly, this intensification was largely confined to α- and γ-diversity, whereas β-diversity exhibited weak temporal structure, with no clear sign changes in temporal trends and only limited evidence of acceleration (Fig. S15; Table S8). The decoupling of accelerated local and regional change from stable spatial turnover indicates that recent biodiversity change reflects faster reorganization within communities and regional species pools, rather than increased volatility or breakdown of spatial structure among bays. This finding also has practical implications for ecological monitoring: acceleration in the pace of change need not trigger alarm about structural breakdown if the metacommunity’s scaling relationships remain intact, but it does signal that the window for detecting and responding to reorganization is narrowing.

### 4.4. Local processes dominate regional biodiversity outcomes

Metacommunity theory predicts that local, spatial, and regional components of diversity are linked through stable scaling relationships, even as species identities and abundances change through time (Whittaker 1960, 1972; Leibold et al. 2004). Consistent with these predictions, α–β–γ relationships along the Texas coast were largely preserved despite pronounced long-term changes in diversity magnitude and community composition, indicating that multiscale diversity structure can persist under sustained ecological reorganization. Empirical scaling relationships indicate that variation in regional diversity scales strongly and positively with local diversity across all Hill orders (Fig. 12), demonstrating that regional diversity change emerged from parallel shifts within local communities across bays rather than from increasing spatial differentiation among them. This coupling is most plausibly explained by a shared coastwide environmental signal, particularly the long-term warming trend documented along the Texas coast (Tolan and Fisher 2009; Wang et al. 2023), driving parallel responses within local communities across all bays simultaneously. Consistent with this interpretation, the strongest diversity gains were concentrated in subtropical and low-to-intermediate dispersal taxa (Section 4.6), groups whose responses reflect thermal filtering rather than active redistribution among bays. Had dispersal-mediated redistribution been the primary mechanism, high-dispersal taxa would be expected to show the strongest and most synchronized responses across bays, the opposite of what was observed.

The persistence of α–β–γ relationships across richness (q = 0), Shannon-equivalent diversity (q = 1), and dominance-weighted diversity (q = 2), together with robustness to moderate nonlinearity (Table S8), indicates that the fundamental structure of the Texas estuarine metacommunity has been conserved despite substantial shifts in diversity magnitude and composition. These results demonstrate that biodiversity reorganization occurred within a conserved multiscale framework rather than through breakdown of diversity integration across scales. Although the relative contributions of diversity components shifted through time, the structural relationships linking them persisted, indicating resilience of metacommunity structure under sustained environmental change, a property that emerges here from parallel, habitat-specific responses of local communities to shared environmental conditions, rather than from stability in spatial turnover or from dispersal-mediated redistribution among bays. This reorganization was habitat dependent (Figs. 2–3), trait structured (Figs. 4–5; Table 3), and followed a chronic rather than episodic pattern (Table 4), while fundamental scaling relationships were maintained (Fig. 11). Formally distinguishing between a shared-driver mechanism and dispersal-mediated coupling would require experimental or model-based approaches that isolate environmental synchrony from organismal connectivity, a direction that merits explicit investigation in future work.

These results also speak directly to Levin’s (1992) concept of domains of scale, the idea that ecological patterns exhibit coherence only within characteristic spatial domains, and that relationships observed at one scale need not hold at another. Our findings suggest that the Texas estuarine metacommunity is organized into two nested scale domains. Within the horizontal domain, among bays spanning the coastline, diversity change is coherent, predictable, and synchronized, with all bays responding in parallel to shared environmental conditions regardless of their geographic position. Within the habitat domain, across the depth gradient from shallow nearshore to deeper benthic habitats, diversity trajectories diverge sharply, with shallow assemblages gaining and deeper assemblages losing diversity under the same regional conditions. The horizontal domain therefore acts as a single integrated unit in which local and regional diversity are tightly coupled, whereas the habitat domain defines the axis along which ecological differentiation is expressed. This interpretation suggests that the relevant scale domain for predicting biodiversity reorganization in Texas estuaries is not horizontal geographic space but rather the habitat gradient, and that forecasting frameworks must be parameterized within, not across, habitat domains to produce reliable predictions.

### 4.5. Homogenization reflects convergence in dominance structure

Both theory and empirical work indicate that biodiversity reorganization can proceed through decoupled changes in local, spatial, and temporal components, such that strong trends in α- and γ-diversity need not be accompanied by pronounced changes in spatial turnover (Levin 1992; Magurran et al. 2010). For all sampling conducted along the Texas coast, log-transformed β-diversity exhibited weak temporal trends across all gears (Figs. 6–8), contrasting sharply with strong directional increases in α- and γ-diversity (Figs. 2–3). In bag seine and gillnet assemblages, fitted log β remained near constant throughout the time series (Fig. 8), indicating relatively stable multiplicative spatial turnover despite substantial local and regional change.

The modest increases in incidence-based spatial dissimilarity alongside declining dominance-weighted β-diversity are consistent with a dual process: range-expanding subtropical taxa appearing in some bays before others temporarily increases species-level turnover in the rare-species layer, while the same warm-adapted dominant taxa consolidating across all bays simultaneously drives dominance-weighted convergence. This dual signal, divergence in rare newcomers, convergence in dominant residents, is a mechanistically coherent signature of tropicalization combined with homogenization, and one that would be invisible to studies relying on either metric alone.

This interpretation is reinforced by the weak sensitivity of β-diversity to environmental forcing. Removal of year or chronic anomaly terms resulted in small or negative ΔAIC values (Table 4), marginal R² values were consistently low (Archive S1), and year-to-year temporal β-diversity showed no consistent directional trend (Fig. 8; Figs. S12–S13). Together, these patterns indicate that spatial and temporal dimensions of biodiversity change operate along distinct axes, and that short-term variability can obscure slow, spatially expressed reorganization when inference relies solely on interannual turnover metrics (Levin 1992; Magurran et al. 2010). This decoupling between richness-based and dominance-weighted β-diversity metrics has a direct monitoring implication: studies that rely on conventional incidence-based or richness-weighted turnover measures will systematically fail to detect homogenization that is already underway through convergence in abundant taxa, producing false assurance of spatial stability.

Despite the temporal stability of conventional β-diversity metrics, dominance-weighted spatial β-diversity declined through time (q = 2), revealing gradual spatial homogenization driven by convergence in dominant community structure rather than widespread species loss or accelerating turnover (Fig. 6; Figs. S7–S9). This dominance-weighted decline is consistent with biotic homogenization arising from convergence in dominant taxa (Olden and Rooney 2006; Magurran and Henderson 2010; McGill et al. 2007) and demonstrates the limitations of richness-based β-diversity metrics. However, the mechanism underlying this convergence differed between habitat domains. In shallow assemblages sampled by bag seines and gillnets, where α-and γ-diversity increased, homogenization reflected gain-driven convergence, the same warm-adapted dominant taxa increasing simultaneously across all bays. In deeper assemblages sampled by bay trawls, where α- and γ-diversity declined, homogenization instead reflected loss-driven convergence, a narrowing of the dominant assemblage as taxa were lost, producing increasingly similar impoverished communities across bays. Both processes generate declining dominance-weighted β-diversity and both are invisible to richness-based metrics, but they have fundamentally different implications for ecosystem function and resilience: gain-driven convergence reflects community reorganization under sustained environmental change, whereas loss-driven convergence signals progressive erosion of the dominant assemblage in habitat-constrained communities with limited compensatory capacity.

### 4.6. Functional traits and habitat influence biodiversity

Functional ecology predicts that species traits mediate exposure and response to environmental change, shaping how communities reorganize even when taxonomic outcomes are complex (Lavorel and Garnier 2002; McGill et al. 2006). In coastal systems, dispersal capacity and thermal affinity are particularly relevant because they influence persistence and redistribution within spatially heterogeneous, hydrodynamically connected habitats (Winemiller and Rose 1992; Pinsky et al. 2013). Consistent with these expectations, trait-based analyses revealed clear differences in the direction and magnitude of biodiversity change among dispersal and thermal guilds (Figs. 4–5; Table 3). However, these traits did not explain divergence among habitats, which instead reflected habitat-specific constraints acting on otherwise comparable functional guilds. This pattern extends trait-based frameworks by demonstrating that species-level dispersal capacity and thermal affinity shape community-level biodiversity responses, but also that their expression is strongly influenced by habitats. Habitat context constrains how these trait-mediated responses scale across α-, β-, and γ-diversity, an integration rarely tested in long-term empirical systems.

In shallow habitats sampled by bag seines and gillnets, increases in α- and γ-diversity were most pronounced for intermediate- and low-dispersal guilds (Fig. S5; Table S2) and coincided with weak or declining dominance-weighted β-diversity (Figs. 6, S7–S9), suggesting that the same taxa gained simultaneously across bays in response to a shared environmental signal rather than through active dispersal-mediated redistribution. In contrast, bay trawl assemblages exhibited negative α- and γ-diversity trends across multiple dispersal guilds, particularly among low-dispersal taxa (Fig. S5; Table S2), indicating limited compensatory capacity within deeper soft-bottom habitats. Importantly, these patterns emerged despite broadly similar representation of dispersal guilds across gears (Table 2), reinforcing that dispersal traits influenced how taxa responded to long-term change rather than which traits were present.

Thermal affiliation further modulated biodiversity responses. Subtropical taxa exhibited consistently positive α- and γ-diversity trends in shallow habitats, whereas temperate taxa declined most strongly in bay trawl assemblages (Fig. S6; Table S3). These contrasting responses align with expectations under sustained warming, which preferentially favors warm-affinity species while constraining temperate taxa near their lower-latitude range limits (Sunday et al. 2012; Pinsky et al. 2013). Together with strong support for chronic environmental anomalies in α- and γ-diversity models (Table 4), these results indicate that functional traits modulate habitat-specific biodiversity responses to long-term environmental change without fundamentally restructuring functional composition across habitats. These thermal guild responses also provide the most direct evidence for the mechanism underlying local–regional diversity coupling: because warming operates coastwide and simultaneously across all bays, subtropical taxa gain and temperate taxa decline in parallel within each bay, producing the tight α–γ scaling relationship observed without requiring dispersal-mediated redistribution among bays.

## 5. Conclusions

In conclusion, we found that habitat context determined the direction of trait-mediated biodiversity reorganization, with the same dispersal and thermal guilds exhibiting opposite diversity trajectories across gear-defined habitat domains despite broadly similar functional composition, demonstrating that trait-based forecasting frameworks will mispredict reorganization direction when habitat context is ignored. Spatial homogenization proceeded undetected by richness-based β-diversity metrics but was revealed through dominance-weighted measures, operating through gain-driven convergence on shared warm-adapted dominants in shallow assemblages and loss-driven convergence toward impoverished communities in deeper habitats, both processes invisible to conventional richness-based monitoring, but with fundamentally different implications for ecosystem resilience. Despite a measurable intensification in the pace of local and regional diversity change after approximately 2000, spatial turnover among bays remained stable and α–β–γ scaling relationships were preserved, demonstrating that rapid reorganization can proceed without destabilizing fundamental metacommunity structure. These patterns were embedded within a broader context of persistent, directional, and habitat-dependent diversity change following a chronic rather than episodic trajectory, consistent with sustained environmental processes as the primary driver. Together, these findings advance understanding of how metacommunities reorganize under sustained change by revealing that resilience can coexist with acceleration, that habitat context constrains the expression of trait-mediated responses, and that detecting ongoing reorganization requires diversity metrics sensitive to changes in dominance structure rather than species presence alone.

## Acknowledgments

This work was conducted while M.F. was on sabbatical leave from Texas A&M University.

## Supporting Information

Supporting Information is available online for this article.

**Document S1. Supplementary Tables (PDF).**

A compiled PDF containing supplementary tables reporting detailed model outputs, parameter estimates, and statistical summaries supporting the main analyses.

**Document S2. Supplementary Figures (PDF).**

A compiled PDF containing supplementary figures illustrating spatial and temporal patterns of α-, β-, and γ-diversity across gears, bays, and functional guilds.

**Archive S1. Model comparison tables (ZIP).**

A compressed archive containing comma-separated value (CSV) files with full AIC-based model comparison results for log-transformed β-diversity. These files provide complete candidate model AIC values by gear and for the unified Stage 2 analysis, serving as an archive to support transparency and reproducibility.

## References

Baselga, A. 2010. Partitioning the turnover and nestedness components of beta diversity. Global Ecology and Biogeography 19:134–143.

Baselga, A., and C. D. L. Orme. 2012. betapart: an R package for the study of beta diversity. Methods in Ecology and Evolution 3:808–812.

Bates, D., M. Mächler, B. Bolker, and S. Walker. 2015. Fitting linear mixed-effects models using lme4. Journal of Statistical Software 67.

Benjamini, Y., and Y. Hochberg. 1995. Controlling the False Discovery Rate: A Practical and Powerful Approach to Multiple Testing. Journal of the Royal Statistical Society Series B: Statistical Methodology 57:289–300.

Biggs, C.R., S.K. Lowerre-Barbieri, and B. Erisman. 2018. Reproductive resilience of an estuarine fish in the eye of a hurricane. Biology Letters 14.11: 20180579.

Blowes, S. A., B. McGill, V. Brambilla, C. F. Y. Chow, T. Engel, A. Fontrodona-Eslava, I. S. Martins, D. McGlinn, F. Moyes, A. Sagouis, H. Shimadzu, R. van Klink, W. B. Xu, N. J. Gotelli, A. Magurran, M. Dornelas, and J. M. Chase. 2024. Synthesis reveals approximately balanced biotic differentiation and homogenization. Sci Adv 10:eadj9395.

Blowes, S. A., S. R. Supp, L. H. Antao, A. Bates, H. Bruelheide, J. M. Chase, F. Moyes, A. Magurran, B. McGill, I. H. Myers-Smith, M. Winter, A. D. Bjorkman, D. E. Bowler, J. E. K. Byrnes, A. Gonzalez, J. Hines, F. Isbell, H. P. Jones, L. M. Navarro, P. L. Thompson, M. Vellend, C. Waldock, and M. Dornelas. 2019. The geography of biodiversity change in marine and terrestrial assemblages. Science 366:339–345.

Boettiger, C., D. T. Lang, and P. C. Wainwright. 2012. rfishbase: exploring, manipulating and visualizing FishBase data from R. Journal of Fish Biology 81:2030–2039.

Bolker, B. M., M. E. Brooks, C. J. Clark, S. W. Geange, J. R. Poulsen, M. H. Stevens, and J. S. White. 2009. Generalized linear mixed models: a practical guide for ecology and evolution. Trends Ecol Evol 24:127–135, <10.1016/j.tree.2008.10.008>.

Bugica, K., B. Sterba-Boatwright, and M. S. Wetz. 2020. Water quality trends in Texas estuaries. Marine Pollution Bulletin 152:110903.

Chamberlain, S., and B. Vanhoorne. 2024. worrms: World Register of Marine Species (WoRMS) Client. R package version 0.4.3. https://CRAN.R-project.org/package=worrms

Chao, A., C.-H. Chiu, and L. Jost. 2014. Unifying species diversity, phylogenetic diversity, functional diversity, and related similarity and differentiation measures through Hill numbers. Annual Review of Ecology, Evolution, and Systematics 45:297–324.

Chao, A., and L. Jost. 2012. Coverage-based rarefaction and extrapolation: standardizing samples by completeness rather than size. Ecology 93:2533–2547.

Day, J. W., B. C. Crump, W. M. Kemp, and A. Yáñez-Arancibia. 2012. Estuarine Ecology, 2nd Ed. Wiley-Blackwell, Hoboken, New Jersey.

Dornelas, M., N. J. Gotelli, B. McGill, H. Shimadzu, F. Moyes, C. Sievers, and A. E. Magurran. 2014. Assemblage time series reveal biodiversity change but not systematic loss. Science 344:296–299.

Du, J., and K. Park. 2019. Estuarine salinity recovery from an extreme precipitation event: Hurricane Harvey in Galveston Bay. Science of the Total Environment 670: 1049–1059.

Elliott, M., and A. K. Whitfield. 2011. Challenging paradigms in estuarine ecology and management. Estuarine, Coastal and Shelf Science 94:306–314.

Feagin, R.A., J.E. Lerner, C. Noyola, T.P. Huff, J. Madewell, and B. Balboa. 2024. Hypersalinity in coastal wetlands and potential restoration solutions, Lake Austin and East Matagorda Bay, Texas, USA. Journal of Marine Science and Engineering 12.5: 829.

Froese, R., and D. Pauly. 2019. FishBase, World Wide Web electronic publication. www.fishbase.org, version (12/2019).

Fujiwara, M., F. Martinez-Andrade, R. J. D. Wells, M. Fisher, M. Pawluk, and M. C. Livernois. 2019. Climate-related factors cause changes in the diversity of fish and invertebrates in subtropical coast of the Gulf of Mexico. Communications Biology 2:403.

Fujiwara, M., A. Simpson, M. Torres-Ceron, and F. Martinez-Andrade. 2022. Life-history traits and temporal patterns in the incidence of coastal fishes experiencing tropicalization. Ecosphere 13.

GBIF. 2025. GBIF Home Page, World Wide Web electronic publication. https://www.gbif.org/.

Gotelli, N. J., and R. K. Colwell. 2001. Quantifying biodiversity: procedures and pitfalls in the measurement and comparison of species richness. Ecology Letters 4:379–391.

Hsieh, T. C., K. H. Ma, A. Chao, and G. McInerny. 2016. iNEXT: an R package for rarefaction and extrapolation of species diversity (Hill numbers). Methods in Ecology and Evolution 7:1451–1456.

Hubert, W. A., K. L. Pope, and J. M. Dettmers. 2012. Passive capture techniques. Pages 223–265 in A. V. Zale, D. L. Parrish, and T. M. Sutton, editors. Fisheries Techniques, 3rd Ed. American Fisheries Society, Bethesda, MD.

IPCC. 2021. Climate Change 2021: The Physical Science Basis. Contribution of Working Group I to the Sixth Assessment Report of the Intergovernmental Panel on Climate Change. Cambridge, United Kingdom; New York, NY, USA.

Jentsch, A., J. Kreyling, and C. Beierkuhnlein. 2007. A new generation of climate-change experiments: events, not trends. Frontiers in Ecology and the Environment 5:365–374.

Jost, L. 2006. Entropy and diversity. Oikos 113:363–375.

Kuznetsova, A., P. B. Brockhoff, and R. H. B. Christensen. 2017. lmerTest Package: Tests in Linear Mixed Effects Models. Journal of Statistical Software 82.

Lavorel, S., and E. Garnier. 2002. Predicting changes in community composition and ecosystem functioning from plant traits: revisiting the Holy Grail. Functional Ecology 16:545–556.

Leibold, M. A., M. Holyoak, N. Mouquet, P. Amarasekare, J. M. Chase, M. F. Hoopes, R. D. Holt, J. B. Shurin, R. Law, D. Tilman, M. Loreau, and A. Gonzalez. 2004. The metacommunity concept: a framework for multi-scale community ecology. Ecology Letters 7:601–613.

Lenth, R. V. 2024. emmeans: Estimated Marginal Means, aka Least-Squares Means. R package.

Levin, S. A. 1992. The problem of pattern and scale in ecology: The Robert H. MacArthur award lecture. Ecology 73:1943–1967.

Longley, W. L., editor. 1994. Freshwater inflows to Texas bays and estuaries: ecological relationships and methods for determination of needs. Texas Water Development Board; Texas Parks and Wildlife Department, Austin, TX.

Loreau, M., S. Naeem, P. Inchausti, J. Bengtsson, J. P. Grime, A. Hector, D. U. Hooper, M. A. Huston, D. Raffaelli, B. Schmid, D. Tilman, and D. A. Wardle. 2001. Biodiversity and ecosystem functioning: current knowledge and future challenges. Science 294:804–808.

Magurran, A. E., S. R. Baillie, S. T. Buckland, J. M. Dick, D. A. Elston, E. M. Scott, R. I. Smith, P. J. Somerfield, and A. D. Watt. 2010. Long-term datasets in biodiversity research and monitoring: assessing change in ecological communities through time. Trends Ecol Evol 25:574–582.

Magurran, A. E., M. Dornelas, F. Moyes, P. A. Henderson, and D. Storch. 2019. Temporal β diversity—A macroecological perspective. Global Ecology and Biogeography 28:1949–1960.

Magurran, A. E., and P. A. Henderson. 2010. Temporal turnover and the maintenance of diversity in ecological assemblages. Philos Trans R Soc Lond B Biol Sci 365:3611–3620.

Martinez-Andrade, F. 2018. Marine Resource Monitoring Operations Manual. Page 137. Texas Parks & Wildlife Department, Coastal Fisheries Division, Austin, Texas.

McCann, K. S. 2000. The diversity-stability debate. Nature 405:228–233.

McEachron, L. W., G. C. Matlock, C. E. Bryan, P. Unger, T. J. Cody, and J. H. Martin. 1994. Winter mass mortality of animals in Texas bays. Northeast Gulf Science 13 (2). DOI: 10.18785/negs.1302.06

McGill, B. J., M. Dornelas, N. J. Gotelli, and A. E. Magurran. 2015. Fifteen forms of biodiversity trend in the Anthropocene. Trends in Ecology & Evolution 30:104–113.

McGill, B. J., B. J. Enquist, E. Weiher, and M. Westoby. 2006. Rebuilding community ecology from functional traits. Trends in Ecology & Evolution 21:178–185.

McGill, B. J., R. S. Etienne, J. S. Gray, D. Alonso, M. J. Anderson, H. K. Benecha, M. Dornelas, B. J. Enquist, J. L. Green, F. He, A. H. Hurlbert, A. E. Magurran, P. A. Marquet, B. A. Maurer, A. Ostling, C. U. Soykan, K. I. Ugland, and E. P. White. 2007. Species abundance distributions: moving beyond single prediction theories to integration within an ecological framework. Ecology Letters 10:995–1015.

NOAA National Hurricane Center. 2025. Tropical Cyclone Reports. National Oceanic and Atmospheric Administration, Miami, Florida, USA.

OBIS. 2025. Ocean Biodiversity Information System. World Wide Web electronic publication. https://obis.org. Intergovernmental Oceanographic Commission of UNESCO.

Olden, J. D., and T. P. Rooney. 2006. On defining and quantifying biotic homogenization. Global Ecology and Biogeography 15:113–120.

Palomares, M. L. D., and D. Pauly. 2025. SeaLifeBase. World Wide Web electronic publication. www.sealifebase.org.

Patrick, C.J., L. Yeager, A.R. Armitage, F. Carvallo, V.M. Congdon, K.H. Dunton, M. Fisher, A.K. Hardison, J.D. Hogan, J. Hosen, and X. Hu. 2020. A system level analysis of coastal ecosystem responses to hurricane impacts. Estuaries and Coasts 43.5: 943–959.

Parmesan, C., and G. Yohe. 2003. A globally coherent fingerprint of climate change impacts across natural systems. Nature 421:37–42.

Pawluk, M., M. Fujiwara, and F. Martinez-Andrade. 2021. Climate effects on fish diversity in the subtropical bays of Texas. Estuarine, Coastal and Shelf Science 249:107121.

Pinsky, M. L., B. Worm, M. J. Fogarty, J. L. Sarmiento, and S. A. Levin. 2013. Marine taxa track local climate velocities. Science 341:1239–1242.

Poirrier, M.A., Z.R. Del Rey, and E.A. Spalding. 2008. Acute disturbance of Lake Pontchartrain benthic communities by Hurricane Katrina. Estuaries and Coasts 31.6: 1221–1228.

R Core Team. 2024. R: A language and environment for statistical computing. Version 4.5.0. R Foundation for Statistical Computing, Vienna, Austria. https://www.R-project.org/.

Scheffer, M., S. Carpenter, J. A. Foley, C. Folke, and B. Walker. 2001. Catastrophic shifts in ecosystems. Nature 413:591–596.

Sheaves, M., R. Baker, I. Nagelkerken, and R. M. Connolly. 2015. True Value of Estuarine and Coastal Nurseries for Fish: Incorporating Complexity and Dynamics. Estuaries and Coasts 38:401–414.

Simpson, G. L. 2023. gratia: Graceful ggplot-based graphics and utilities for GAMs fitted using mgcv, Version 0.9.0. R Foundation for Statistical Computing, Vienna, Austria.

Smith, M. D. 2011. An ecological perspective on extreme climatic events: a synthetic definition and framework to guide future research. Journal of Ecology 99:656–663.

Sunday, J. M., A. E. Bates, and N. K. Dulvy. 2012. Thermal tolerance and the global redistribution of animals. Nature Climate Change 2:686–690.

Texas Parks and Wildlife Department. 2021. At least 3.8 million fish killed by winter weather on the Texas coast. Texas Parks and Wildlife Department News Release, 10 March 2021. Texas Parks and Wildlife Department, Austin, Texas, USA. https://tpwd.texas.gov/newsmedia/releases/?req=20210310c

Thibaut, L. M., and S. R. Connolly. 2013. Understanding diversity-stability relationships: towards a unified model of portfolio effects. Ecology Letters 16:140–150.

Tolan, J. M., and M. Fisher. 2009. Biological response to changes in climate patterns: population increases of Gray Snapper (Lutjanus griseus) in Texas bays and estuaries. Fishery Bulletin 107:36–44.

Torres Ceron, M., M. Fujiwara, and F. Martinez-Andrade. 2023. Changes in species compositions of fish in the bays of the Northwestern Gulf of Mexico. Frontiers in Marine Science 10.

Turner, M. G. 2010. Disturbance and landscape dynamics in a changing world. Ecology 91:2833–2849.

Vergés, A., E. McCosker, M. Mayer-Pinto, M. A. Coleman, T. Wernberg, T. Ainsworth, P. D. Steinberg, and G. Williams. 2019. Tropicalisation of temperate reefs: Implications for ecosystem functions and management actions. Functional Ecology 33:1000–1013.

Vincenzi, S., A. J. Crivelli, W. H. Satterthwaite, and M. Mangel. 2014. Eco-evolutionary dynamics induced by massive mortality events. Journal of Fish Biology 85.1: 8–30.

Wang, Z., T. Boyer, J. Reagan, and P. Hogan. 2023. Upper-Oceanic Warming in the Gulf of Mexico between 1950 and 2020. Journal of Climate 36:2721–2734.

Whitfield, A. K., M. Elliott, A. Basset, S. J. M. Blaber, and R. J. West. 2012. Paradigms in estuarine ecology – A review of the Remane diagram with a suggested revised model for estuaries. Estuarine, Coastal and Shelf Science 97:78–90.

Whittaker, R. H. 1960. Vegetation of the Siskiyou Mountains, Oregon and California. Ecological Monographs 30:279–338.

Whittaker, R. H. 1972. Evolution and Measurement of Species Diversity. Taxon 21:213–251.

Williford, D., and J. Anderson. 2025. Assessing Long-Term Changes to Estuarine Benthic Communities in the Northwestern Gulf of Mexico. Estuaries and Coasts 48.

Winemiller, K. O., and K. A. Rose. 1992. Patterns of Life-History Diversification in North American Fishes: implications for Population Regulation. Canadian Journal of Fisheries and Aquatic Sciences 49:2196–2218.

Withers, K., B. R. Chapman, J. W. Tunnell, Jr., and F. W. Judd, editors. 2017. The Laguna Madre of Texas and Tamaulipas, Revised Edition. Texas A&M University Press, College Station, Texas.

Wood, S. N. 2017. Generalized additive models: An introduction with R, 2nd Ed. Chapman and Hall/CRC, Boca Raton, FL.

Zeileis, A., and G. Grothendieck. 2005. zoo: S3 infrastructure for regular and irregular time series. Journal of Statistical Software 14:1–27.

