## Supplement Figures for "Community reorganization without collapse in a warming world: Habitat-contingent, trait-mediated biodiversity change within a conserved multi-scale structure"

### SUPPLEMENTARY FIGURES

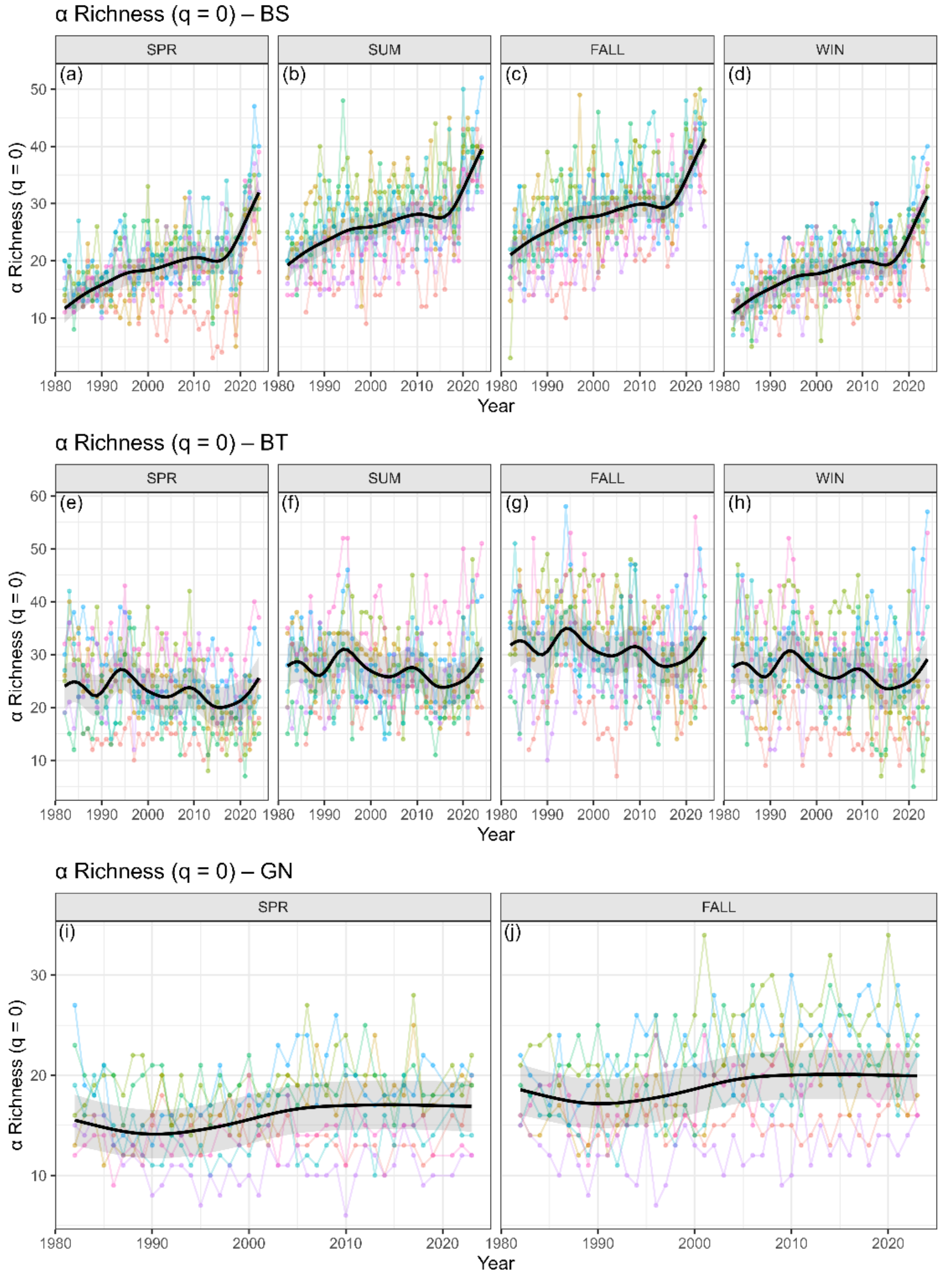


**Figure** S1. α-diversity trends for richness ($q_0$). Coverage-standardized α-richness ($q_0$) for each gear × season × year combination. Points represent annual estimates for individual bays, and black curves show fitted temporal trends with 95% confidence intervals.

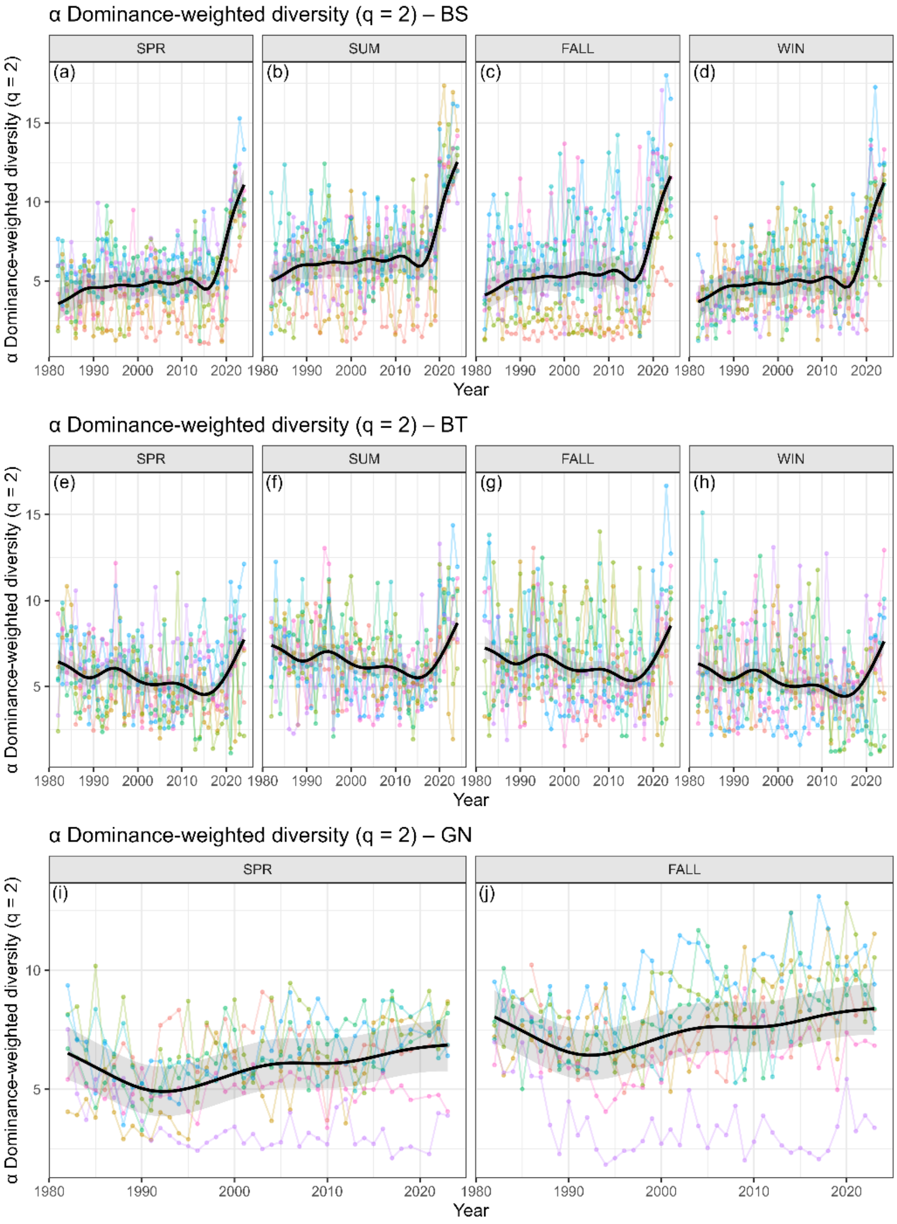


**Figure S2.** α-diversity trends for Hill $q_2$. Coverage-standardized α-diversity for Hill number $q_2$ for each gear × season × year combination. Points represent annual estimates for individual bays, and black curves show fitted temporal trends with 95% confidence intervals.

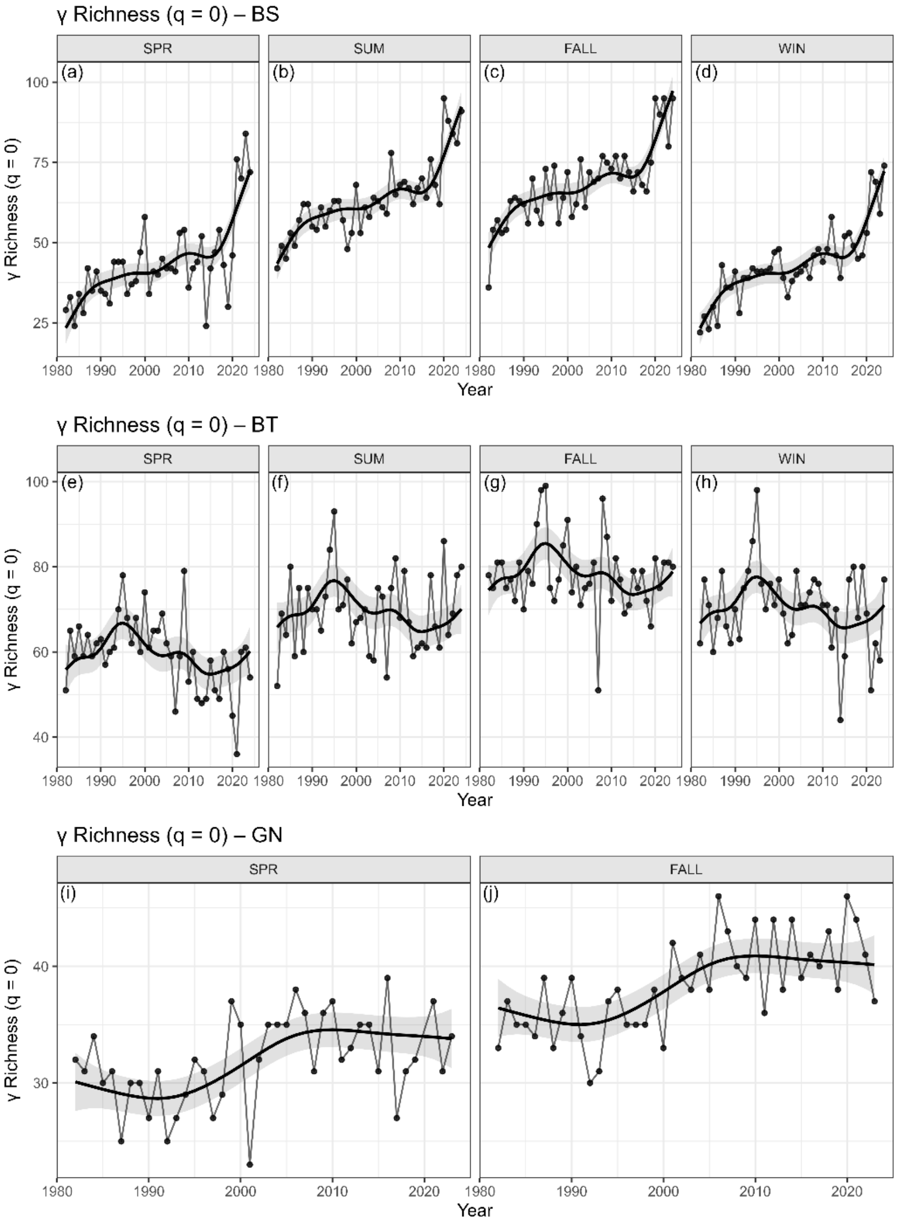


**Figure S3.** γ-diversity trends for richness ($q_0$). Coverage-standardized γ-richness ($q_0$), pooled across bays, shown for each gear × season × year combination. Points represent annual estimates, and fitted temporal trends with 95% confidence intervals are shown for each gear.

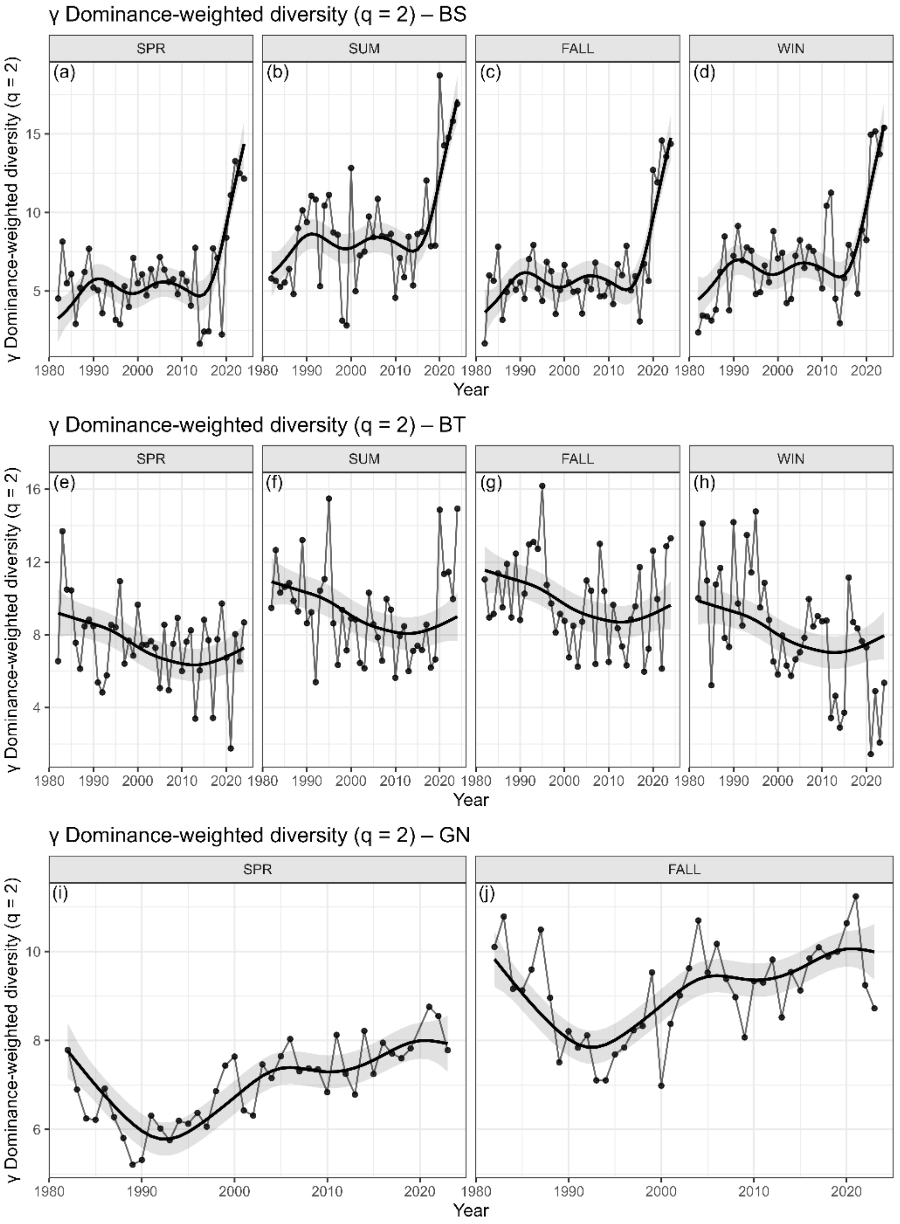


**Figure S4.** γ-diversity trends for Hill $q_2$. Coverage-standardized γ-diversity for Hill number $q_2$, pooled across bays, shown for each gear × season × year combination. Points represent annual estimates, and fitted temporal trends with 95% confidence intervals are shown for each gear.

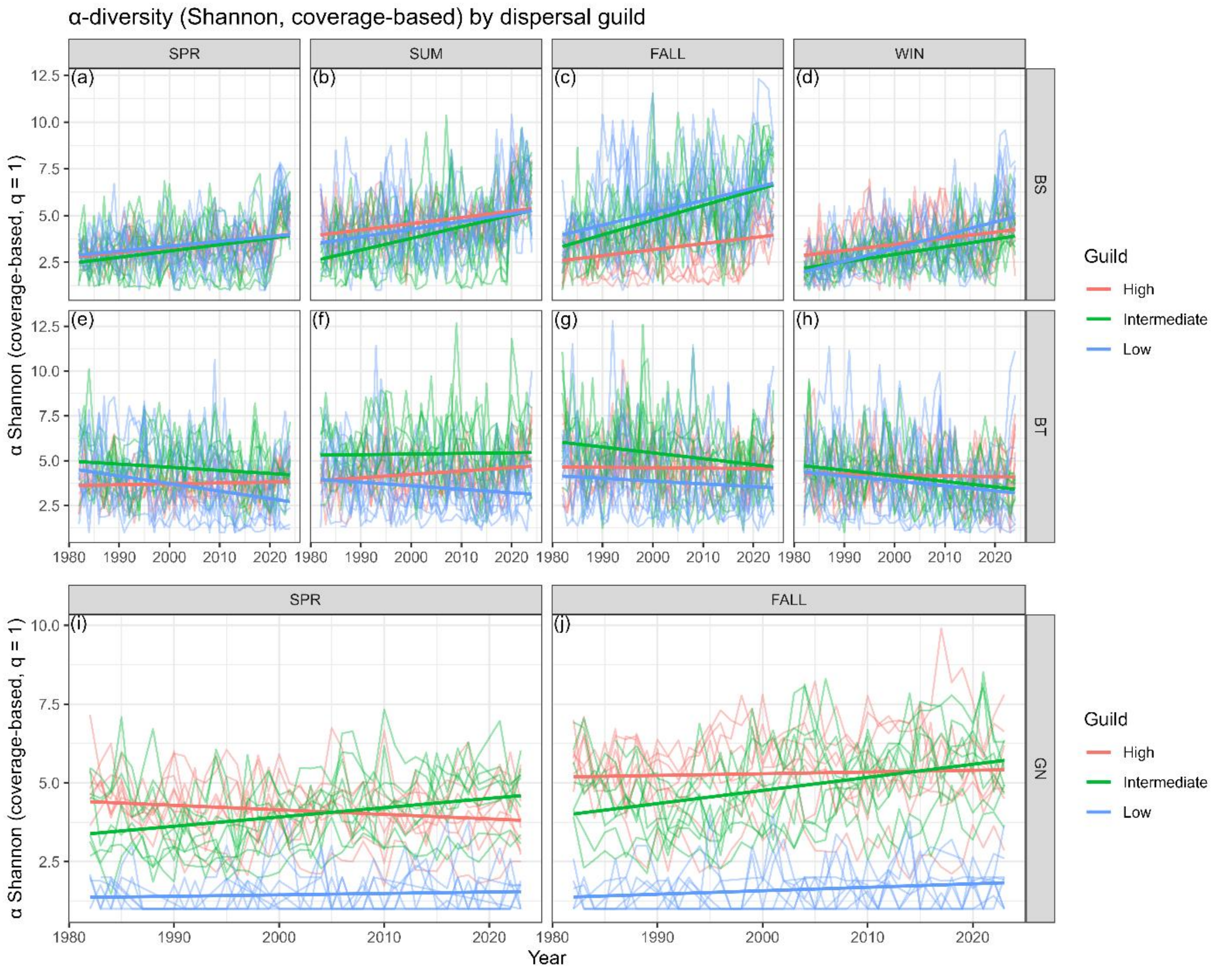


**Figure S5.** Long-term trends in coverage-based Shannon α-diversity (q = 1) for dispersal guilds across gears and seasons. Panels show annual α-diversity for Low-, Intermediate-, and High-dispersal taxa within each gear (bag seine, bay trawl, and gillnet) and season (WIN, SPR, SUM, FALL). Thin lines represent individual bay × guild time series, and thick lines show linear trends estimated from mixed-effects models. Panel letters (a–j) correspond to the order of gear–season combinations.

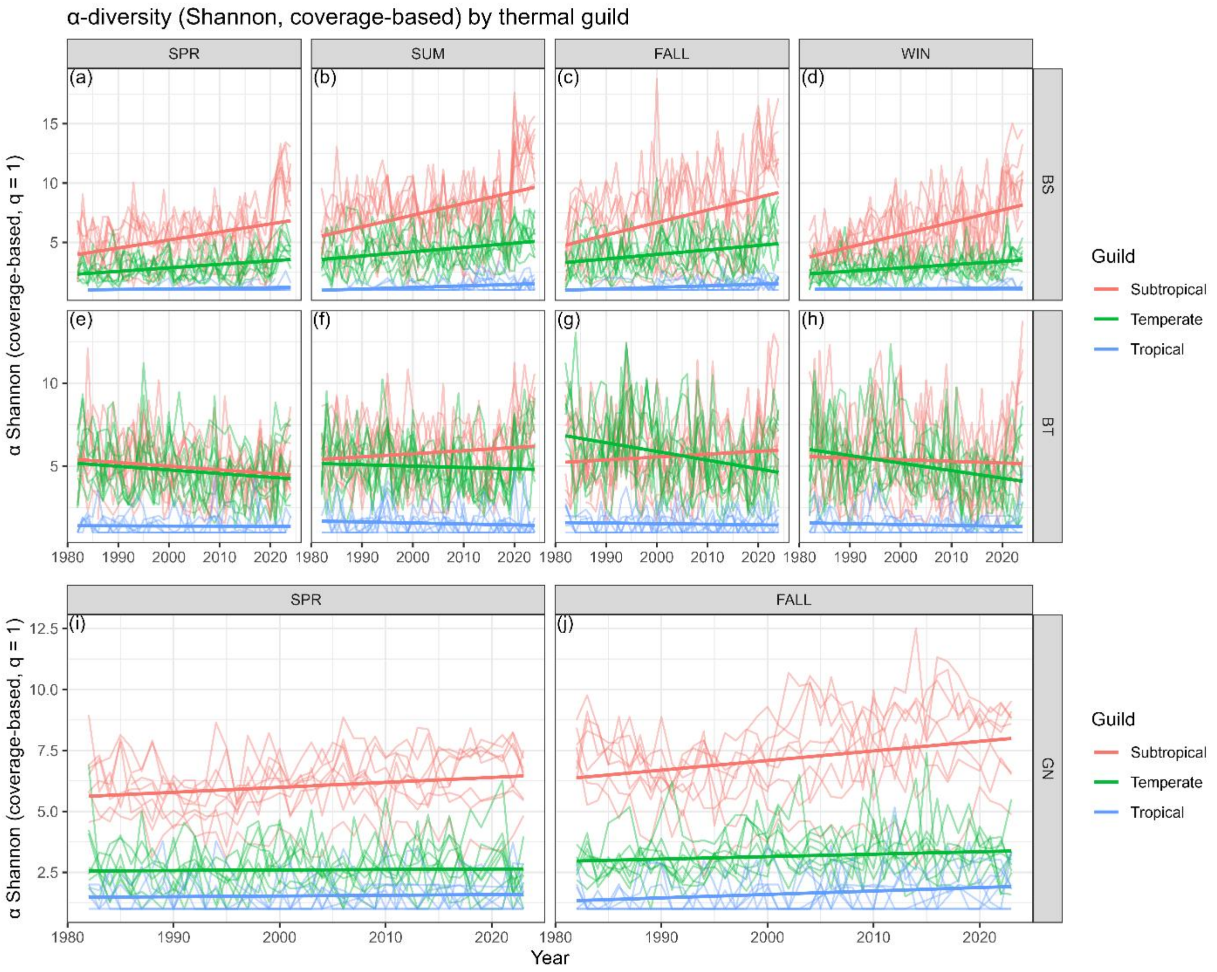


**Figure S6.** Long-term trends in coverage-based Shannon α-diversity (q = 1) for thermal-affinity guilds across gears and seasons. Panels show annual α-diversity for Tropical, Subtropical, and Temperate taxa within each gear (bag seine, bay trawl, and gillnet) and season (WIN, SPR, SUM, FALL). Thin lines represent individual bay × guild time series, and thick lines denote linear trends from mixed-effects models. Panel letters (a–j) correspond to the sequence of gear–season combinations.

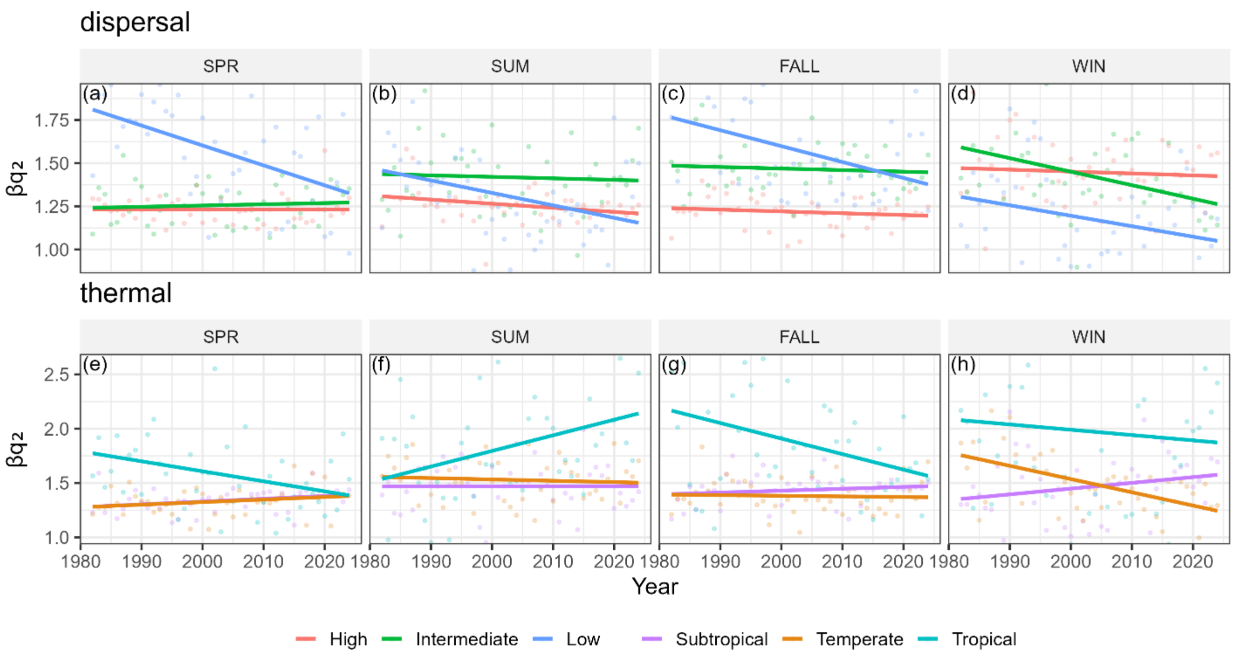


**Figure S7.** Spatial dominance-weighted β-diversity ($\beta q_2$) by functional guild, pooled across gears and bays. Panels show dispersal guilds (high, intermediate, low) and thermal-affinity guilds (subtropical, temperate, tropical), with seasons arranged in columns. Points represent annual mean $\beta q_2$ values averaged across major bays and sampling gears for each year × season × guild combination, and solid lines indicate fitted linear trends through time. Declining trends indicate increasing spatial homogenization when β-diversity is weighted toward common and dominant taxa.

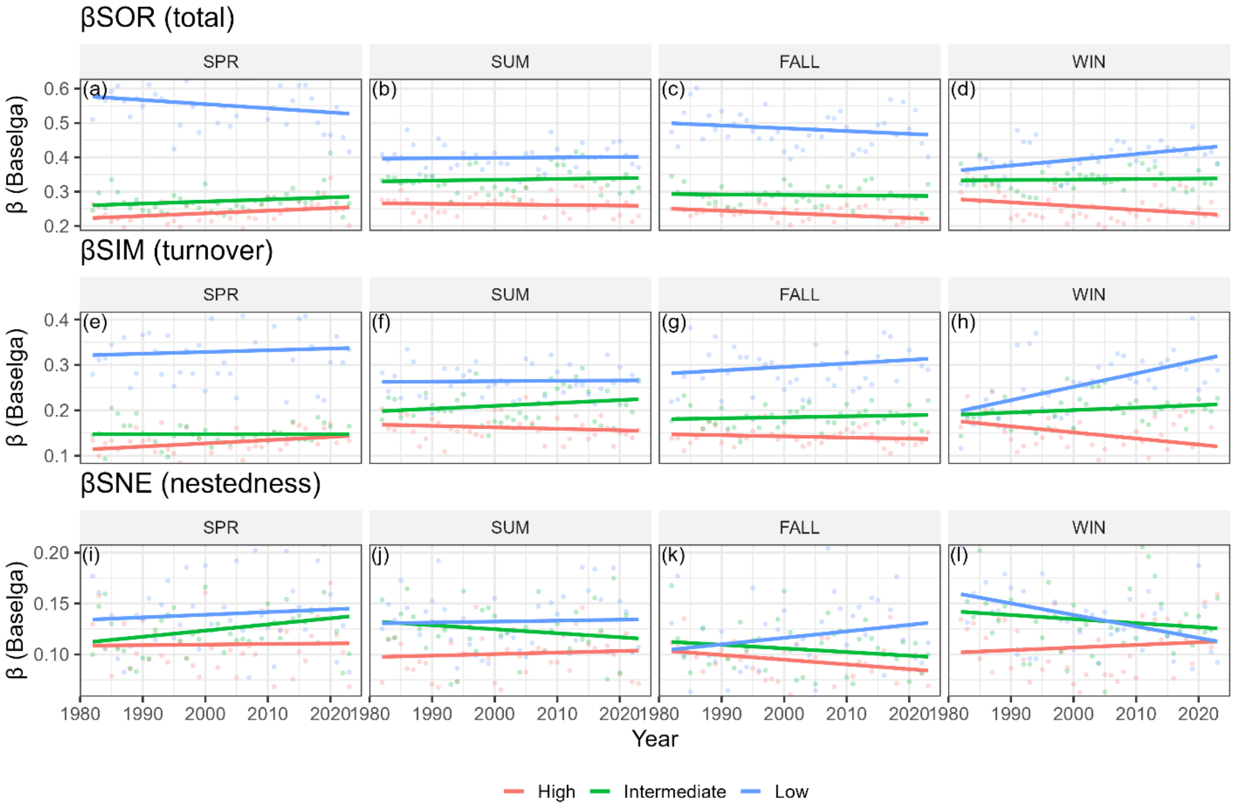


**Figure S8**. Temporal Baselga components for dispersal guilds. Data are pooled across major bays and sampling gears. Panels are arranged by season (columns) and diversity component (rows: total βSOR, turnover βSIM, nestedness βSNE). Colored lines denote fitted trends for High, Intermediate, and Low dispersal guilds.

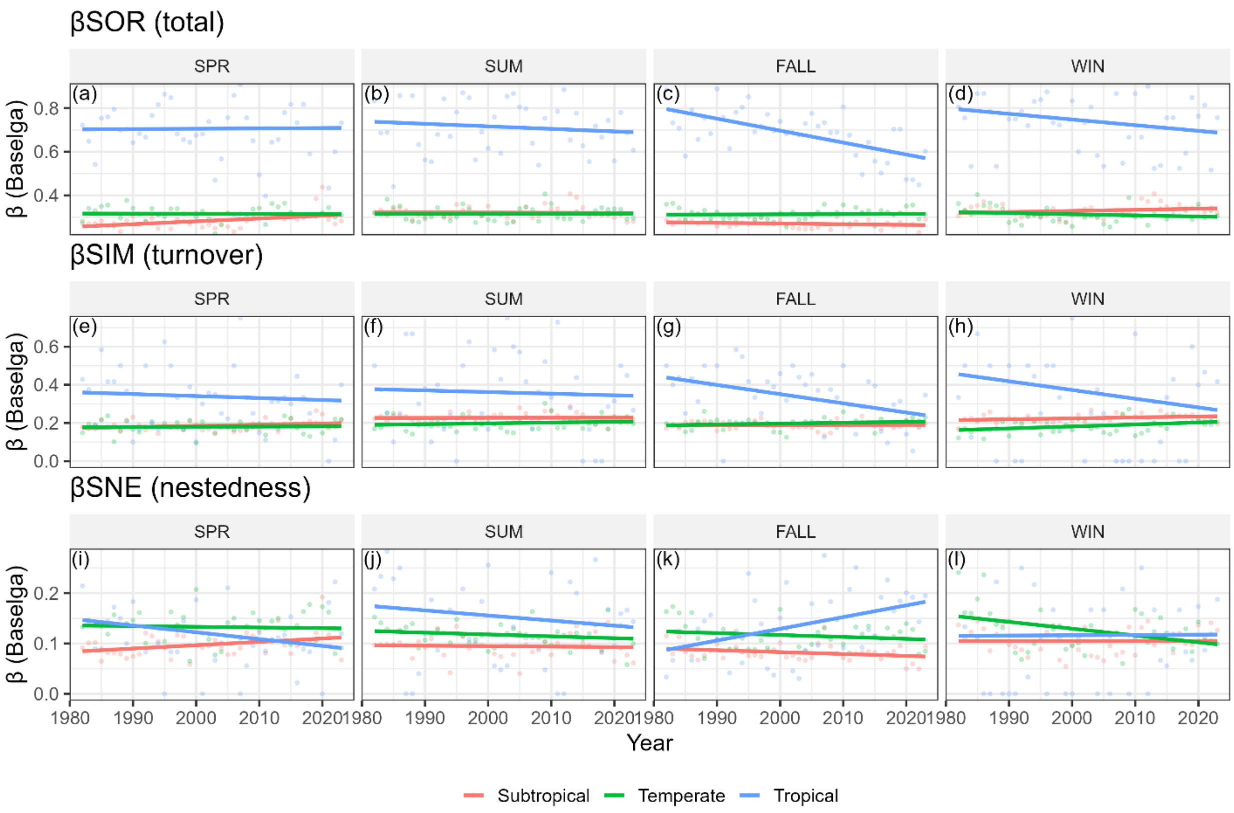


**Figure S9.** Temporal Baselga components for thermal guilds. Data are pooled across major bays and sampling gears. Panels are arranged by season (columns) and diversity component (rows: total βSOR, turnover βSIM, nestedness βSNE). Colored lines denote fitted trends for Subtropical, Temperate, and Tropical thermal guilds.

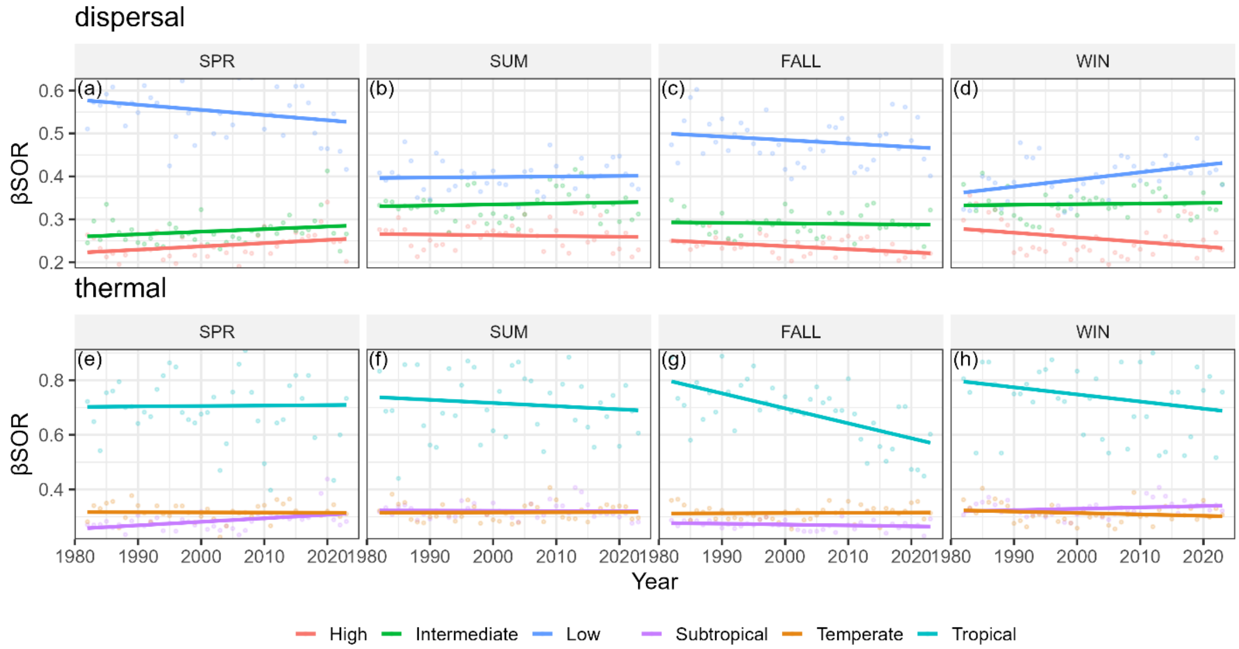


**Figure S10.** Temporal β-diversity (βSOR, Δyear = 1) by functional guild levels, pooled across gears and bays. Annual mean βSOR values are shown separately for dispersal guild levels (low, intermediate, high) and thermal-affinity guild levels (tropical, subtropical, temperate), with seasons displayed in columns. Points represent year-to-year compositional dissimilarity averaged across major bays and sampling gears for each consecutive year pair, and solid lines indicate fitted linear trends. The figure illustrates differences in long-term temporal homogenization dynamics among functional guilds.

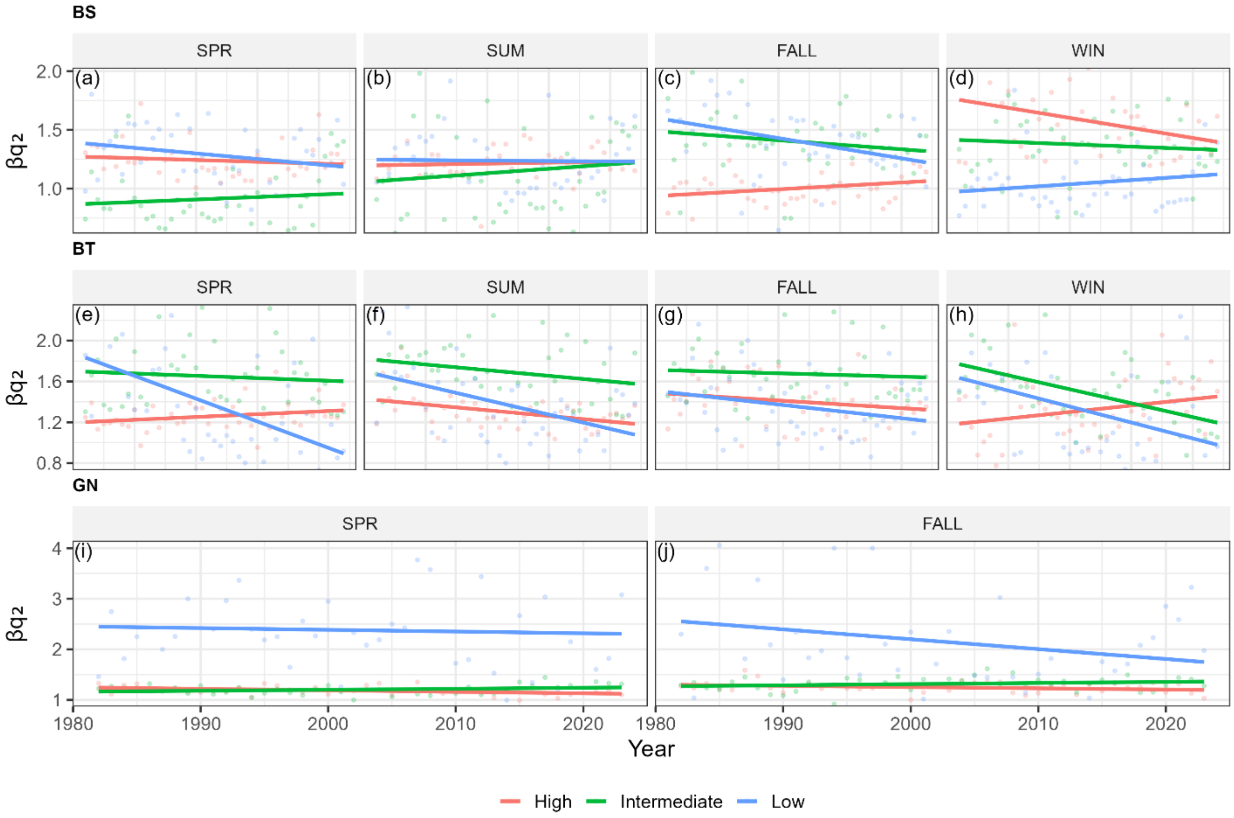


**Figure S11.** Gear-specific trends in spatial dominance-weighted β-diversity ($\beta q_2$) for dispersal guilds. Panels are arranged by gear (rows) and season (columns). Points show annual mean $\beta q_2$ values pooled across bays within each gear × season × guild combination, and solid lines indicate fitted linear trends. Patterns illustrate strong habitat dependence in the magnitude and direction of spatial homogenization among dispersal guilds.

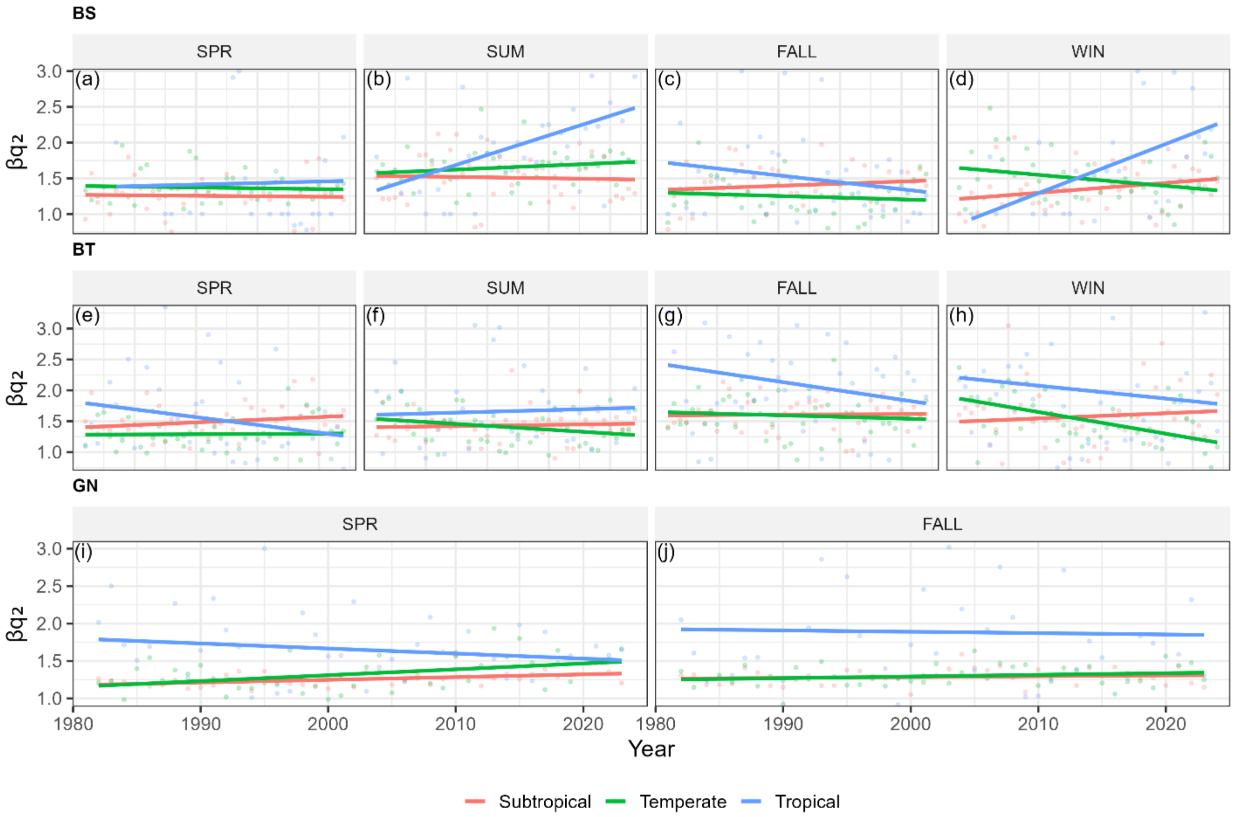


**Figure S12**. Gear-specific trends in spatial dominance-weighted β-diversity ($\beta q_2$) for thermal-affinity guilds. Panels are organized by gear (rows) and season (columns). Points represent annual mean $\beta q_2$ values pooled across bays within each gear × season × guild combination, and solid lines show linear trends through time. Results highlight contrasting habitat-specific trajectories of spatial convergence and divergence among thermal-affinity groups.

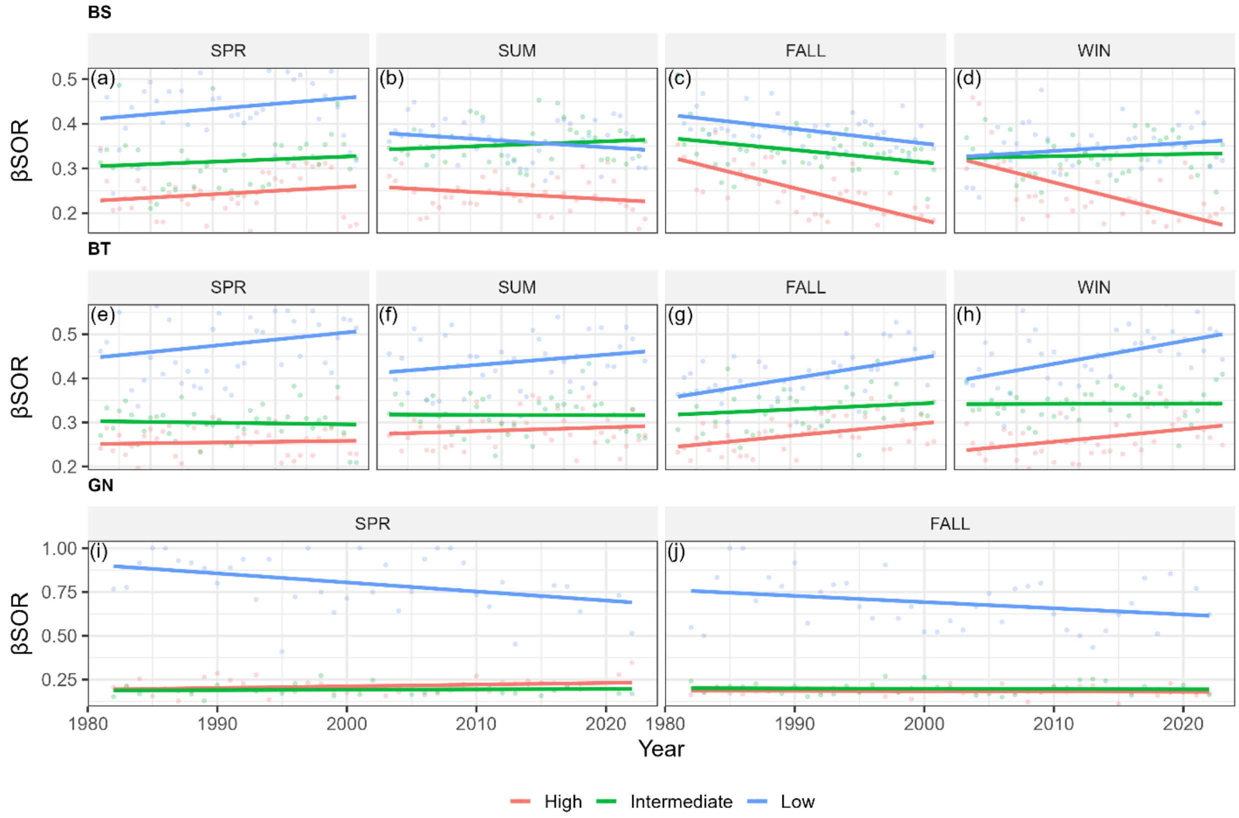


**Figure S13**. Gear-specific trends in temporal β-diversity (βSOR, Δyear = 1) for dispersal guilds. Panels are arranged by gear (rows) and season (columns). Points show annual mean βSOR values pooled across bays within each gear × season × guild combination, and solid lines indicate linear trends through time. Y-axis ranges are scaled independently by gear to emphasize within-habitat temporal dynamics and reveal limited directional change in most dispersal guilds.

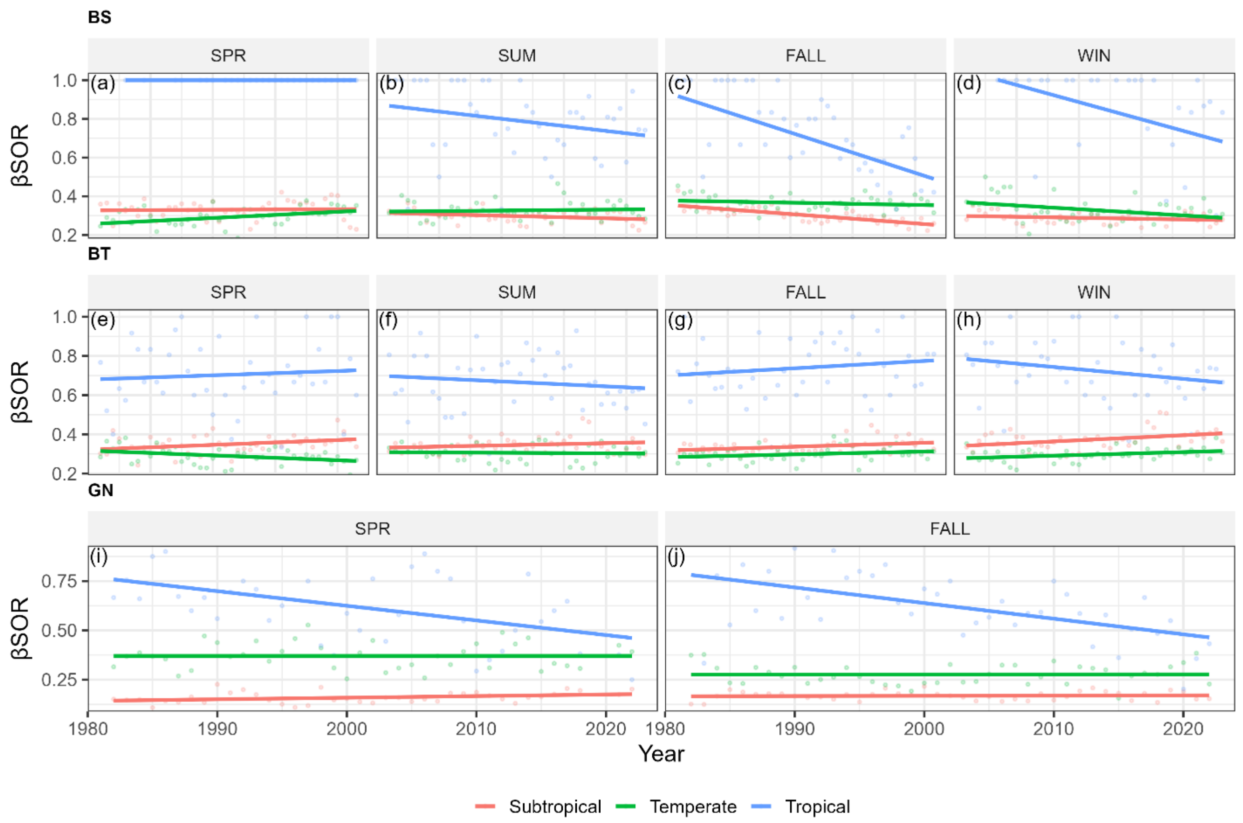


**Figure S14.** Gear-specific trends in temporal β-diversity (βSOR, Δyear = 1) for thermal-affinity guilds. Panels are arranged by gear (rows) and season (columns). Points represent annual mean βSOR values pooled across bays within each gear × season × guild combination, and solid lines denote fitted linear trends. The figure highlights pronounced temporal homogenization in tropical taxa relative to subtropical and temperate groups, with habitat-dependent variation in trend strength.

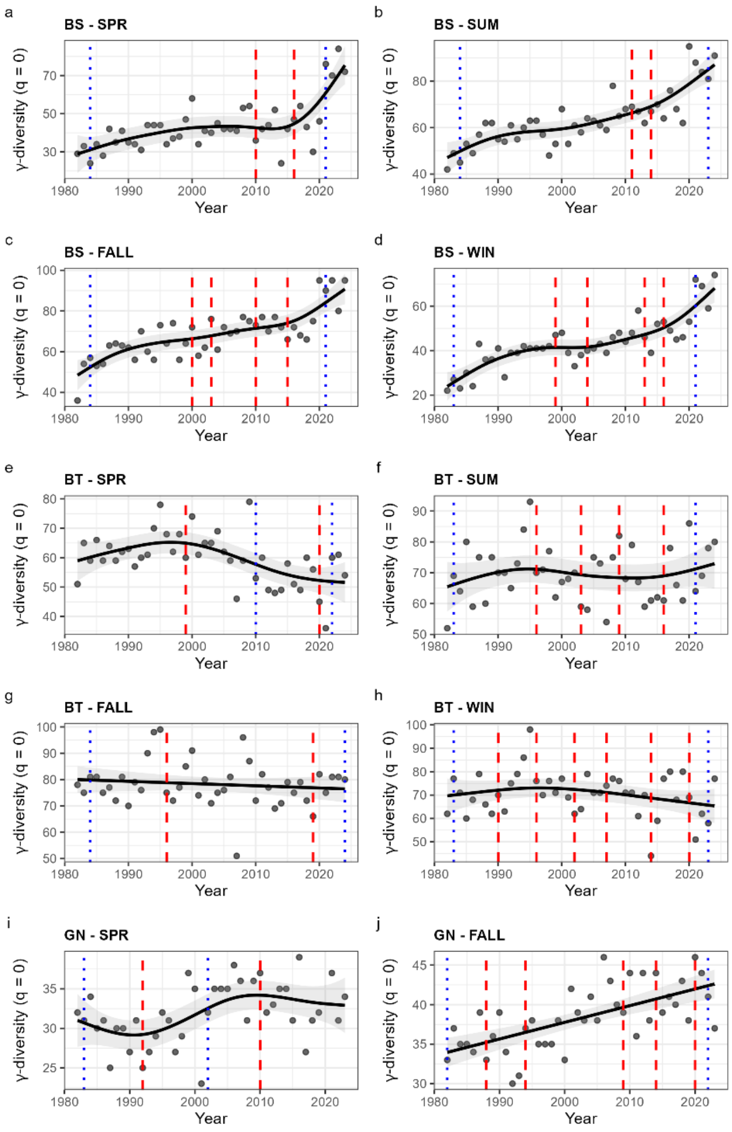


**Figure S15.** Long-term trajectories of regional (γ) diversity based on coverage-standardized richness (q = 0) across all gear and season combinations. Points show annual estimates averaged across bays, and solid lines show fitted generalized additive model (GAM) smooths with shaded ribbons indicating 95% confidence intervals. Panels are ordered by gear and season and labeled sequentially (a–j). Vertical red dashed lines mark years in which the first derivative of the fitted smooth exhibited a confidence-supported sign change, indicating a statistically supported reversal in long-term trend direction. Vertical black dotted lines mark peaks in the rate of change (extrema in the absolute second derivative), indicating periods of maximum acceleration or deceleration. GAMs were fit separately for each gear and season using shrinkage smooths, and diagnostic years correspond to Table S8.
