## Supplement Tables for "Community reorganization without collapse in a warming world: Habitat-contingent, trait-mediated biodiversity change within a conserved multi-scale structure"

### SUPPLEMENTARY TABLES

**Table S1.** Long-term temporal trends in α- and γ-diversity across sampling gears. Slopes represent annual and decadal rates of change in coverage-standardized Hill diversity (q = 0, 1, 2). α-diversity trends were estimated using linear mixed-effects models with year and season as fixed effects and bay as a random intercept, while γ-diversity trends were estimated using linear models with year and season as fixed effects when multiple seasons were available (otherwise year only). False-discovery-rate (FDR) correction was applied within each diversity order across gears. Positive slopes indicate increasing diversity through time, whereas negative slopes indicate declining diversity.

| **Metric** | **Gear** | **Slope per year** | **Slope per decade** | **SE** | **t** | **FDR-p** |
|---|---|---|---|---|---|---|
| α Richness (q=0) | BS | **0.316** | **3.163** | **0.012** | **26.702** | **$4.925 \times 10^{-126}$** |
| α Richness (q=0) | BT | ***-0.079*** | ***-0.786*** | **0.014** | **-5.437** | **$6.436 \times 10^{-8}$** |
| α Richness (q=0) | GN | **0.083** | **0.833** | **0.009** | **8.871** | **$1.048 \times 10^{-17}$** |
| α Shannon (q=1) | BS | **0.143** | **1.433** | **0.007** | **20.033** | **$6.663 \times 10^{-78}$** |
| α Shannon (q=1) | BT | ***-0.026*** | ***-0.258*** | **0.010** | **-2.696** | **$7.244 \times 10^{-3}$** |
| α Shannon (q=1) | GN | **0.043** | **0.425** | **0.006** | **7.372** | **$1.582 \times 10^{-13}$** |
| α Dominance (q=2) | BS | **0.094** | **0.942** | **0.006** | **15.177** | **$4.602 \times 10^{-33}$** |
| α Dominance (q=2) | BT | *-0.009* | *-0.088* | 0.008 | -1.035 | $3.009 \times 10^{-1}$ |
| α Dominance (q=2) | GN | **0.034** | **0.343** | **0.005** | **7.358** | **$1.745 \times 10^{-13}$** |
| γ Richness (q=0) | BS | **0.728** | **7.280** | **0.049** | **14.953** | **$3.762 \times 10^{-32}$** |
| γ Richness (q=0) | BT | *-0.116* | *-1.164* | 0.054 | -2.153 | $3.275 \times 10^{-2}$ |
| γ Richness (q=0) | GN | **0.170** | **1.705** | **0.030** | **5.660** | **$3.413 \times 10^{-7}$** |
| γ Shannon (q=1) | BS | **0.192** | **1.922** | **0.023** | **8.517** | **$2.745 \times 10^{-14}$** |
| γ Shannon (q=1) | BT | ***-0.116*** | ***-1.163*** | **0.021** | **-5.578** | **$9.643 \times 10^{-8}$** |
| γ Shannon (q=1) | GN | **0.055** | **0.547** | **0.009** | **6.074** | **$5.983 \times 10^{-8}$** |
| γ Dominance (q=2) | BS | **0.124** | **1.236** | **0.017** | **7.456** | **$1.385 \times 10^{-11}$** |
| γ Dominance (q=2) | BT | ***-0.067*** | ***-0.672*** | **0.015** | **-4.461** | **$1.497 \times 10^{-5}$** |
| γ Dominance (q=2) | GN | **0.039** | **0.389** | **0.008** | **4.979** | **$5.404 \times 10^{-6}$** |

**Table S2.** Dispersal guild–specific long-term trends in α- and γ-diversity (Shannon, q = 1) Based on three-way LMMs (YEAR × gear × dispersal group + Season + (1|MAJOR_AREA)). Slopes represent yearly change in coverage-standardized Shannon α- or γ-diversity for each combination of gear (BS = bag seine; BT = bay trawl; GN = gillnet) and dispersal guild (High, Intermediate, Low). Significant slopes after FDR correction are shown in bold.

| **α-diversity** | | | | | |
|---|---|---|---|---|---|
| **Dispersal guild** | **Gear** | **Slope / yr** | **SE** | **t** | **p_FDR** |
| High | BS | 0.0328 | 0.0032 | 10.16 | **1.40E-23** |
| High | BT | 0.0051 | 0.0032 | 1.59 | 1.69E-01 |
| High | GN | -0.0029 | 0.0048 | -0.60 | 5.51E-01 |
| Intermediate | BS | 0.0549 | 0.0032 | 17.02 | **5.27E-64** |
| Intermediate | BT | -0.0183 | 0.0032 | -5.66 | **2.97E-08** |
| Intermediate | GN | 0.0368 | 0.0048 | 7.68 | **3.45E-14** |
| Low | BS | 0.0520 | 0.0032 | 16.06 | **3.03E-57** |
| Low | BT | -0.0251 | 0.0032 | -7.78 | **1.81E-14** |
| Low | GN | 0.0061 | 0.0057 | 1.06 | 3.44E-01 |
| **γ-diversity** | | | | | |
| **Dispersal guild** | **Gear** | **Slope / yr** | **SE** | **t** | **p_FDR** |
| High | BS | 0.0379 | 0.0092 | 4.11 | **8.72E-05** |
| High | BT | 0.0104 | 0.0092 | 1.12 | 3.13E-01 |
| High | GN | -0.0124 | 0.0137 | -0.91 | 4.11E-01 |
| Intermediate | BS | 0.0727 | 0.0092 | 7.89 | **1.35E-14** |
| Intermediate | BT | -0.0516 | 0.0092 | -5.60 | **6.46E-08** |
| Intermediate | GN | 0.0439 | 0.0137 | 3.21 | **2.65E-03** |
| Low | BS | 0.0711 | 0.0092 | 7.72 | **4.20E-14** |
| Low | BT | -0.0987 | 0.0092 | -10.71 | **8.65E-26** |
| Low | GN | 0.0098 | 0.0137 | 0.72 | 5.02E-01 |

**Table S3.** Thermal guild–specific long-term trends in α- and γ-diversity (Shannon, q = 1) Based on three-way LMMs (YEAR × gear × thermal group + Season + (1|MAJOR_AREA)). Slopes represent yearly change in coverage-standardized Shannon α- or γ-diversity for each combination of gear (BS, BT, GN) and thermal guild (Subtropical, Temperate, Tropical). Significant slopes after FDR correction are shown in bold.

**α-diversity**

| Thermal guild | Gear | Slope / yr | SE | t | p_FDR |
|---|---|---|---|---|---|
| Subtropical | BS | 0.0952 | 0.0038 | 24.78 | **2.83E-134** |
| Subtropical | BT | 0.0024 | 0.0038 | 0.64 | 5.51E-01 |
| Subtropical | GN | 0.0318 | 0.0057 | 5.57 | **4.62E-08** |
| Temperate | BS | 0.0340 | 0.0038 | 8.85 | **3.07E-18** |
| Temperate | BT | -0.0307 | 0.0038 | -8.00 | **3.71E-15** |
| Temperate | GN | 0.0073 | 0.0057 | 1.28 | 2.58E-01 |
| Tropical | BS | 0.0178 | 0.0076 | 2.33 | **3.20E-02** |
| Tropical | BT | -0.0035 | 0.0050 | -0.70 | 5.46E-01 |
| Tropical | GN | 0.0099 | 0.0071 | 1.39 | 2.28E-01 |

**γ-diversity**

| Thermal guild | Gear | Slope / yr | SE | t | p_FDR |
|---|---|---|---|---|---|
| Subtropical | BS | 0.1304 | 0.0101 | 12.89 | **9.00E-37** |
| Subtropical | BT | 0.0196 | 0.0101 | 1.93 | 7.35E-02 |
| Subtropical | GN | 0.0331 | 0.0150 | 2.21 | **4.49E-02** |
| Temperate | BS | 0.0484 | 0.0101 | 4.78 | **4.45E-06** |
| Temperate | BT | -0.0814 | 0.0101 | -8.05 | **4.99E-15** |
| Temperate | GN | 0.0215 | 0.0150 | 1.43 | 1.95E-01 |
| Tropical | BS | 0.0382 | 0.0124 | 3.08 | **3.72E-03** |
| Tropical | BT | -0.0218 | 0.0101 | -2.15 | **4.70E-02** |
| Tropical | GN | 0.0061 | 0.0150 | 0.40 | 6.87E-01 |

**Table S4.** Long-term β-diversity trends (non-guild analyses). Summary of long-term temporal trends in spatial and temporal β-diversity estimated from coverage-standardized community data pooled across taxa. Spatial β-diversity was quantified using multiplicative Hill-number–based β (q = 0, 1, 2), corresponding to richness-, Shannon-, and dominance-weighted dissimilarity. Spatial compositional change was further decomposed into Baselga components—total dissimilarity (βSOR), turnover (βSIM), and nestedness (βSNE). Temporal β-diversity was estimated using Baselga βSOR calculated between consecutive years (Δyear = 1). Reported values are estimated slopes per year from linear (mixed-)effects models including sampling gear and season as fixed effects, with bay included as a random intercept where applicable. For spatial metrics, bootstrap-based 95% confidence intervals (500 resamples) are shown. Statistical inference focuses on estimated slopes rather than omnibus tests.

| **Response** | **Slope per year** | **Slope Se** | **t** | **P value** | **Slope per decade** | **Boot SE** |
|---|---|---|---|---|---|---|
| Spatial β (multiplicative) | | | | | | |
| Beta q0 | 0.0024 | 0.0012 | 1.95 | 0.052 | 0.0236 | 0.0012 |
| Beta q | -0.0026 | 0.0010 | -2.56 | 0.010 | -0.0259 | 0.0010 |
| Beta q2 | -0.0025 | 0.0011 | -2.35 | 0.019 | -0.0252 | 0.0011 |
| Spatial Baselga components | | | | | | |
| Total dissimilarity (βSOR) | 0.0003 | 0.0001 | 2.25 | 0.025 | 0.0031 | 0.0002 |
| Turnover (βSIM) | 0.0005 | 0.0002 | 2.80 | 0.005 | 0.0045 | 0.0002 |
| Nestedness (βSNE) | -0.0001 | 0.0001 | -0.97 | 0.330 | -0.0014 | 0.0002 |
| Temporal β (Δyear = 1) | | | | | | |
| Beta temporal baselga sor | 0.0001 | 0.0001 | 0.89 | 0.375 | 0.0010 | nan |

**Table S5.** Guild-structured β-diversity trends. Estimated long-term trends in β-diversity stratified by functional guilds, including dispersal groups (Low, Intermediate, High) and thermal-affinity groups (Tropical, Subtropical, Temperate). Spatial guild-level β-diversity was quantified using dominance-weighted multiplicative β (q = 2), while temporal guild-level β-diversity was estimated using Baselga βSOR calculated between consecutive years (Δyear = 1). Slopes represent estimated rates of change per year from linear (mixed-)effects models that account for sampling gear and seasonal structure, with bay included as a random intercept where applicable. Temporal βSOR models were fitted on the logit-transformed scale (with bounds to ensure numerical stability), although values shown in figures are on the raw scale. These results summarize guild-specific trends; formal tests of differences among guild slopes are presented separately.

| Response | term | Slope per year | Slope SE | t | P value | Slope per decade |
|---|---|---|---|---|---|---|
| **Spatial $\beta q_2$ by guild** | | | | | | |
| **Beta q2 dispersal** | YEAR | -0.0040 | 0.0005 | -7.61 | <0.001 | -0.0398 |
| **Beta q2 thermal** | YEAR | -0.0014 | 0.0006 | -2.18 | 0.029 | -0.0138 |
| **Temporal Baselga βSOR by guild (logit)** | | | | | | |
| **Baselga temporal beta sor dispersal logit** | Year1 | -0.0026 | 0.0028 | -0.92 | 0.357 | -0.0262 |
| **Baselga temporal beta sor thermal logit** | Year1 | -0.0150 | 0.0045 | -3.31 | <0.001 | -0.1503 |

**Table S6.** Estimated temporal slopes for guild-specific β-diversity metrics derived from mixed-effects and linear models. Slopes quantify rates of change in dominance-weighted spatial β-diversity ($\beta q_2$), Baselga spatial components (total dissimilarity βSOR, turnover βSIM, and nestedness βSNE), and temporal Baselga dissimilarity (βSOR; Δyear = 1). Spatial $\beta q_2$ slopes are reported both pooled across sampling gears and estimated separately by gear. Baselga spatial components are based on spatial comparisons among bays, whereas Baselga temporal βSOR represents year-to-year compositional change within bays. Models included year as a continuous predictor, with gear and season as fixed effects and bay as a random effect where applicable; temporal Baselga βSOR models were fitted on the logit scale. Positive slopes indicate increasing compositional differentiation or turnover through time, whereas negative slopes indicate homogenization. Reported values include slope estimates, standard errors, test statistics, and associated p-values.

| **response** | **guild** | **slope** | **se** | **df** | **stat** | **p** | **guild type** | **gear** | **component** |
|---|---|---|---|---|---|---|---|---|---|
| **Spatial $\beta q_2$ (pooled across gears)** | | | | | | | | | |
| beta q2 | High | -0.0013 | 0.0009 | inf | -1.48 | 0.139 | disp. | nan | nan |
| beta q2 | Intermediate | -0.0022 | 0.0009 | inf | -2.44 | 0.015 | disp. | nan | nan |
| beta q2 | Low | -0.0087 | 0.0009 | inf | -9.44 | <0.001 | disp. | nan | nan |
| beta q2 | Subtropical | 0.0018 | 0.0010 | inf | 1.83 | 0.067 | therm. | nan | nan |
| beta q2 | Temperate | -0.0028 | 0.0010 | inf | -2.85 | 0.004 | therm. | nan | nan |
| beta q2 | Tropical | -0.0050 | 0.0014 | inf | -3.49 | <0.001 | therm. | nan | nan |
| **Spatial $\beta q_2$ (by gear)** | | | | | | | | | |
| beta q2 | High | -0.0019 | 0.0012 | inf | -1.64 | 0.101 | disp. | BS | nan |
| beta q2 | Intermediate | -0.0006 | 0.0012 | inf | -0.49 | 0.625 | disp. | BS | nan |
| beta q2 | Low | -0.0030 | 0.0012 | inf | -2.53 | 0.011 | disp. | BS | nan |
| beta q2 | Subtropical | 0.0010 | 0.0015 | inf | 0.67 | 0.500 | therm. | BS | nan |
| beta q2 | Temperate | -0.0027 | 0.0015 | inf | -1.82 | 0.069 | therm. | BS | nan |
| beta q2 | Tropical | 0.0071 | 0.0030 | inf | 2.38 | 0.017 | therm. | BS | nan |
| beta q2 | High | -0.0006 | 0.0014 | inf | -0.41 | 0.681 | disp. | BT | nan |
| beta q2 | Intermediate | -0.0064 | 0.0014 | inf | -4.57 | <0.001 | disp. | BT | nan |
| beta q2 | Low | -0.0151 | 0.0014 | inf | -10.73 | <0.001 | disp. | BT | nan |

| | | | | | | | | | |
|---|---|---|---|---|---|---|---|---|---|
| beta q2 | Subtropical | 0.0024 | 0.0017 | inf | 1.40 | 0.163 | therm. | BT | nan |
| beta q2 | Temperate | -0.0065 | 0.0017 | inf | -3.86 | <0.001 | therm. | BT | nan |
| beta q2 | Tropical | -0.0092 | 0.0022 | inf | -4.20 | <0.001 | therm. | BT | nan |
| beta q2 | High | -0.0025 | 0.0020 | 1748.2 | -1.26 | 0.207 | disp. | GN | nan |
| beta q2 | Intermediate | 0.0024 | 0.0020 | 1748.2 | 1.18 | 0.240 | disp. | GN | nan |
| beta q2 | Low | -0.0084 | 0.0024 | 1748.2 | -3.49 | <0.001 | disp. | GN | nan |
| beta q2 | Subtropical | 0.0023 | 0.0018 | 1708.1 | 1.31 | 0.191 | therm. | GN | nan |
| beta q2 | Temperate | 0.0050 | 0.0018 | 1708.1 | 2.82 | 0.005 | therm. | GN | nan |
| beta q2 | Tropical | -0.0042 | 0.0022 | 1708.3 | -1.91 | 0.056 | therm. | GN | nan |
| **Spatial Baselga components** | | | | | | | | | |
| baselga spatial | High | -0.0004 | 0.0003 | 1211.0 | -1.26 | 0.206 | disp. | nan | total |
| baselga spatial | Intermediate | -0.0002 | 0.0003 | 1211.0 | -0.56 | 0.573 | disp. | nan | total |
| baselga spatial | Low | 0.0014 | 0.0003 | 1211.0 | 4.72 | <0.001 | disp. | nan | total |
| baselga spatial | High | -0.0002 | 0.0003 | 1180.0 | -0.59 | 0.558 | disp. | nan | turnover |
| baselga spatial | Intermediate | 0.0003 | 0.0003 | 1180.0 | 0.88 | 0.381 | disp. | nan | turnover |
| baselga spatial | Low | 0.0024 | 0.0003 | 1180.0 | 7.56 | <0.001 | disp. | nan | turnover |
| baselga spatial | High | -0.0002 | 0.0002 | 1180.0 | -1.19 | 0.234 | disp. | nan | nestedness |
| baselga spatial | Intermediate | -0.0004 | 0.0002 | 1180.0 | -2.39 | 0.017 | disp. | nan | nestedness |
| baselga spatial | Low | -0.0007 | 0.0002 | 1180.0 | -3.60 | <0.001 | disp. | nan | nestedness |
| baselga spatial | Subtropical | 0.0005 | 0.0004 | 958.0 | 1.20 | 0.229 | therm. | nan | total |
| baselga spatial | Temperate | 0.0000 | 0.0004 | 958.0 | 0.10 | 0.921 | therm. | nan | total |

| | | | | | | | | | |
|---|---|---|---|---|---|---|---|---|---|
| baselga spatial | Tropical | -0.0052 | 0.0007 | 958.0 | -7.63 | <0.001 | therm. | nan | total |
| baselga spatial | Subtropical | 0.0003 | 0.0004 | 872.0 | 0.72 | 0.471 | therm. | nan | turnover |
| baselga spatial | Temperate | 0.0008 | 0.0004 | 872.0 | 2.16 | 0.031 | therm. | nan | turnover |
| baselga spatial | Tropical | -0.0039 | 0.0013 | 872.0 | -3.12 | 0.002 | therm. | nan | turnover |
| baselga spatial | Subtropical | 0.0002 | 0.0002 | 872.0 | 1.07 | 0.285 | therm. | nan | nestedness |
| baselga spatial | Temperate | -0.0007 | 0.0002 | 872.0 | -3.73 | <0.001 | therm. | nan | nestedness |
| baselga spatial | Tropical | 0.0009 | 0.0006 | 872.0 | 1.41 | 0.160 | therm. | nan | nestedness |
| **Temporal Baselga βSOR (logit)** | | | | | | | | | |
| baselga temporal | High | 0.0016 | 0.0049 | inf | 0.33 | 0.742 | disp. | nan | nan |
| baselga temporal | Intermediate | 0.0059 | 0.0049 | inf | 1.20 | 0.229 | disp. | nan | nan |
| baselga temporal | Low | -0.0156 | 0.0050 | inf | -3.16 | 0.002 | disp. | nan | nan |
| baselga temporal | Subtropical | 0.0013 | 0.0072 | inf | 0.18 | 0.856 | therm. | nan | nan |
| baselga temporal | Temperate | 0.0040 | 0.0072 | inf | 0.56 | 0.577 | therm. | nan | nan |
| baselga temporal | Tropical | -0.0770 | 0.0096 | inf | -8.04 | <0.001 | therm. | nan | nan |

**Table S7**. Pairwise contrasts testing differences in temporal slopes of guild-specific β-diversity metrics. Comparisons evaluate whether rates of change in dominance-weighted spatial β-diversity ($\beta q_2$), Baselga spatial components (βSOR, βSIM, βSNE), and temporal Baselga dissimilarity (βSOR; Δyear = 1) differ among guild levels within dispersal and thermal classifications. Slope differences (Δslope) were estimated using post hoc trend contrasts derived from fitted models, with p-values adjusted for multiple comparisons using the Holm method. Positive Δslope values indicate faster increases (or weaker declines) in β-diversity for the first-listed guild relative to the second-listed guild, whereas negative values indicate slower increases or stronger homogenization. Models correspond to those summarized in Table S7.

| model | contrast | diff slope | se | df | stat | p | guild type | gear | component |
|---|---|---|---|---|---|---|---|---|---|
| Spatial $\beta q_2$ (pooled across gears) | | | | | | | | | |
| beta q2 | High - Intermediate | 0.0009 | 0.0013 | inf | 0.68 | 0.498 | dispersal | nan | nan |
| beta q2 | High - Low | 0.0074 | 0.0013 | inf | 5.75 | <0.001 | dispersal | nan | nan |
| beta q2 | Intermediate - Low | 0.0065 | 0.0013 | inf | 5.08 | <0.001 | dispersal | nan | nan |
| beta q2 | Subtropical - Temperate | 0.0047 | 0.0014 | inf | 3.32 | 0.002 | thermal | nan | nan |
| beta q2 | Subtropical - Tropical | 0.0068 | 0.0017 | inf | 3.91 | <0.001 | thermal | nan | nan |
| beta q2 | Temperate - Tropical | 0.0021 | 0.0017 | inf | 1.23 | 0.219 | thermal | nan | nan |
| Spatial $\beta q_2$ (by gear) | | | | | | | | | |
| beta q2 | High - Intermediate | -0.0014 | 0.0017 | inf | -0.81 | 0.831 | dispersal | BS | nan |
| beta q2 | High - Low | 0.0011 | 0.0017 | inf | 0.64 | 0.831 | dispersal | BS | nan |
| beta q2 | Intermediate - Low | 0.0024 | 0.0017 | inf | 1.45 | 0.441 | dispersal | BS | nan |
| beta q2 | Subtropical - Temperate | 0.0037 | 0.0021 | inf | 1.76 | 0.135 | thermal | BS | nan |
| beta q2 | Subtropical - Tropical | -0.0061 | 0.0033 | inf | -1.83 | 0.135 | thermal | BS | nan |

| | | | | | | | | | |
|---|---|---|---|---|---|---|---|---|---|
| beta q2 | Temperate - Tropical | -0.0098 | 0.0033 | inf | -2.95 | 0.010 | thermal | BS | nan |
| beta q2 | High - Intermediate | 0.0058 | 0.0020 | inf | 2.94 | 0.003 | dispersal | BT | nan |
| beta q2 | High - Low | 0.0145 | 0.0020 | inf | 7.30 | <0.001 | dispersal | BT | nan |
| beta q2 | Intermediate - Low | 0.0087 | 0.0020 | inf | 4.36 | <0.001 | dispersal | BT | nan |
| beta q2 | Subtropical - Temperate | 0.0088 | 0.0024 | inf | 3.72 | <0.001 | thermal | BT | nan |
| beta q2 | Subtropical - Tropical | 0.0115 | 0.0028 | inf | 4.18 | <0.001 | thermal | BT | nan |
| beta q2 | Temperate - Tropical | 0.0027 | 0.0028 | inf | 0.97 | 0.332 | thermal | BT | nan |
| beta q2 | High - Intermediate | -0.0049 | 0.0028 | 1748.01544875716 | -1.72 | 0.125 | dispersal | GN | nan |
| beta q2 | High - Low | 0.0058 | 0.0031 | 1748.1812344548 | 1.86 | 0.125 | dispersal | GN | nan |
| beta q2 | Intermediate - Low | 0.0107 | 0.0031 | 1748.18123308872 | 3.43 | 0.002 | dispersal | GN | nan |
| beta q2 | Subtropical - Temperate | -0.0027 | 0.0025 | 1707.99781722813 | -1.07 | 0.284 | thermal | GN | nan |
| beta q2 | Subtropical - Tropical | 0.0065 | 0.0028 | 1708.08478327351 | 2.31 | 0.042 | thermal | GN | nan |
| beta q2 | Temperate - Tropical | 0.0092 | 0.0028 | 1708.08470883112 | 3.25 | 0.003 | thermal | GN | nan |
| Spatial Baselga components | | | | | | | | | |
| baselga spatial | High - Intermediate | -0.0002 | 0.0004 | 1211.0 | -0.50 | 0.621 | dispersal | nan | total |
| baselga spatial | High - Low | -0.0018 | 0.0004 | 1211.0 | -4.31 | <0.001 | dispersal | nan | total |

| | | | | | | | | | |
|---|---|---|---|---|---|---|---|---|---|
| baselga spatial | Intermediate - Low | -0.0016 | 0.0004 | 1211.0 | -3.83 | <0.001 | dispersal | nan | total |
| baselga spatial | High - Intermediate | -0.0004 | 0.0004 | 1180.0 | -1.03 | 0.301 | dispersal | nan | turnover |
| baselga spatial | High - Low | -0.0026 | 0.0004 | 1180.0 | -6.03 | <0.001 | dispersal | nan | turnover |
| baselga spatial | Intermediate - Low | -0.0022 | 0.0004 | 1180.0 | -5.05 | <0.001 | dispersal | nan | turnover |
| baselga spatial | High - Intermediate | 0.0002 | 0.0003 | 1180.0 | 0.85 | 0.556 | dispersal | nan | nestedness |
| baselga spatial | High - Low | 0.0005 | 0.0003 | 1180.0 | 1.89 | 0.178 | dispersal | nan | nestedness |
| baselga spatial | Intermediate - Low | 0.0003 | 0.0003 | 1180.0 | 1.09 | 0.556 | dispersal | nan | nestedness |
| baselga spatial | Subtropical - Temperate | 0.0004 | 0.0005 | 958.0 | 0.78 | 0.435 | thermal | nan | total |
| baselga spatial | Subtropical - Tropical | 0.0057 | 0.0008 | 958.0 | 7.23 | <0.001 | thermal | nan | total |
| baselga spatial | Temperate - Tropical | 0.0053 | 0.0008 | 958.0 | 6.69 | <0.001 | thermal | nan | total |
| baselga spatial | Subtropical - Temperate | -0.0005 | 0.0005 | 872.0 | -1.02 | 0.309 | thermal | nan | turnover |
| baselga spatial | Subtropical - Tropical | 0.0042 | 0.0013 | 872.0 | 3.20 | 0.003 | thermal | nan | turnover |
| baselga spatial | Temperate - Tropical | 0.0047 | 0.0013 | 872.0 | 3.60 | <0.001 | thermal | nan | turnover |

| | | | | | | | | | |
|---|---|---|---|---|---|---|---|---|---|
| baselga spatial | Subtropical - Temperate | 0.0009 | 0.0003 | 872.0 | 3.40 | 0.002 | thermal | nan | nestedness |
| baselga spatial | Subtropical - Tropical | -0.0007 | 0.0007 | 872.0 | -1.05 | 0.294 | thermal | nan | nestedness |
| baselga spatial | Temperate - Tropical | -0.0016 | 0.0007 | 872.0 | -2.40 | 0.033 | thermal | nan | nestedness |
| Temporal Baselga βSOR (logit) | | | | | | | | | |
| baselga temporal | High - Intermediate | -0.0043 | 0.0069 | inf | -0.62 | 0.537 | dispersal | nan | nan |
| baselga temporal | High - Low | 0.0173 | 0.0070 | inf | 2.48 | 0.027 | dispersal | nan | nan |
| baselga temporal | Intermediate - Low | 0.0215 | 0.0070 | inf | 3.09 | 0.006 | dispersal | nan | nan |
| baselga temporal | Subtropical - Temperate | -0.0027 | 0.0102 | inf | -0.27 | 0.791 | thermal | nan | nan |
| baselga temporal | Subtropical - Tropical | 0.0783 | 0.0120 | inf | 6.52 | <0.001 | thermal | nan | nan |
| baselga temporal | Temperate - Tropical | 0.0810 | 0.0120 | inf | 6.75 | <0.001 | thermal | nan | nan |

**Table S8.** Statistical support for temporal trends and derivative-based diagnostics (α-, β-, and γ-diversity).

Generalized additive models (GAMs) were fit separately to each gear × season time series using a population-smooth formulation that included s(YEAR) and a bay random effect s(MAJOR_AREA, bs = "re") (used in all fits here). Reported statistics include the estimated degrees of freedom (edf) and p-value for the temporal smooth (p_smooth), along with Benjamini–Hochberg FDR-adjusted p-values computed within each metric (p_smooth_fdr). Derivative diagnostics summarize (i) confidence-interval–supported sign changes in the first derivative (sign_change_years) and (ii) peaks in the absolute second derivative (accel_peak_years; acceleration/curvature extrema). For GN, only SPR and FALL were analyzed.

| metric | gear | Season | n_obs | edf | p_smooth | p_smooth_fdr | n | sign_change_years | n | accel_peak_years |
|---|---|---|---|---|---|---|---|---|---|---|
| alpha | BS | SPR | 340 | 7.928 | <1e-16 | <1e-16 | 0 | NA | 2 | 2022;2016 |
| alpha | BS | SUM | 340 | 6.852 | <1e-16 | <1e-16 | 0 | NA | 2 | 2020;1986 |
| alpha | BS | FALL | 340 | 9.205 | <1e-16 | <1e-16 | 2 | 2012;2016 | 2 | 1983;2018 |
| alpha | BS | WIN | 340 | 7.468 | <1e-16 | <1e-16 | 0 | NA | 2 | 2020;1989 |
| alpha | BT | SPR | 340 | 5.673 | 1.49e-05 | 1.86e-05 | 1 | 2019 | 1 | 2021 |
| alpha | BT | SUM | 340 | 6.652 | <1e-16 | <1e-16 | 1 | 2015 | 2 | 2024;2018 |
| alpha | BT | FALL | 340 | 7.241 | <1e-16 | <1e-16 | 2 | 1996;2018 | 2 | 2020;1990 |
| alpha | BT | WIN | 340 | 5.006 | 0.001239 | 0.001549 | 1 | 2018 | 2 | 2019;2010 |
| alpha | GN | SPR | 324 | 5.981 | <1e-16 | <1e-16 | 1 | 1994 | 2 | 1983;1988 |
| alpha | GN | FALL | 332 | 5.657 | <1e-16 | <1e-16 | 1 | 1996 | 2 | 1984;2000 |
| beta | BS | SPR | 340 | 0.776 | 0.034821 | 0.084014 | 0 | NA | 0 | NA |
| beta | BS | SUM | 340 | 0.428 | 0.212618 | 0.319401 | 0 | NA | 0 | NA |
| beta | BS | FALL | 340 | 0.000227 | 0.686248 | 0.814314 | 0 | NA | 0 | NA |
| beta | BS | WIN | 340 | 2.744 | 0.116534 | 0.194223 | 0 | NA | 1 | 1985 |
| beta | BT | SPR | 340 | 0.002345 | 0.319401 | 0.399252 | 0 | NA | 0 | NA |
| beta | BT | SUM | 340 | 0.920 | 0.008861 | 0.044306 | 0 | NA | 2 | 1992;1986 |
| beta | BT | FALL | 340 | 0.814 | 0.053977 | 0.089962 | 0 | NA | 0 | NA |
| beta | BT | WIN | 340 | 0.935 | 0.001815 | 0.018152 | 0 | NA | 2 | 2020;2013 |
| beta | GN | SPR | 324 | 0.775 | 0.042007 | 0.084014 | 0 | NA | 0 | NA |
| beta | GN | FALL | 332 | 2.49e-05 | 0.972072 | 0.972072 | 0 | NA | 0 | NA |
| gamma | BS | SPR | 340 | 7.156 | <1e-16 | <1e-16 | 2 | 2010;2016 | 2 | 2021;1984 |
| gamma | BS | SUM | 340 | 7.835 | <1e-16 | <1e-16 | 2 | 2011;2014 | 2 | 1984;2023 |
| gamma | BS | FALL | 340 | 8.465 | <1e-16 | <1e-16 | 4 | 2000;2003;2010;2015 | 2 | 1984;2021 |
| gamma | BS | WIN | 340 | 8.284 | <1e-16 | <1e-16 | 4 | 1999;2004;2013;2016 | 2 | 2021;1983 |
| gamma | BT | SPR | 340 | 5.961 | <1e-16 | <1e-16 | 2 | 1999;2020 | 2 | 2022;2010 |
| gamma | BT | SUM | 340 | 8.347 | <1e-16 | <1e-16 | 4 | 1996;2003;2009;2016 | 2 | 1983;2021 |
| gamma | BT | FALL | 340 | 6.982 | <1e-16 | <1e-16 | 2 | 1996;2019 | 2 | 2024;1984 |
| gamma | BT | WIN | 340 | 8.589 | <1e-16 | <1e-16 | 6 | 1990;1996;2002;2007;2014;2020 | 2 | 2023;1983 |
| gamma | GN | SPR | 324 | 6.211 | <1e-16 | <1e-16 | 2 | 1992;2010 | 2 | 1983;2002 |
| gamma | GN | FALL | 332 | 7.321 | <1e-16 | <1e-16 | 5 | 1988;1994;2009;2014;2020 | 2 | 1982;2022 |
